# Gut hormone signaling drives sex differences in metabolism and behavior

**DOI:** 10.1101/2025.09.10.675306

**Authors:** Olga Kubrak, Alina Malita, Nadja Ahrentløv, Stanislav Nagy, Michael J. Texada, Kim Rewitz

**Author notes:** Equal contribution (Olga Kubrak, Alina Malita).

## Abstract

Males and females have different physiological and reproductive demands, and consequently exhibit widespread differences in metabolism and behavior. One of the most consistent differences across animals is that females store more body fat than males, a metabolic trait conserved from flies to humans. Given the central role of gut hormones in energy balance, we asked whether gut endocrine signaling underlies these sex differences. We therefore performed a multidimensional screen of enteroendocrine cell (EEC)-derived signaling across a broad panel of metabolic and behavioral traits in male and female *Drosophila*. This screen uncovered extensive sex-biased roles for EEC-derived signals – many of which are conserved in mammals – in energy storage, stress resistance, feeding, and sleep. We found that EEC-derived amidated peptide hormones sustain female-typical states, including elevated fat reserves, enhanced stress resilience, and protein-biased food choice. In contrast, the non-amidated peptide Allatostatin C (AstC) promoted male-like traits by stimulating energy mobilization, thereby antagonizing amidated-peptide function. Female guts contained more AstC-positive EECs. Disruption of peptide amidation by eliminating peptidylglycine α-hydroxylating monooxygenase – the enzyme required for maturation of most gut peptide hormones – abolished female-typical physiology and behavior, shifting females toward a male-like state. Among individual amidated peptides, Diuretic hormone 31 (DH31) and Neuropeptide F (NPF) emerged as key mediators of female physiology. These findings establish gut hormone signaling as a determinant of sex-specific metabolic and behavioral states.

## Introduction

Males and females differ in many aspects of their physiology, from hormonal regulation to tissue composition. One of the most consistent differences across species is in body-fat storage. In humans, women on average carry a markedly higher proportion of body fat than men, typically 25–35% of body mass in women compared to 10–20% in men^1-3^. Similar patterns are seen in the fruit fly *Drosophila*, in which adult females accumulate significantly higher lipid reserves than males^4-6^. Sex-specific differences in energy storage suggest that these reserves may contribute to distinct physiological and behavioral outcomes in males and females. However, the mechanisms that establish and maintain these differences remain poorly understood, representing an underexplored aspect of metabolic research. To begin addressing this gap, it is essential to decipher not only differences in energy storage but also sex-specific behaviors and the hormonal systems that drive divergent male and female physiological states.

The *Drosophila* model enables systematic dissection of the endocrine signals that shape sex-specific physiology, making it possible to uncover fundamental principles of sex-biased metabolic regulation. In addition to clear sex differences in metabolic traits, similar to those observed in mammals, *Drosophila* exhibits striking sex-dependent behavioral variation. For example, females feed more often and for longer, show a stronger appetite for protein-rich food, and sleep less than males^7-9^. These behavioral patterns align with females’ higher reproductive energy demands and greater nutrient investment in egg production. In flies, as in mammals, metabolism and energy balance are regulated by two central opposing hormonal systems, with an insulin-like pathway and a glucagon-like pathway that together control the balance between nutrient storage and mobilization^10,11^. *Drosophila* insulin-like peptides (Ilps) are evolutionarily conserved and signal through an insulin receptor that can also be activated by human insulin^12^. Insulin promotes anabolic processes such as nutrient storage and growth, while adipokinetic hormone (AKH) functions in a glucagon-like manner to mobilize energy and favor sugar utilization. These opposing pathways contribute to sex-specific physiology, with females generally showing higher insulin activity and lower AKH activity than males, consistent with their greater fat storage^6,13^.

Despite these robust differences, the origins of sex-biased metabolic and behavioral states remain poorly understood. They may arise from intrinsic properties of peripheral tissues, systemic endocrine regulation, or inter-organ communication. The gut has recently emerged as a key site of sex differences, with sex-biased gene expression, stem-cell proliferation, and microbial interactions reported^14-16^. Sex differences in gut-derived endocrine signaling have been described, and many gut-derived hormones in *Drosophila* regulate feeding and metabolism by converging on insulin and AKH pathways, perhaps contributing to the sex-dependent differences in activity of these systems^17^. This rich endocrine signaling positions the gut as a potential driver of systemic sex differences in metabolism and behavior.

Enteroendocrine cells (EECs) are central to this process as the principal endocrine cells of the gut. EECs of molecularly distinct classes are distributed throughout the gut epithelium in both flies and mammals, from which they secrete, into the nearby cellular milieu and the systemic circulation, diverse peptide hormones and other signals that regulate appetite, metabolism, and stress resistance^17,18^. In *Drosophila*, EECs release neuropeptide F (NPF), allatostatin C (AstC), tachykinin (Tk), and other conserved hormones whose mammalian counterparts – peptide YY (PYY), somatostatin, and substance P – are also expressed by intestinal EECs, underscoring the conservation of gut endocrine regulation across species. In flies, NPF and Tk regulate feeding decisions and food choice between sugar and protein by modulating AKH^8,19^. Gut-derived NPF acts in a sex-specific manner in females, suppressing sugar intake after mating by inhibiting AKH activity^8^. This shift favors nutrient storage and biases appetite toward protein-rich yeast food, which is critical for reproduction. AstC also acts in a sex-biased manner, promoting energy mobilization specifically in females by stimulating AKH release^20^, thereby opposing the storage-promoting actions of NPF.

Although several gut hormones have been implicated in traits that differ between the sexes, whether the EEC population itself is sexually dimorphic, and the overall extent to which gut hormone signaling contributes to male–female physiological differences, remain poorly defined and have never been systematically studied in any organism. Here, we fill this gap by systematically dissecting anatomical sex differences in EECs and the contribution of EEC-derived signals to sex-specific physiology and behavior in the fly. Our findings establish the gut endocrine system as a major determinant of sex-specific physiological states.

## Results

### Sexual dimorphism in adult gut-endocrine anatomy

The gastrointestinal tract plays a central role in coordinating whole-body metabolism and physiology through the release of hormones and other signaling molecules^21-25^. Enteroendocrine cells (EECs), a relatively small but highly diverse population of intestinal secretory cells, act as nutrient sensors and orchestrators of interorgan communication. Despite the importance of these cells, the functional diversity among the signals they produce remains incompletely understood. Moreover, increasing evidence suggests that gut physiology differs between the sexes, potentially contributing to sex-specific mechanisms of maintaining energy balance and regulating behavior^16,26,27^. However, whether and how EEC-derived signals contribute to sex-linked traits remains largely unknown, and systematic studies addressing the impact of sex on these signals’ functions are lacking.

To address this question, we used as our model the adult *Drosophila* midgut. This system allows precise genetic manipulation and *in-vivo* monitoring of EEC populations, making it a uniquely useful target for systematic functional analysis. The great majority of adult fly EECs fall into one of two very broad subtypes, with cells of one type producing the peptide AstC, a conserved Somatostatin-like factor, and cells of the other producing Tk, homologous to mammalian tachykinin-like factors such as Substance P. Within these broad categories, EECs can be divided into functionally and anatomically distinct classes, each defined by stereotyped distribution and co-expression of two to five peptide hormones^17,28,29^.

The AstC-positive population is concentrated in the middle and posterior midgut, and within this group are classes that also produce AstA and other peptides^17^. AstC itself contributes to systemic energy storage and stress resistance^17,20^. The conserved mammalian homolog of AstA is galanin, which is expressed in enteric neurons and regulates the release of the incretin hormones glucagon-like peptide-1 (GLP-1) and glucose-dependent insulinotropic polypeptide (GIP), key enteroendocrine-cell-derived signals that regulate appetite^30,31^. The Tk-positive EECs are more evenly distributed along the gut, with enrichment in the middle midgut and depletion in a posterior-mid segment (R4), and this group comprises subsets that also produce the peptide factors Diuretic hormone 31 (Dh31) and NPF. The mammalian homolog of *Drosophila* Dh31 is calcitonin gene-related peptide (CGRP), which is expressed in enteric neurons^32^, whereas the mammalian homolog of NPF is PYY, an abundantly expressed product of EECs of the human intestine^8,33^. The Tk, Dh31, and NPF hormones derived from Tk-positive EECs regulate feeding, gut motility, and systemic nutrient signaling^8,17,19,34-36^.

However, it is worth noting that existing general anatomical and molecular descriptions of adult midgut EECs, such as single-cell transcriptomic analyses^28,29,37^, are derived from work with female tissues alone. We wished to compare the endocrine-related anatomical properties of male and female midguts, and therefore we imaged multiple fixed and stained tissues in their entirety at high resolution. This allowed comprehensive mapping of all gut cells, including EEC populations genetically defined and labeled by GFP expressed under EEC-specific drivers. We used *voilà-GAL4* (*voilà*>), an insertion into the pan-enteroendocrine-cell marker gene *prospero* that therefore labels all EECs^20^, to drive GFP expression and thereby obtain a global view of the adult gut endocrine system in males and females. To assess whether sex-dependent differences exist within the AstC-positive or Tk-positive cell classes, we used *AstC::T2A::GAL4* (*AstC*>) and *Tk::T2A::GAL4* (*Tk*>) knock-in drivers inserted at the endogenous loci, ensuring faithful labeling of the corresponding peptide-expressing cells^19,20^.

In our analysis, male midguts were consistently shorter and contained correspondingly fewer total cells compared to age-matched mated female midguts, as previously reported (Fig. 1a and S1a,b,c). Part of this difference reflects overall body-size scaling, since males are generally smaller than females. However, female midguts also undergo sex-specific growth and remodeling, particularly after mating, which increases gut length and cell number^38,39^. Thus, the observed differences represent both scaling with body size and intrinsic sex- and mating-associated variation in gut anatomy.

**Figure 1.**
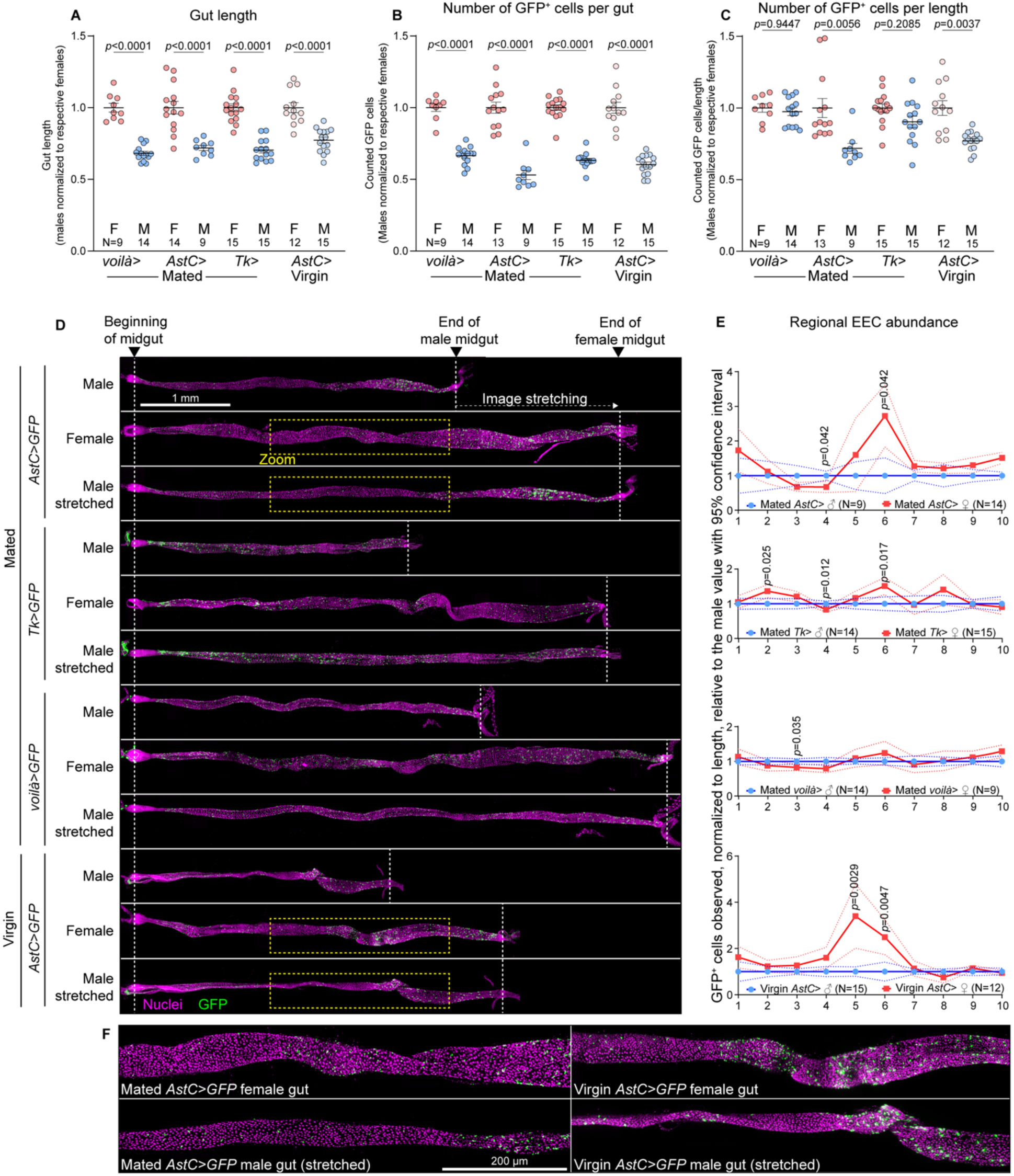
Sexually dimorphic organization of the adult gut endocrine system. (a) The length of the midgut in males (M) and females (F) expressing GFP in defined enteroendocrine-cell (EEC) populations using *voilà-GAL4*, *AstC::T2A::GAL4* (*AstC*>), or *Tk::T2A::GAL4* (*Tk*>) drivers, or *AstC*> in virgin females and age-matched males. Values for each sex and genotype are normalized to the female average. (b) The number of GFP-positive EECs per gut, normalized as in (a). (c) The number of GFP-positive EECs per unit length, removing the effect of increased female gut length. Values were further normalized within each genotype to the female reference. (d) Representative midgut images showing the spatial distribution of GFP-positive EECs (green; nuclei, red) in males and females. (e) Quantification of spatial distribution across ten equal-length regions of the midgut. Females displayed a two- to threefold enrichment of AstC-positive cells in the middle midgut, while Tk-positive EECs showed little sex difference. (f) High-magnification images confirming enrichment of AstC-positive cells in female midguts, observed in both mated and virgin females. Scale bar, 200 microns. Statistics: A-D: Welch’s ANOVA with Dunnett’s T3 multiple comparisons; F-I: Significance of differences between male and female for each decile was calculated using unpaired two-sided *t* tests with Holm-Šidák correction for multiple tests. Data in (a–c, e) are mean ± s.e.m. In (e), *p* values greater than 0.05 are not shown.

Consistent with their reduced length, male midguts exhibited fewer EECs than those of females in the pan-EEC *voilà>GFP* analysis (Fig. 1b). This difference was also observed when the AstC- and Tk-positive populations were assessed (Fig. 1b). The sum of AstC-positive and Tk-positive cells was roughly comparable to the *voilà*-positive population, suggesting that together they capture nearly all EECs (Fig. S1d). When normalized for differences in gut length between sexes, there was little, if any, difference in the number of *prospero*-expressing EECs (the pan-EEC population), and Tk-positive EECs were also similarly numerous in males and females after normalization for midgut length (Fig. 1c).

However, mated female guts contained significantly more AstC-positive cells than would be expected if these numbers simply scaled with gut length (Fig. 1c and S1e), suggesting differences in cellular proliferation, lifespan, or loss between the sexes. The female midgut undergoes growth and remodeling after mating as a means to support the nutritional needs of egg production^38,39^, and the excess AstC cells we observed in mated female guts might have arisen as part of this remodeling. We therefore analyzed virgin females alongside age-matched males and found a similar female-specific enrichment in AstC-positive cells in these unmated animals, indicating that this difference represents an inherent feature of the female gut rather than an adaptive response to mating (Fig. 1c and S1e). This suggests that sexual dimorphism in the gut endocrine system is cell-type-specific, with AstC-positive EECs being disproportionately enriched in females.

We next asked whether the enrichment of AstC-positive cells in the female gut was localized to a specific region of the midgut. We computationally divided each gut into ten equal-length regions (deciles) and counted the GFP-positive cells in each. Since the number of EECs per unit gut length appears equivalent between males and females (*voilà>GFP*), the greater length of female guts means that each region spans more tissue and therefore contains more EECs than the corresponding male region (Fig. S1d). To identify deviations from this equal-density expectation, we first estimated how many EECs a female gut would be expected to have in each region if the only difference was length, using the male data as a baseline. We then compared the observed number of EECs in each female region to the expected value. A ratio greater than 1 indicates more cells than expected in females (overabundance), while a ratio less than 1 indicates fewer cells than expected (underabundance). This analysis showed only minor differences between male and female EEC density along the length of the midgut for the total EEC population marked by *voilà*> (Fig. 1d,e). The Tk-positive EEC population also exhibited only relatively small regional differences between male and female guts. However, this analysis revealed a striking enrichment of AstC-positive EECs in the middle midgut region of females, where two to three times as many AstC-positive cells were noted than would have been expected based on extrapolation from the corresponding male regions (Fig. 1d-f). A similar enrichment was also apparent in virgin females, indicating that this feature is not induced by mating but represents an intrinsic, sex-specific characteristic of midgut endocrine anatomy.

The region-restricted variation we identify here provides a likely cellular basis for sex-biased roles of AstC signaling. In our previous work, we showed that NPF/NPFR signaling regulates female physiology and behavior through AstC-positive EECs in this region, with manipulations in males having no detectable effect^20^. These findings suggest that the female-specific enrichment of AstC EECs may constitute a regulatory module with NPF that is functionally important in females^8^. Consistent with this, published single-cell transcriptomic data^28^, derived from females, describe a molecularly distinct AstC-positive population present only this region of the midgut that is notable for its strong expression of NPFR. Taken together, our anatomical and functional data, in combination with transcriptomic insights, suggest that AstC cells in the middle midgut region likely represent a specialized female-enriched EEC population that contributes to sex-biased regulation of energy balance and behavior. The broad anatomical dimorphism we identify here provides a plausible cellular basis for sex-biased physiological roles of AstC signaling, in conjunction with sex-linked differences in AstC-receptive target tissues.

### A systematic RNAi screen uncovers sex-specific roles of enteroendocrine cell-derived signals

We observed above that sex-dependent differences in the gut endocrine system indeed exist, with anatomical and cellular specificity. We next asked how EEC-derived signals, originating from cells that may differ in abundance or physiological properties between sexes, contribute to systemic phenotypes. We performed a multidimensional RNAi screen targeting genes expressed in adult EECs to characterize these genes’ functions, including any sex-specific aspects (Fig. 2a). Genes were selected based on a curated target list integrating multiple criteria, including enrichment in the adult midgut (FlyAtlas2)^40^, expression in EECs in particular (FlyGutSeq)^37^, predicted extracellular localization (GO terms and protein-sequence features), and prior evidence of involvement in hormonal signaling or regulation by starvation (from publicly available RNAseq datasets)^41^. This list included genes encoding secreted factors (*e.g.*, protein and peptide hormones), enzymes (such as those involved in peptide processing or biogenic-amine biosynthesis), receptors (including orphan and hormone-binding GPCRs), and transporters for metabolites and nutrients including sugars and amino acids.

**Figure 2.**
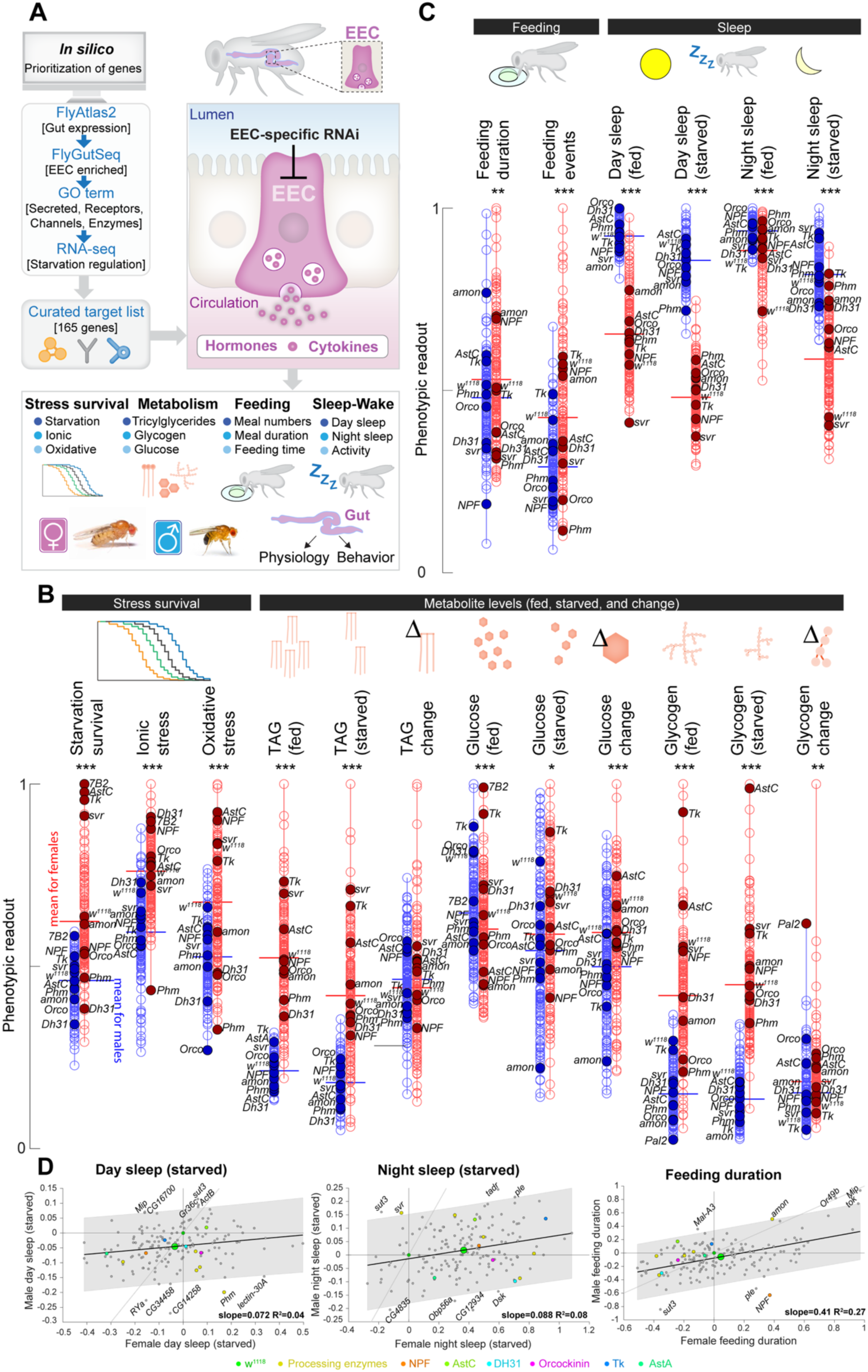
A systematic RNAi screen reveals sex-specific roles of enteroendocrine cell-derived signals in systemic physiology and behavior. (a) Schematic of the screening pipeline. Schematic of the screening pipeline. Screened genes were selected based upon multiple criteria from a candidate pool comprising secreted factors, relevant enzymes, receptors, transporters, and GPCRs. Genes were targeted using RNAi in adult EECs, and a diverse array of phenotypes was assessed. (b) Results from the RNAi screen across stress survival and metabolic assays. Each point represents the mean phenotype for an individual gene knockdown in males (blue) or females (red). For each assay, values are scaled so that the maximum observed value (male or female) is set to 1.0. Horizontal lines indicate the mean phenotype across all knockdowns for each sex. Stress assays include starvation (1% agar in water), ionic stress (4% NaCl in food), and oxidative stress (5% H_2_O_2_ in food). Metabolic assays include measurements of triacylglycerides (TAG), glycogen, and glucose in fed and starved animals as well as their utilization during starvation. Selected knockdowns are labeled. (c) Screen results for behavioral assays. Feeding was quantified as feeding duration and number of feeding events in the FLIC assay, and sleep-wake behavior was assessed under fed and starved conditions. Charts are plotted as in (b). (d) Comparative analysis of male and female responses. Plots of female phenotypic changes (*e.g.*, a 10% increase in fat or sleep amount, relative to the control = 0.1) with knockdown (X axis) versus male changes (Y axis). Each knockdown genotype is represented by a point, and a trend line with 95% confidence intervals is shown. The control genotype falls at position (0, 0) by definition, and the mean of all genotypes is shown as a larger green dot. Points along the diagonal represent knockdowns with similar effects in both sexes. Selected knockdowns are labeled. (d) Distance from the diagonal indicates the degree of sex bias in effect size. Points falling into opposite quadrants, where the effect is positive in one sex but negative in the other, represent strong discordance, with the same knockdown producing opposing phenotypes in males and females.

To target gene knockdown to adult EECs in males and females, we expressed RNAi constructs using the *voilà>* driver, which is broadly expressed throughout the midgut (R1–R5) and widely used to target EECs^20,34,42-44^. To restrict expression to the adult stage, we combined it with *Tub-GAL80^TS^*, hereafter together referred to as *voilà>*^20^. However, EECs and neurons share many cellular features, among which is fate determination via Prospero. Therefore, this driver also exhibits some expression in the brain^20^, mostly but not entirely transient in younger animals, which may represent a potential limitation when interpreting certain systemic phenotypes. All animals were reared at 18 °C on standard diet to inhibit GAL4 activity until eclosion, after which they were transferred to an adult-optimized diet and shifted to 29 °C to permit GAL4 activity. Using this system we silenced 165 genes in the EECs of adult males and females and, after ∼5 days of knockdown induction, we measured phenotypes across multiple physiological domains. Our readouts were (1) survival under stress conditions – starvation (1% agar in water), ionic stress (4% NaCl in adult diet), and oxidative stress (5% H_2_O_2_ in adult food); (2) energy metabolism, including levels of triacylglycerides (TAG), glycogen, and glucose in fed and starved states, as well as changes in these metabolites during starvation, which we used as independent measures of energy mobilization; and (3) behavior, including feeding with sugar-only liquid food (meal number and duration) and sleep-wake cycles (day/night sleep and activity, in fed and starved conditions, which generally induce locomotor hyperactivity and sleep suppression). Sex differences in each of these traits were assessed. In total, we analyzed phenotypes in approximately 90,000 animals across all assays and conditions, including both sexes and controls (Table S1).

### EEC-derived factors differentially regulate energy storage, stress resistance and behavior in males and females

The global distribution of phenotypes across all assays highlighted both sex differences in traits and how individual EEC-derived signals influence metabolism, stress resistance, feeding, and sleep in males and females (Fig. 2b). While some EEC gene knockdowns produced sex-specific effects, others resulted in phenotypic effects shared between sexes.

For phenotypes linked to metabolic and nutrient status, knockdown of *AstC* in the EECs markedly increased starvation resistance in females, whereas males expressing *AstC* knockdown remained similar to controls, indicating a female-specific role for this signaling system in nutrient allocation (Fig. 2b). Consistent with this, *AstC* was also among the strongest hits for glycogen utilization during starvation in females, and EEC-targeted loss of this hormone promoted a female-specific increase in TAG levels in the fed state.

These results are consistent with previous evidence that gut AstC is required to mobilize glycogen and TAG in females, but not in males, and that it influences starvation resistance only in females^20^. In contrast, EEC knockdown of *Tachykinin* (*Tk*) regulated starvation phenotypes in the same direction – towards prolonged survival – in both sexes. Consistent with these phenotypes, fed-state TAG levels in both were elevated upon *Tk* knockdown. The metabolic changes observed upon EEC-specific *Tk* knockdown indicate that loss of this gut hormone does not elicit sex-divergent metabolic effects.

The enzyme peptidylglycine-α-hydroxylating monooxygenase (encoded by *Phm*) emerged as a strikingly sex-biased factor in stress resistance. Phm catalyzes the initial step in peptide amidation, a modification required for the maturation, stability, and function of most known peptide hormones^45-47^, with the notable exception of AstC, which undergoes cyclization rather than amidation. Knockdown of *Phm* in the EECs markedly reduced starvation resistance in females but had little effect in males. Despite these divergent life-historical outcomes, changes in metabolite levels under starvation (reductions in TAG and glycogen) were largely similar in males and females (Fig. 2b). This suggests that both sexes mobilize nutrients in comparable ways during nutrient deprivation, while females achieve higher survival through greater baseline energy storage.

Baseline total glucose levels did not show strong sex differences, indicating that males and females maintain similar glycemic levels. Taken together, these results demonstrate that EEC-derived factors – including peptide hormones and peptide-processing enzymes – exert both shared and sex-specific influences across metabolic, stress, and behavioral traits, providing a first functional map of gut endocrine signaling in adult males and females.

In stress-resilience phenotypes not directly linked to nutrient status – ionic and oxidative stress resistance – EEC-targeted *Phm* knockdown again revealed striking sex-biased effects (Fig. 2b). RNAi-expressing females showed markedly reduced resistance to these stressors, ranking among the lowest performers, whereas males were indistinguishable from controls. These findings point to a disproportionately female-specific requirement for amidated gut peptides in maintaining resilience under ionic and oxidative stress. In addition, *Phm* knockdown significantly reduced feeding behavior in females while having minimal effects in males, further underscoring the trend of amidated gut peptides’ having stronger systemic influence on female physiology.

We also observed clear sex-specific effects in behavioral phenotypes relating to feeding and sleep (Fig. 2c). Females normally display higher feeding activity than males, and our data confirm that females engage in more feeding events overall. This elevated female feeding was reduced by EEC *Phm* knockdown, while males were unaffected, indicating that amidated EEC-derived peptide hormones collectively are required to sustain higher female feeding. In addition, *Phm* knockdown in EECs produced distinct sleep phenotypes between sexes. EEC-specific loss of Phm increased sleep during starvation in females while reducing it in males during starvation, demonstrating that loss of gut-derived amidated peptides can elicit opposing effects between sexes.

These findings illustrate that in some cases the same perturbation can drive opposite outcomes in males and females, a discordant pattern. To capture these relationships systematically for single phenotypes, we plotted the effect of each gene knockdown in females (X axis) against its effect in males (Y axis) (Fig. 2d and S2a). A linear fit with 95% confidence interval is shown. Points along the diagonal represent knockdowns with similar effects in both sexes, and increasing distance from the diagonal indicates stronger sex bias. By contrast, points that fall into opposite quadrants (positive in females but negative in males, or vice versa) denote discordance, where the same knockdown produces opposing phenotypes in the two sexes. Across most metabolic traits, male and female responses were generally proportional or exhibited sex-biased effect sizes, but not effect direction, indicating that many EEC-derived signals regulate nutrient storage and mobilization in a common manner, while certain pathways and signals disproportionately influence phenotypes in one sex (Fig. S2a). Discordant effects, where loss of an EEC factor produced opposing phenotypes in males and females, were rare among stress and metabolic traits but were more common in behavioral phenotypes. Elimination of amidated peptides through *Phm* knockdown provided a clear example, with increased daytime sleep in females but reduced daytime sleep in males under starvation (Fig. 2d). Conversely, knockdown of *silver* (*svr*), which encodes a carboxypeptidase required for processing most Phm-dependent peptides as well as some non-amidated peptides such as AstC, generated discordant effects on nighttime sleep in starved animals – males exhibited increased sleep, whereas females showed reduced sleep. Together, this analysis suggests that while many EEC signals exert concordant, sex-biased, or sex-specific systemic effects in males and females, peptide-processing enzymes such as Phm and Svr generate pronounced sex discordances in sleep regulation. These opposite responses indicate that gut-derived endocrine signals engage sex-specific regulatory networks, which can drive divergent behavioral and physiological outcomes from the same molecular perturbation. Taken together, our findings demonstrate that amidated peptides play central roles in establishing and maintaining sex differences in metabolic regulation, stress resilience, and behavior, with female physiology showing a particular dependence on gut-derived amidated signals.

### A functional atlas of sex-specific gut pathways and hormone signaling across metabolic, behavioral, and stress responses

To illustrate the extent of sex specificity across assays, we compared the distribution of hits in males and females. For this purpose, hits were defined as gene knockdowns whose phenotypic effects deviated more than ±1.5 standard deviations (SD; z-score ±1.5) from the control genotype, except for oxidative-stress survival and nighttime sleep (fed and starved), where a stricter cutoff of ±2 SD (z-score ±2) was applied. For each assay, we calculated the fraction of hits that were sex-specific as the proportion of knockdowns that produced a phenotype in only one sex relative to the total number of hits in that sex. This analysis revealed widespread sex specificity of knockdown phenotypes.

In most assays, the majority of hits – genotypes with extreme phenotypic effects in one sex – did not qualify as hits in the other sex when applying the defined z-score cutoff (Fig. 3a), indicating that gut-derived hormonal signals affect metabolic, behavioral, and stress-related traits differently in males and females. For example, 82% of nighttime-sleep hits identified in fed females were not shared with males, and likewise 86% of daytime-sleep hits in fed males were male-specific. Stress-resistance assays revealed many sex-specific knockdown hits for starvation and ionic stress, but a greater proportion of shared hits for oxidative stress. A quarter (25%) of female starvation hits and 18% of female ionic-stress hits also affected males, but oxidative-stress survival displayed substantial concordance, with 72% of female hits and 42% of male hits also observed in the other sex. Feeding duration similarly showed high overlap, with 73% of female hits and 45% of male hits shared with the other sex. These findings indicate that while many gut peptide knockdowns exert strongly sex-biased effects, certain physiological traits – particularly oxidative-stress survival, among the ones we measured – are shaped to a larger degree by genes and mechanisms operating in both sexes.

**Figure 3.**
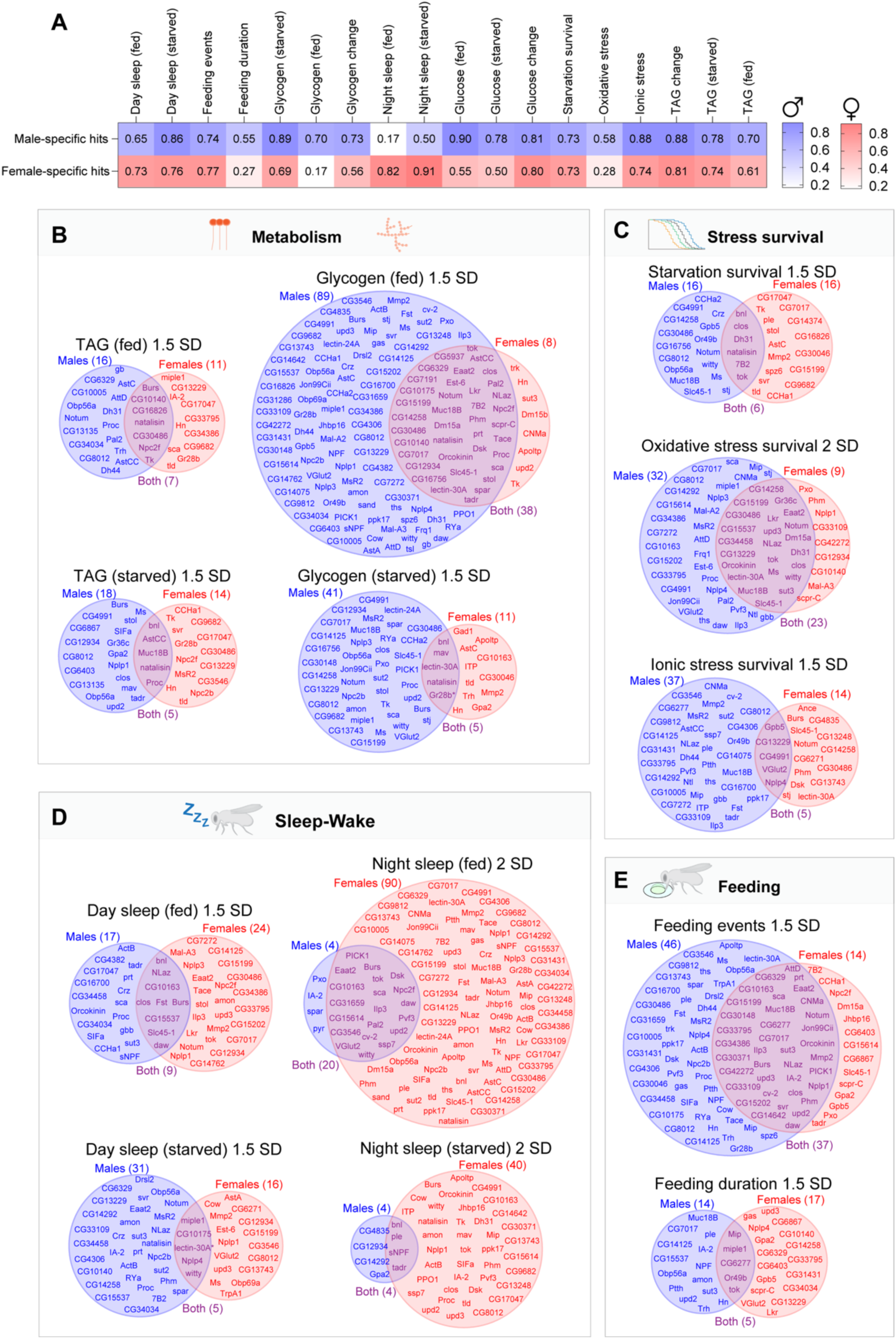
An atlas of sex-specific gut functions and endocrine signaling. Hits were defined as EEC gene knockdowns with phenotypic values outside ±1.5 standard deviations (SD; z-score ±1.5) from the control value (±2 SD; z-score ±2 for oxidative-stress survival and nighttime sleep assays). This definition was applied consistently across all subpanels. (a) Heatmap showing the fraction of sex-specific hits across all phenotypic assays. Blue indicates male-specific hits, and red indicates female-specific hits. (b–e) Venn diagrams showing the distribution of hits between male-specific (blue, left), female-specific (red, right), and both-sexes (purple, overlap) categories. (b) Metabolic traits: glycogen and TAG levels in fed and starved animals. (c) Stress resistance: survival under starvation, ionic stress, and oxidative stress. (d) Sleep behavior: daytime and nighttime sleep in fed and starved animals. (e) Feeding behavior: feeding events and feeding duration in the FLIC assay.

Building on this global comparison (Fig. 3a), Venn diagrams (Fig. 3b–e and S3a) illustrate the degree of overlap in gene-knockdown hits between sexes. This analysis indicates striking sex-specific differences in the effects of EEC-targeted gene knockdowns on systemic physiology and behavior. Metabolic traits, including TAG, glycogen, and glucose levels, exhibited clear sex-specific patterns, with male-biased effects generally more frequent. Glycogen levels in fed animals showed the strongest male-specific bias, with 89 genes affecting glycogen storage exclusively in males, compared to just eight in females (Fig. 3b). Thirty-eight knockdowns induced noted effects in both males and females, and thus 83% of knockdowns affecting females also affected males, and 70% of male hits were male-specific (Fig. 3b). Glucose levels also showed a strong male bias, with 44 male-specific and only 6 female-specific hits. By contrast, TAG levels – reflecting more stable long-term energy reserves – showed a less sex-biased pattern of 16 male-specific and 11 female-specific hits in fed animals, with seven hits shared between sexes. In line with this observation, starvation-resistance phenotypes, which generally correlate with energy stores such as TAG levels, were also evenly distributed, with 16 male-specific and 16 female-specific hits, plus six genes affecting survival in both sexes (Fig. 3c).

Survival in the face of oxidative and ionic stressors also displayed some male bias and sex-associated differences (Fig. 3c). Thirty-two genes gave male-specific knockdown effects on oxidative-stress resistance, whereas only nine genes gave female-specific phenotypes. Since this assay showed one of the largest overlaps between sexes, it might suggest that certain stress-response mechanisms, such as gut regenerative capacity following insults, are fundamental and shared between males and females. For example, gut-derived hormones such as Dh31 that locally regulate intestinal stem-cell proliferation^48^ may underlie such a shared resilience mechanism that limits epithelial damage caused by oxidative stress.

In contrast to metabolic traits, sleep behavior showed a pronounced female-specific bias (Fig. 3d). Nighttime sleep was affected by 90 genes in females, while only four genes produced male-specific effects. Most genes influencing male sleep also affected females, suggesting either broader endocrine sensitivity in females or reduced behavioral flexibility in males. One factor likely contributing to this imbalance is a ceiling effect: since males already sleep more at night, additional sleep-promoting effects will be harder to detect. However, no similar ceiling exists in the direction of reduced sleep, yet we observed only a few such cases, and their effects were relatively small. This lack of strong sleep-reducing phenotypes in males suggests that male sleep duration is constrained by biological factors beyond the endocrine pathways tested here.

Among all traits analyzed, feeding behavior showed the highest degree of gene overlap between sexes. In the FLIC (Fly Liquid-Food Interaction Counter)-based assay for feeding events, ∼38% of gene hits were not sex-specific, an overlap more than twice as large the average seen across other phenotypes (Fig. 3e). This suggests that gut-to-brain signaling pathways regulating feeding behavior are shared to a larger degree than those involved in metabolic or stress responses. On average, only 15% of hits strongly affected a given phenotype in both sexes (Fig. 3b-e and Fig. S3a) – meaning that six out of seven knockdowns with an extreme effect in one sex showed little or no effect in the other, highlighting a profound divergence in how gut-derived hormone signals shape physiology in males and females. These findings suggest that the animal’s sex has a large role in determining how physiological conditions interact with hormonal pathways and cellular processes.

Taken together, our results demonstrate that EEC-derived hormonal signals regulate a wide range of systemic traits in sex-biased or sex-specific ways. The sex-specific architecture of this endocrine network, encompassing both gut hormone production and the responsiveness of downstream tissues, may reflect evolutionary adaptations to differing energetic demands, reproductive roles, and behavioral strategies in males and females.

### Body fat stores correlate with nutritional- and oxidative-stress resistance and with sleep in females

We performed correlation analyses to explore how physiological traits relate to one another across EEC-specific gene knockdowns. This approach allowed us to identify global patterns of co-variation among metabolic, stress-related, behavioral, and sleep traits and to determine whether these correlations differ between sexes. Because females store more fat than males in both flies and humans^1-3^, we asked whether variation in lipid reserves relates to survival under nutritional and non-nutritional stress and to behavioral outcomes. To address this, we tested the correlation between TAG levels in fed animals across all knockdowns and survival under starvation, oxidative stress, and ionic stress, as well as with feeding behavior and sleep. Survival time under starvation showed a positive correlation with TAG levels in both sexes, as expected, as did resistance to oxidative stress, with these associations consistently stronger in females (Fig. 4a,b). Therefore, starvation survival and oxidative-stress resistance were also positively correlated with one another across knockdowns (Fig. S4a). In contrast, TAG levels did not correlate with resistance to ionic stress (Fig. 4c), indicating that lipid stores specifically buffer oxidative and nutritional stress, but not ionic imbalance. One explanation is that lipid droplets act as dynamic reservoirs that protect cells under nutrient limitation and oxidative stress by supplying energy and sequestering toxic lipid species, but they are not directly involved in maintaining ionic or osmotic balance, which depends primarily on ion transporters and membrane potential regulation rather than on lipid metabolism^49^. This suggests that energy storage and ionic-stress resistance are governed by distinct physiological mechanisms.

**Figure 4.**
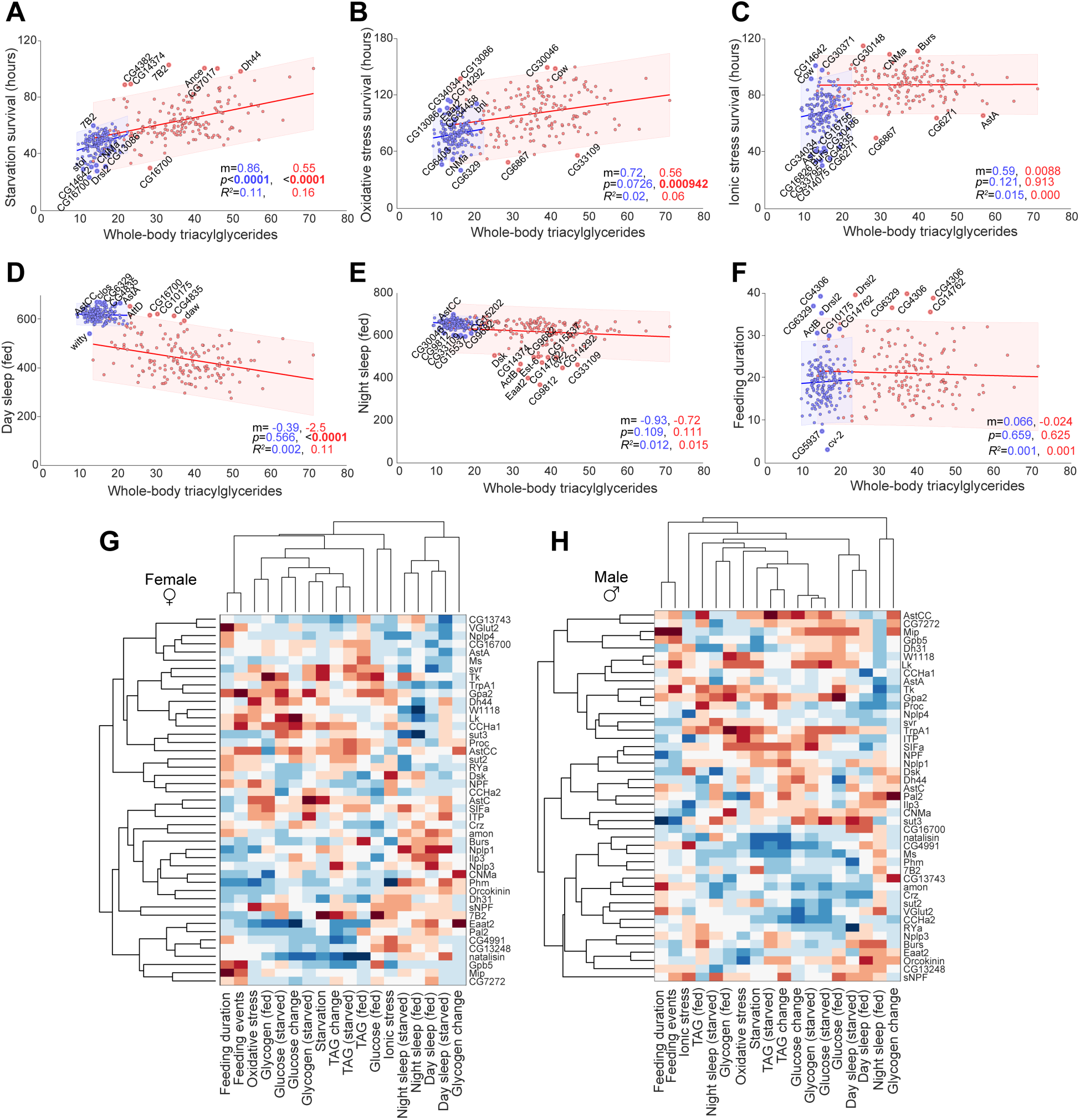
Correlations among phenotypic traits and functional clustering of EEC-derived signaling pathways. (a–f) Scatter plots for male (blue) and female (red) EEC-specific knockdowns, with whole-body triacylglyceride (TAG) levels on the X axis. The Y axis shows survival under (a) starvation, (b) oxidative stress, and (c) ionic stress; (d, e) time spent sleeping during the day (d) or night (e); and (f) feeding duration on 10% sucrose in the FLIC assay. Each dot represents an individual knockdown line. Linear fits are shown with shaded 95% confidence intervals. Knockdowns falling outside these intervals are labeled. Reported values indicate the slope (m), coefficient of determination (R2), and p-value for the slope for males (blue, left) and females (red, right). (g, h) Hierarchical clustering of phenotypes (columns) and selected EEC-specific knockdowns (rows) in females (g) and males (h). Heat maps show differences from controls, with red indicating higher values, white indicating control-like values, and blue indicating lower values. Phenotypic differences were converted to Z-scores, and clustering was based on Spearman’s rank correlation with average linkage. Analyses included secreted peptides, peptide-processing enzymes, and nutrient transporters. See Table S3 for pairwise correlations (rₛ and p values) between all traits.

When examining behavioral phenotypes, we found that daytime, but not nighttime, sleep was negatively correlated with TAG levels in the fed state (Fig. 4d,e). This raised the possibility that animals with greater energy reserves – and thus potentially more robust metabolic status – may require less sleep, or that sleep suppression facilitates energy accumulation, perhaps through increased opportunity for feeding. However, we found no correlation between TAG levels and feeding behavior (Fig. 4f and S4b), suggesting that enhanced energy storage is not simply a consequence of increased food intake during periods of reduced sleep. This also indicates that fat reserves do not strongly influence feeding duration, at least under the sugar-water-only conditions used in the FLIC assay. Taken together, these findings indicate that nutrient storage is more tightly linked to survival capacity in females, potentially reflecting sex-specific metabolic strategies that have evolved to promote male and female survival during periods of stress.

### Clustering analysis suggests possible functional relationships within gut nutrient sensing and hormone signaling

To identify global patterns within the dataset and uncover relationships among gene knockdowns across traits and sexes, we performed hierarchical clustering of all measured phenotypes for a selected subset of knockdowns in males and females (Fig. 4g,h). The selected gene subset included peptide hormones, processing enzymes, sugar and amino-acid transporters, and the heat- and oxidative-stress-activated cation channel Transient Receptor Potential A1 (TrpA1). This unsupervised approach allowed related phenotypes and genotypes to group together based on similarity across multiple traits, and it uncovered several examples where genes with established functional relationships clustered closely, suggesting that the screen captured biologically meaningful functional modules. Phenotypes were assessed based on Z-scored phenotypic differences from the control genotype, allowing us to identify genes that regulate physiology and behavior in similar ways and ensuring that observed clustering patterns reflect biologically meaningful deviations from baseline.

#### In females

In females, the cation channel TrpA1 and the peptide Tk co-cluster (Fig. 4g and Table S2), with highly similar knockdown phenotypes (Spearman’s rank correlation r_s_ = 0.63 across the panel of 18 traits). We recently demonstrated that TrpA1 activity in EECs regulates gut Tk secretion in response to oxidative stress and that this pathway is also required for Tk release following dietary amino-acid intake^19^.

Another example of clustering that aligns with prior work is the grouping of sugar transporter 2 (Sut2) with NPF (r_s_ = 0.21 across the 18-trait panel) consistent with reports that Sut2 is required for EEC-mediated NPF secretion in response to dietary sugar^8^. These examples illustrate that genes with known related physiological and signaling relationships indeed cluster together in our data, supporting the idea that other clustering potentially reflects novel functional relationships and regulatory modules within the gut endocrine system.

Our analysis revealed an interesting cluster in female flies comprising knockdown of genes encoding predicted amino-acid transporters (CG13743 and CG16700) and genes encoding the peptides AstA, Myosuppressin (Ms), and Neuropeptide-like precursor 4 (Nplp4) (Fig. 4g and Table S2). CG13743 is the fly ortholog of mammalian SLC38A11, and CG16700 is orthologous to mammalian SLC36-family transporters. AstA has been linked to regulation of feeding behavior, metabolism, and sleep^50-52^. The clustering of these genes indicates a potential functional relationship operating in a subset of EECs. We speculate that the amino-acid transporters CG13743 and CG16700 might sense dietary amino-acid availability and subsequently modulate the secretion or action of AstA, Ms, and Nplp4. Such a mechanism could enable EECs to directly couple dietary protein content to hormonal signals that regulate metabolism and feeding behavior.

NPF clusters with other peptides CCHamide-2 (CCHa2) and Drosulfakinin (Dsk) across some metabolic and behavioral traits, particularly those affecting lipid and glucose levels, as well as sleep (Fig. 4g and Table S2). Both NPF and CCHa2 are well-established regulators of feeding and metabolism^8,36,53-56^, and their clustering together suggests a coordinated role in modulating systemic energy balance and stress responses through gut-to-brain signaling pathways. Another interesting group includes AstC and the peptide SIFamide (SIFa), which display similar phenotypic profiles, especially in relation to feeding behavior, glycogen and TAG levels, and starvation resistance. This suggests that AstC and SIFa may act in parallel to regulate nutrient storage and mobilization under nutritional stress.

In females, the peptides CNMamide (CNMa), Orcokinin, and Dh31 cluster with Phm, reflecting similar knockdown phenotypes. Since Phm is required for amidation of CNMa and Dh31, this suggests that loss of EEC-derived CNMa and Dh31 may drive some of the phenotypes observed in Phm knockdown (Fig. 4g and Table S2). The knockdown of these genes is associated with reductions in feeding, oxidative-stress resistance, starvation resistance, and TAG levels, suggesting a shared function in stress-related metabolic regulation (Fig. 4g). Finally, a cluster containing Glycoprotein hormone β5 (GpB5), Myoinhibitory peptide (Mip), and CG7272 shows strong phenotypic signatures characterized by increased feeding behavior. Interestingly, EEC loss of Glycoprotein hormone α2 (GpA2), which forms a heterodimer with GpB5 to activate signaling through Leucine-rich repeat-containing G protein-coupled receptor 1 (LGR1), the cognate receptor for the GpA2:GpB5 heterodimer^57^, also promotes feeding but does not cluster with this group (Fig. 4g and Table S2). This supports the hypothesis that the GpA2/GpB5 heterodimer may function as a gut-derived hormone that modulates appetite. CG7272 encodes a predicted sugar transporter orthologous with mammalian SLC50A1 (SWEET), suggesting it may play a role in regulating the secretion or activity of GpB5 or Mip in EECs in response to sugar consumption.

#### In males

Consistent with females, GPB5 and Mip form a cluster in males and negatively regulate sugar feeding, as loss of either elevates consumption (Fig. 4h and Table S2). This consistent phenotype across sexes suggests that GpB5 and Mip may function as gut-derived hormones suppressing appetite in both males and females. In males, Tk clusters with TrpA1, as observed in females. This clustering in both males and females, together with our more-detailed, independent description of their functional relanionship^19^, indicates that this gut hormone serves similar functions in both sexes. In contrast to the female data, where NPF-knockdown phenotypes cluster with those of the sugar transporter Sut2, the male data do not give rise to such a grouping (Fig. 4h). In females, NPF knockdown phenotypes cluster with those of the sugar transporter Sut2, which is required for EEC-mediated NPF secretion in response to dietary sugar in females^8^. In males, this grouping is absent (Fig. 4h and Table S2). This apparent decoupling suggests sex-specific regulation of NPF and differences in its functional partners.

Another intriguing male-specific cluster includes CNMa, sugar transporter 3 (Sut3), and the amino-acid transporter CG16700 (Fig. 4h and Table S2). Knockdown of these genes is associated with reduced feeding behavior. Whereas CNMa has been shown to regulate feeding decision-making when secreted from enterocytes^58^, its role in EECs remains to be defined. Given its clustering with Sut3 and CG16700 – both putative nutrient transporters – our data suggest that CNMa may also act as an EEC-derived signal modulating appetite in response to nutrient intake, possibly through regulatory mechanism involving these transporters. This highlights a potential nutrient-sensing regulatory module in the gut that promotes feeding in response to dietary cues.

### Functional analysis indicates that EEC-derived signals drive sex differences in energy balance and behavior

To compare sex-specific physiological and behavioral patterns more effectively between males and females, we clustered absolute phenotypic values (for example, total TAG stores) rather than the Z-scored differences from controls used in Fig. 4g,h. This approach allows us to assess whether knockdown of individual genes shifts phenotypes of a male or female genotype toward a more male-like or female-like profile. By analyzing absolute values, we aimed to identify gene perturbations that either reinforce or disrupt sex-specific traits, thereby revealing the molecular signals that contribute to the establishment and maintenance of metabolic and behavioral states in males and females. Clustering revealed robust, consistent differences between males and females (Fig. 5a and Table S3), consistent with well-documented sex-dependent variation in traits such as sleep, lipid stores, starvation resistance, and feeding behavior. Across the screen data, females generally exhibited a higher number of feeding events and longer feeding duration, along with increased energy stores (TAG and glycogen), enhanced resistance to starvation, oxidative, and ionic stress, and lower total sleep compared to males. The increased feeding and energy levels observed in females is consistent with their higher energy demands and storage capacity^4,6^. These sex-specific traits drive the clustering analysis to organize the data into distinct male and female branches, with sex emerging as a major axis of phenotypic variation across EEC gene knockdowns.

**Figure 5.**
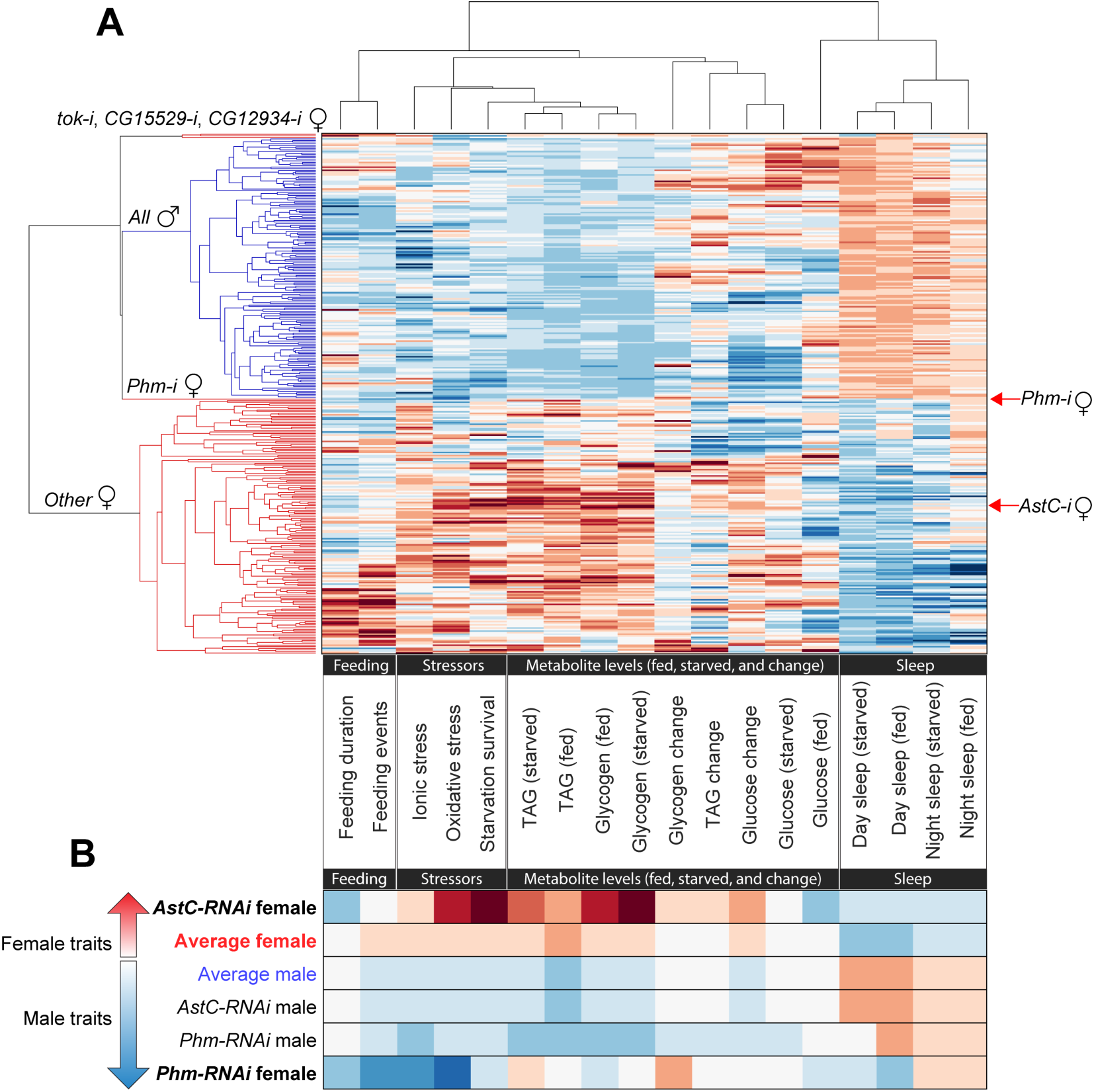
Clustering shows sex-specific metabolic and behavioral traits dependent on gut signaling. (a) Hierarchical clustering of absolute phenotypic values (columns) across feeding, stress-resistance, metabolic, and sleep traits for EEC-specific knockdowns (rows) in females (red branches) and males (blue branches). Heat maps show trait values, with blue indicating lower values, white the cross-sex average, and red higher values. (b) Heat map of representative traits comparing average performance within males and females to that of Phm-RNAi and AstC-RNAi animals. The “average” genotype was calculated for visualization only and not included in the clustering in (a). See Table S4 for pairwise correlations (rₛ and p values) between all traits.

Strikingly, however, knockdown of the peptide-processing enzyme Phm did not follow this pattern. Among the 165 genes analyzed, *Phm* was the only one whose knockdown in EECs led to a female phenotypic profile that clustered among the male knockdowns (Fig. 5a,b and Table S3), suggesting that amidated gut peptide hormones together drive physiology and behavior across a broad range of traits toward a female-like state. Phm-mediated amidation enhances peptide stability and receptor-binding affinity, and it is generally required for full hormonal function^45^. In *Drosophila*, a large fraction of known EEC-secreted peptides – including AstA, Tk, NPF, Dh31, and others – depend on amidation for their activity. Thus, knockdown of Phm in adult EECs likely impairs the function of a wide range of gut peptide hormones and broadly disrupts gut hormone signaling, while leaving other secreted factors unaffected, including cytokines, growth factors, biogenic amines, and non-amidated peptides.

The clustering of Phm-knockdown females into the male branch of the hierarchical tree (Fig. 5a,b and Table S3) suggests that the loss of bioactive amidated peptides disproportionately affects female physiology, which may rely more heavily on gut hormone signals for maintaining higher energy stores, stress resistance, and specific behavioral patterns such as reduced sleep. EEC knockdown of Phm in females reduced TAG and glycogen levels, decreased survival under starvation, ionic, and oxidative stress, and increased sleep, thereby mirroring the male physiological profile. Feeding activity was also reduced, further aligning with male-like behavior and suggesting that amidated gut hormones normally promote higher nutrient intake, energy storage, and stress resilience in females. These findings suggest that amidated gut peptides play a key role in establishing and maintaining female-typical physiology, including elevated energy storage, stress resistance, and distinct activity and sleep patterns.

Unlike many other gut peptides, AstC is not amidated and thus does not require Phm for its activity. In contrast to the broad loss of amidated peptides caused by knockdown of Phm, knockdown of the non-amidated AstC did not shift females toward the male cluster (Fig. 5a,b and Table S3) but instead reinforced the female-like profile across most traits. EEC-targeted AstC has previously been shown to increase the storage of fat and to reduce its mobilization, leading to increased starvation resistance^20^, traits that are associated with female physiology. Consistent with this, EEC AstC knockdown in females clustered with gene knockdowns that promote female-typical traits, including increased energy storage, greater stress resistance, and reduced sleep. Thus, loss of AstC in the EECs enhanced female-typical traits, suggesting that AstC shapes female physiology in the male-like direction, in opposition to Phm-dependent amidated gut peptide signals that collectively drive energy storage and reduced activity.

### Amidated gut peptide signaling promotes female physiological and behavioral states and is counterbalanced by non-amidated gut signals

To obtain a more global assessment of sex-specific EEC-driven effects, we next analyzed the impact of knocking down individual peptide genes and peptide-processing enzymes required for the generation of amidated versus non-amidated peptides. A large fraction of the genes targeted in our screen encode peptide hormones that require enzymatic processing before secretion. These maturation steps determine which subsets of gut signals are produced, and thus whether EECs can generate amidated versus non-amidated peptides. Secreted factors first enter the canonical secretory pathway, where signal peptides are removed, after which prohormones undergo specific cleavage and modification steps. Initial cleavage at dibasic motifs is performed primarily by the prohormone convertase Amontillado (Amon)^46^. Basic residues exposed by this cleavage are then trimmed from the new C terminus by the carboxypeptidase Silver (Svr)^47^. Peptides ending in glycine are further amidated by Phm together with Peptidyl-α-hydroxyglycine-α-amidating lyase (PAL). These sequential steps yield mature amidated peptides such as Tk, NPF, AstA, Dh31, and PDF, while others like AstC require Svr but not Phm, and still others such as AstCC require only Amon cleavage. Thus, targeting *amon*, *svr*, and *Phm* in EECs disrupts overlapping but distinct subsets of gut hormones (Fig. 6a,b). This strategy provided a systems-level framework for understanding how broad classes of peptide signals – amidated versus non-amidated – contribute to the regulation of energy balance and behavior in a sex-specific manner.

**Figure 6.**
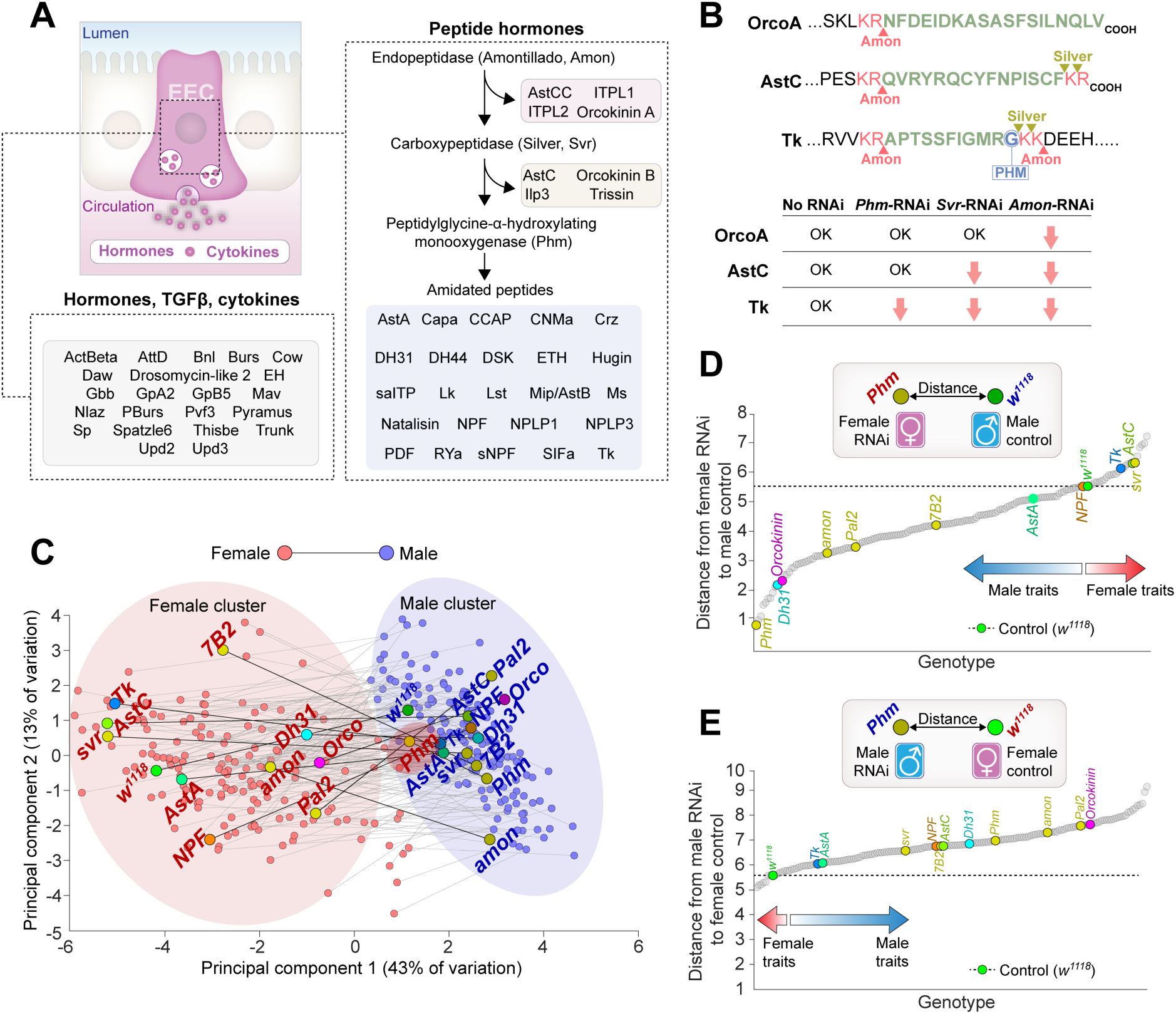
Amidated and non-amidated gut peptides drive female-specific physiology in opposing directions. (a) Schematic of enteroendocrine cell (EEC) hormone secretion and peptide-processing steps. Prohormones are cleaved by Amontillado (Amon), trimmed by Silver (Svr), and amidated by Peptidylglycine-α-hydroxylating monooxygenase (Phm) together with Peptidyl-α-hydroxyglycine-α-amidating lyase (PAL). Processed peptides include amidated (*e.g.*, Tk, NPF, AstA, Dh31, PDF) and non-amidated (*e.g*., AstC, AstCC) hormones. (b) Representative peptide sequences showing predicted cleavage and modification sites. Red arrows indicate processing steps disrupted by RNAi targeting *Amon*, *Svr*, or *Phm* in EECs. (c) Principal-component analysis (PCA) of absolute phenotypic values. Male (blue) and female (red) samples of the same genotype are linked by a thin line. Selected knockdowns are labeled. (d) Euclidean distance in the PCA1 x PCA2 plane from each female knockdown to the male control. The schematic inset illustrates how the distance was calculated between female RNAi samples and the male control. The control (*voilà*> x *w^1118^*) is shown in green. (e) Euclidean distance in the PCA1 x PCA2 plane from each male knockdown to the female control. The schematic inset illustrates how the distance was calculated between male RNAi samples and the female control. The control (*voilà*> x *w^1118^*) is shown in green. Statistics: Phenotypic values were converted into Z-scores, and PCA was performed with centering and the singular value decomposition method.

To further quantify how EEC-derived factors contribute to sex-biased physiology, we performed principal component analysis (PCA) of absolute phenotypic values across all genotypes (Fig. 6c). This approach organizes animals in “phenotypic space” according to shared metabolic and behavioral traits. As expected, control males and females separated into distinct clusters, reflecting their characteristic physiological differences. Most EEC knockdowns retained this separation, but knockdown of *Phm* in females caused a striking shift toward the male cluster, consistent with the idea that amidated peptides broadly promote female-typical traits. To assess the extent to which individual knockdowns shifted traits toward the opposite sex, we calculated the distance in PCA space of each female knockdown to the male control (Fig. 6d). In multivariate phenotypic space, females with EEC *Phm* knockdown clustered closest to the male control among all female knockdowns, indicating that loss of peptide amidation disproportionately reduces female-typical features such as elevated energy storage and stress resistance. Strikingly, knockdown of the single peptide *Dh31* also shifted females strongly toward males, suggesting that Dh31 is one of the major amidated gut hormones driving female physiology, and that the broad effects of *Phm-RNAi* may in part reflect loss of *Dh31*. In contrast, EEC knockdown of *AstC* shifted females further from males, reinforcing female-like states and suggesting that gut-derived AstC acts in an antagonistic manner to amidated peptides such as Dh31. Removing the processing enzyme *Svr* in EECs – which eliminates both Phm-dependent peptides and AstC – produced the same female-like shift as AstC knockdown alone. This divergence from *Phm-RNAi* indicates that AstC normally acts to suppress female-typical traits, counteracting the action of amidated gut peptides. Moreover, the similarity between *Svr-RNAi* and *<u>AstC-</u> <u>RNAi</u>* suggests that the loss of AstC dominates over the loss of amidated peptides, placing AstC function epistatic to that of amidated gut hormones in shaping systemic physiology.

We next performed the reciprocal analysis, measuring the PCA-space distance of male knockdowns to the female control (Fig. 6e). Unlike females, most male knockdowns shifted only modestly toward female-like traits, and very few moved closer to the female control. This asymmetry indicates that female physiology is more plastic and more strongly dependent on amidated peptide signaling, whereas male physiology is less readily shifted toward a female-like state. Finally, we quantified the distance between male and female animals for each knockdown pair (Fig. S5a). *Phm-RNAi* pairs showed the smallest sex difference, consistent with amidated peptides broadly driving male-*vs.*-female divergence in physiology and behavior. By contrast, *Svr-RNAi* and *AstC-RNAi* pairs displayed among the largest sex differences, highlighting that EEC-derived AstC normally drives processes whose effect tends to dampen sexual dimorphism. Together, these analyses reveal that different classes of gut-derived signals exert opposing influences on systemic physiology: amidated peptides (including Dh31) enhance female-typical traits more in females than in males and thereby expand sex differences, while AstC promotes male-like states – again, to a greater degree in females than in males – and thus has the effect of limiting dimorphism. More broadly, the PCA framework provides a systems-level view of how ensembles of gut hormones coordinate to establish and maintain sex-specific metabolic and behavioral phenotypes.

### EEC-derived peptide hormones promote female energy storage, confer stress resistance, and direct food choice

To determine whether the sex-specific effects observed in our screen were mediated directly by gut hormones, we restricted RNAi to the adult EEC population with specific removal of neuronal effects. For this, we used *voilà*> in combination with *R57C10-GAL80*, which constitutively inhibits GAL4 in all neurons, a combination hereafter referred to as *voilà^GUT^>*. Using this system, we found that EEC-specific knockdown of *Phm* reproduced the strong female-specific phenotypes detected in the broader screen. Females with EEC-specific *Phm* loss were as susceptible to starvation as males (Fig. 7a), indicating that amidated gut peptides are required to sustain the enhanced starvation resistance normally seen in females. Similarly, *AstC* knockdown using *voilà^GUT^>* also matched the results obtained with the potentially broader *voilà*> driver (Fig. 7b), consistent with earlier findings that *voilà*> primarily reduces gut rather than brain AstC^20^. EEC-specific *Phm* loss also uncovered striking female-specific susceptibility to oxidative stress (Fig. 7c), again consistent with the screen data (Fig. 2b). By contrast, *AstC* knockdown increased oxidative-stress resistance in both sexes, without generating a sex difference (Fig. S6a). These findings support that amidated peptides, but not AstC, are essential for the enhanced nutrient and oxidative stress resistance that distinguish females from males.

**Figure 7.**
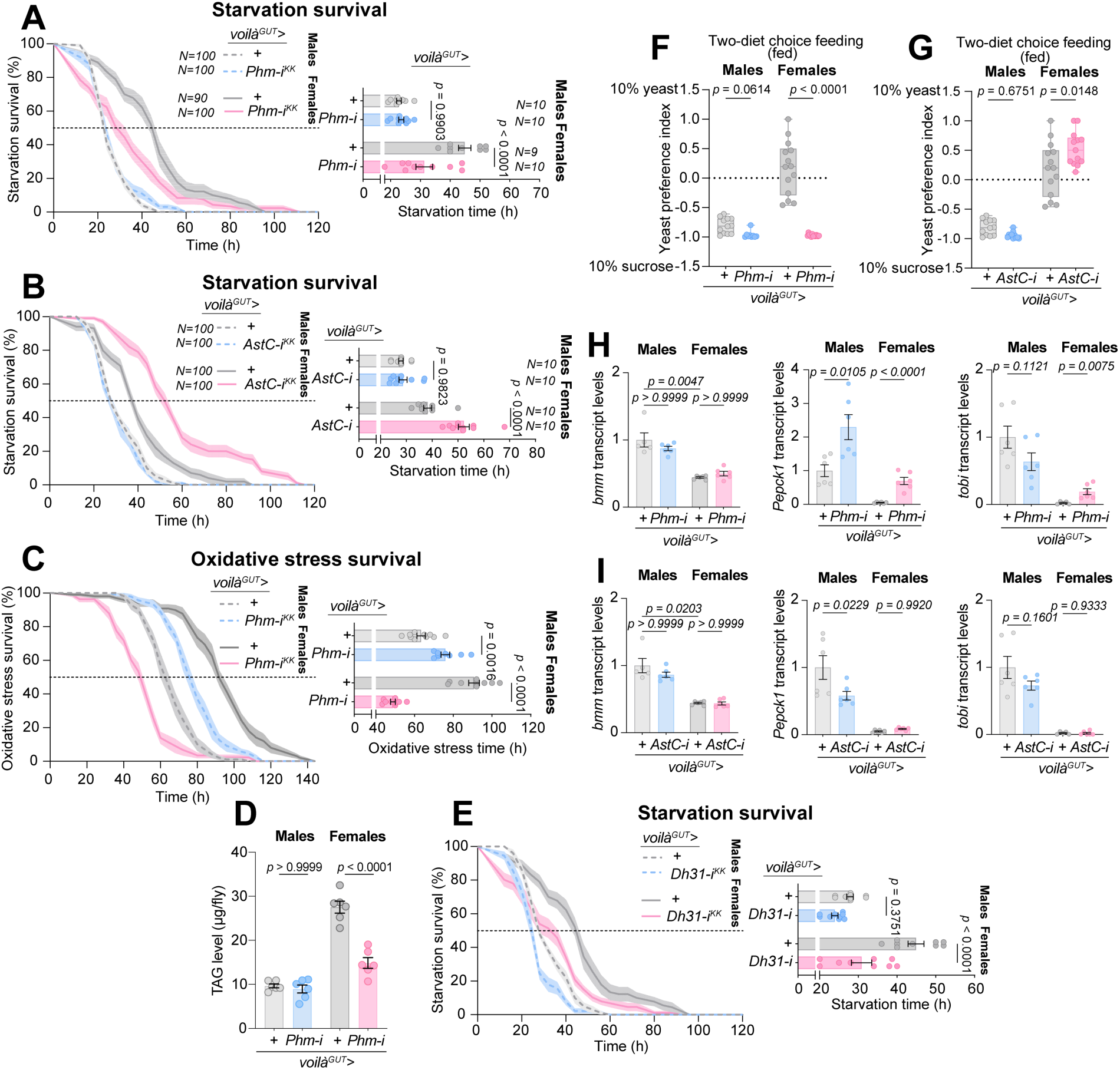
EEC-derived peptide hormones promote female energy storage, confer stress resistance, and bias feeding preferences. (a–c) Survival under starvation (a, b) and oxidative stress (c) upon EEC-specific knockdown of AstC or Phm using *voilà^GUT^>* (*voilà>* combined with *R57C10-GAL80* to restrict knockdown to adult EECs). (d) Triacylglyceride (TAG) levels in males and females with EEC-specific *Phm* knockdown. (e) Survival under starvation upon EEC-specific knockdown of *Dh31*. (f, g) Two-diet choice feeding assays with EEC-specific knockdown of *Phm* (f) or *AstC* (g). (h, i) Expression of metabolic regulators (*bmm*, *PEPCK1*, *tobi*) in males and females upon EEC-specific knockdown of *Phm* (h) or *AstC* (i). Boxplots show median and interquartile range with individual data points; bar graphs show mean ± s.e.m. Statistical tests: survival assays (a–c, e) were analyzed by one-way ANOVA with Tukey’s post hoc tests; TAG levels (d) and feeding assays (f, g) by unpaired two-sided *t-*test; and gene-expression assays (h, i) by one-way ANOVA with Tukey’s post hoc multiple comparisons or unpaired two-sided *t*-test (pairwise comparisons).

TAG storage is a central determinant of resistance to both starvation and oxidative stress, and females characteristically store more lipids than males. We therefore asked whether amidated gut peptides contribute to this sex-specific energy-storage state. EEC-specific knockdown of *Phm*, but not of *AstC*, strongly reduced TAG levels in females, while having no effect in males (Fig. 7d and S6b). This loss of female-specific fat storage effectively shifted female TAG levels to the lower male baseline, establishing that amidated gut peptides are critical determinants of female energy reserves. To identify which specific amidated peptide(s) account for this effect, we tested individual candidates. Knockdown of *Tk* in the EECs did not reduce starvation resistance in either sex (Fig. S6c), ruling out a major role for gut-derived Tk in establishing sex-biased energy storage. Knockdown of *NPF* produced a modest increase in female starvation resistance (Fig. S6d), consistent with its reported inhibition of the catabolic hormone AKH^8,36^, but insufficient to explain the strong phenotype of *Phm* loss. In contrast, EEC-specific loss of *Dh31* nearly phenocopied *Phm* knockdown (Fig. 7a,e), strongly implicating *Dh31* as a key mediator of female-biased energy storage. While Dh31 has previously been associated with feeding regulation in males^35^, its role in female systemic metabolism has remained unexplored. Our findings identify Dh31 as a likely major EEC-derived signal that promotes elevated energy reserves and stress resistance in females. Together, these results demonstrate that amidated gut peptides are essential drivers of the female metabolic state, with Dh31 emerging as a primary candidate underlying the sex-specific enhancement of nutrient storage and resilience.

Given these strong effects on energy storage and stress resistance, we next asked whether EEC-derived amidated peptides also influence feeding behavior, a major sex-biased trait in flies. Females, owing to their higher reproductive energy demands, more strongly prefer protein-rich yeast food than males do^8,9^. This preference has been linked to gut-brain signaling through NPF, which suppresses the catabolic hormone AKH. In mated females, reduced AKH activity not only limits AKH-driven energy mobilization but also shifts feeding preference away from sugar and toward protein-rich yeast^8^. Conversely, AstC promotes AKH release^20^, and since AKH is a key hormone that favors sugar intake over protein consumption^19^, AstC activity would be expected to bias food choice in the opposite direction of NPF. Since loss of *Phm* rendered females male-like in their starvation resistance, we tested whether it also altered their feeding preferences. Indeed, EEC-specific *Phm* knockdown abolished the female preference for yeast and shifted feeding behavior toward a male-like, sugar-biased pattern (Fig. 7f). In contrast, EEC-specific loss of *AstC* led to increased yeast preference in females, but not males (Fig. 7g), consistent with its AKH-stimulatory role and with our broader observation that AstC loss enhances female-associated traits. Together, these findings show that EEC-derived amidated peptides not only sustain the female metabolic state of high energy storage and stress resistance but also coordinate sex-specific feeding preferences.

### EEC-derived peptides modulate systemic insulin and AKH signaling in a sex-specific manner

We next asked whether the metabolic effects of gut hormone loss could be explained by alterations in the two key hormonal pathways that govern energy storage and utilization in insects, insulin and AKH^10^. These pathways are central determinants of sex differences in fat storage, with females generally exhibiting higher anabolic insulin signaling and lower AKH activity^6,13,59^. To address this, we measured transcript levels of insulin- and AKH-pathway components as well as their downstream effectors. Knockdown of *AstC* or *Phm* in EECs did not alter expression of the insulin-like peptides *Ilp2* and *Ilp3* – two of the main insulins produced by the insulin-producing cells (IPCs) – or *Akh* and its receptor, consistent with the notion that these hormones are primarily regulated at the level of secretion rather than transcription (Fig. S6e,f).

For *AstC* knockdown, no female-specific differences in insulin-pathway activity were detected when assessing *Eukaryotic translation initiation factor 4E-binding protein* (*4EBP*) or the insulin receptor (*InR*) (Fig. S6f). 4EBP is a translational repressor whose expression is suppressed by insulin signaling, and thus serves as a sensitive transcriptional readout of systemic insulin activity^60^. InR itself is also negatively regulated by insulin signaling, such that elevated transcript levels reflect reduced pathway activity^61^. Although *AstC* knockdown did not alter these markers, we noted that *4EBP* levels were consistently lower in females than in males, indicating higher systemic insulin activity in females, consistent with female-biased insulin signaling that promotes anabolism and greater energy storage. By contrast, *Phm* knockdown in EECs increased *4EBP* expression in both sexes, indicating reduced insulin-pathway activity when amidated gut peptides are absent (Fig. S6e). This was further supported by elevated *InR* expression in females upon *Phm* knockdown, again consistent with diminished insulin signaling and in line with the observed reduction in fat storage in these animals.

We next examined metabolic regulators downstream of insulin and AKH signaling. The triglyceride lipase Brummer (Bmm), the *Drosophila* homolog of adipose triglyceride lipase (ATGL), is a key enzyme driving lipolysis. Its expression is normally lower in females than in males, consistent with reduced basal lipolytic activity and greater fat storage in females^4^. Neither *AstC* nor *Phm* knockdown significantly affected *bmm* expression, suggesting that altered lipolysis is not the primary mechanism underlying the observed metabolic changes (Fig. 7h,i). In contrast, *phosphoenolpyruvate carboxykinase 1* (*Pepck1*), encoding the rate-limiting enzyme of gluconeogenesis, was strongly upregulated in *Phm* knockdown animals. Elevated *Pepck1* expression signals a shift toward catabolism and increased energy expenditure, providing a plausible explanation for the reduced triacylglyceride (TAG) storage in females lacking amidated gut peptides. Finally, we analyzed *target of brain insulin* (*tobi*), which encodes an α-glucosidase that mobilizes glycogen and sugar. *Tobi* is unusual in being transcriptionally regulated by both insulin and AKH, thereby serving as a readout of the balance between these two antagonistic pathways^62^. Intriguingly, *tobi* expression responded to *Phm* knockdown in a sex-specific manner, with females exhibiting increased expression while males showed a tendency towards reduced expression (*p*=0.11). These data suggest that amidated gut peptides fine-tune the balance between insulin and AKH signaling in a sex-specific fashion. In females, increased *tobi* expression upon *Phm* knockdown likely accelerates sugar mobilization and contributes to energy-store depletion, whereas the opposite trend in males highlights divergent transcriptional regulation between sexes. Taken together, these findings indicate that amidated-peptide signaling is a central mechanism through which EECs regulate systemic and sex-specific metabolic homeostasis and feeding behavior, and that disrupting this signaling in females can partially shift systemic physiology toward a male-like state.

## Discussion

Our study demonstrates that EEC-derived signals exert profound and sex-dependent influences on systemic physiology and behavior in *Drosophila* and that sex-specific anatomical differences exist in the EEC population itself. By systematically interrogating gut-derived peptide signaling, we reveal that amidated peptides as a group broadly sustain female-typical traits, including elevated lipid storage, enhanced stress resistance, and altered feeding and sleep behavior, while the non-amidated peptide AstC acts in the opposite direction. These findings establish a functional framework in which ensembles of gut hormones and their processing enzymes sculpt differences in male and female physiological states.

Sex-specific differences in gut-hormone biology are not unique to flies. In mammals, emerging evidence suggests that incretin-based therapies targeting glucagon-like peptide-1 (GLP-1) exhibit sexually dimorphic effects on energy balance and glucose metabolism^63,64^. Women generally exhibit stronger insulinotropic responses to GLP-1 receptor agonist therapy than men, and several clinical studies have reported greater reductions in body weight and glycemic measures in women than in men^65,66^. Although the mechanisms underlying these trends remain incompletely defined, possible contributing factors include sex-specific differences in GLP-1 secretion, receptor distribution, estrogen signaling, and central nervous system responses. Beyond GLP-1, other gut hormones also show sex-dependent actions in mammals. Two other key appetite-regulating hormones, peptide YY (PYY) and cholecystokinin (CCK), have also been linked to sex-specific effects, with CCK in particular producing a stronger induction of satiety in women^67,68^. Together, these observations suggest that sex-specific endocrine architecture of the gut, and the sex-dependent responsiveness of downstream tissues, are conserved themes across animals. These findings argue that sex should be considered a central variable in the study of gut hormones, both in basic research and in the clinical application of gut-hormone–based therapies.

An important open question concerns the nature of the mechanisms driving the sex differences we observe in gut endocrine signaling. Several upstream regulatory systems may contribute to establishing or maintaining these dimorphisms. In *Drosophila*, the sex-determination gene *transformer* (*tra*) shapes sexual identity across diverse tissues^69^, and *tra*-dependent programs could extend to the gut epithelium, influencing EEC fate or function in a sex-specific manner. In parallel, systemic hormones with broad roles in development and adult physiology – ecdysone and juvenile hormone (JH) – represent plausible modulators of gut endocrine function. In the gut, ecdysone signaling regulates metabolism and growth, and also controls intestinal stem-cell activity in adults^26,70,71^, whereas JH has been implicated in gut remodeling^38^. Both pathways could therefore tune sex differences in gut-hormone production or release.

In mammals, sex steroids play an important role in metabolic control^72^. Although insect ecdysteroids have not traditionally been considered sex steroids, little is known about their roles in adults. In mammals, testosterone serves as a precursor to estrogen^73^. Likewise, in insects, ecdysone (E) is converted to active molting hormone 20-hydroxyecdysone (20E), but there is evidence that E itself has biological roles independent of its conversion to 20E^74,75^, and E may even be synthesized by the male reproductive system in insects^74^. Moreover, during certain developmental stages, E, rather than 20E, is the predominant ecdysteroid^76^, suggesting additional roles for the precursor E. An intriguing possibility is that different ecdysteroids could act as functional ‘sex hormones’ in *Drosophila* and other insects. However, whether males and females differ in the production, metabolism, or gut sensitivity to these ecdysteroids remains unexplored.

Our findings, together with previous reports, suggest that gut-derived peptides shape systemic metabolism in a sex-dependent manner by intersecting with the two central hormonal systems – insulin and glucagon-like signaling, which antagonistically regulate energy storage and mobilization^8,19,20^. In *Drosophila*, AKH serves as the principal glucagon-like factor, mobilizing lipids and carbohydrates during nutrient stress, while insulin-like peptides (Ilps) promote anabolic storage and growth^10,60^. Previous work has shown that females typically exhibit higher basal insulin pathway activity and lower AKH activity than males, consistent with their larger lipid stores and increased starvation resistance^6,13,59^. A main route through which gut hormones generate sex-specific effects in males and females is likely via their modulation of insulin and glucagon-like signaling. In mammals, comparable sex differences are evident in insulin and glucagon signaling. Like to *Drosophila* females, human females generally exhibit greater insulin sensitivity and anabolic signaling^77,78^. Our findings in flies suggest that such sex differences may arise, at least in part, through modulation by gut-derived peptides – a mechanism that could likewise contribute to sex-specific metabolic regulation in humans.

It also remains an open question at which level sex differences in gut endocrine signaling are established. Our data show that at least part of the dimorphism originates within the gut epithelium itself. Females possess a greater number of AstC-positive EECs, which likely enhances female-specific endocrine output. Additional layers of regulation may exist at the level of target tissues, such as the insulin-producing cells (IPCs) of the brain or the adipokinetic hormone-producing cells (APCs) of the *corpora cardiaca*, where sex-specific receptor abundance, receptor sensitivity, or downstream signaling capacity could shape distinct systemic outcomes from the same gut-derived signals. Moreover, peripheral tissues such as fat body, muscle, or gonads could contribute through differential feedback regulation or nutrient demand. Dissecting whether sex-biased effects arise from the source (EECs), the sensors (receptor-bearing cells), or the responding tissues remains a key future challenge.

## Methods

### *Drosophila* husbandry and stocks

Fly stocks were maintained at 18 °C and 60% relative humidity on standard cornmeal medium (82 g/L cornmeal, 60 g/L sucrose, 34 g/L yeast, 8 g/L agar, 4.8 mL/L propionic acid, and 16 mL/L 10% methyl-4-hydroxybenzoate in ethanol). Animals used for experiments were reared to adulthood in incubators at 18 °C (to suppress GAL4 activity when GAL80^TS^ was present) or at 25 °C (for experiments not requiring GAL80). After eclosion, flies were transferred to an adult-optimized cornmeal-free medium (90 g/L sucrose, 80 g/L yeast, 10 g/L agar, 5 mL/L propionic acid, and 15 mL/L 10% methyl-4-hydroxybenzoate in ethanol). A 12-hour light/dark cycle was maintained throughout development and adult culture.

The *voilà-GAL4* driver, an insertion in the *prospero* locus, was used to direct UAS transgene expression in all enteroendocrine cells (EECs). For adult-specific expression, *voilà-GAL4* was combined with *Tubulin-GAL80^TS^*(together *voilà*>), and for gut-specific expression, *voilà-GAL4* was combined with *R57C10-GAL80* (“*voilà^GUT^*>”). *AstC::T2A::GAL4* and *Tk::T2A::GAL4* knock-in drivers at the endogenous loci were used to label AstC- and Tk-expressing EECs, respectively, with *UAS-GFP*. RNAi lines (Table S4) were obtained from Vienna *Drosophila* Resource Center (VDRC). Other lines were obtained from Bloomington Drosophila Stock Center (BDSC), including *AstC::T2A::GAL4* (#84595)^79^, *UAS-mCD8::GFP* (#5137), *Tk::T2A::GAL4* (#84693)^79^, and *Tub-GAL80^TS^* (#7108). The *voilà-GAL4* driver line^80^ and the *R57C10-GAL80-658-63*^81^. All *GAL4* and *GAL80* driver lines were backcrossed for several generations into our in-house *w^1118^* background to ensure a consistent genetic context. For experimental crosses, these drivers were mated to *UAS-RNAi* lines from VDRC. Controls were generated by crossing the same backcrossed driver lines to the corresponding *w^1118^* background strain underlying the RNAi libraries, ensuring that experimental and control animals were matched for genetic background.

### Gut dissections and immunostaining

Mated and virgin males and females of *voilà>GFP*, *AstC>GFP*, and *Tk>GFP* were dissected in cold PBS. Intact guts ut, were removed and fixed in 4% formaldehyde (Sigma) in PBS for 1 h at room temperature with gentle oscillation in shallow-welled glass dishes. Samples were washed three times for 15 min each in PBS containing 0.1% Triton X-100 (PBST) with gentle oscillation and blocked for 30 min in PBST with 5% goat serum. Tissues were incubated overnight at 4 °C with mouse anti-GFP (clone 3E6, ThermoFisher item A11120, diluted 1:500 in PBST with 5% goat serum) in sealed humidified chambers. After three washing steps (20 min each) with PBST, samples were incubated overnight at 4 °C with Alexa Fluor 488-conjugated goat anti-mouse IgG (cross-adsorbed, ThermoFisher #32723, 1:500 in PBST). Tissues were washed for three times in PBST (20 min each wash), then rinsed withPBS, and guts were finely mounted on poly-lysine–coated slides with a 120-µm Secure Seal imaging spacer (Grace Bio-Labs, #654006) in PBS. Samples were straightened carefully to facilitate imaging. PBS was replaced with ProLong Glass antifade medium containing NucBlue nuclear stain (ThermoFisher #P36985), and samples were sealed with coverslips and cured for ≥24 h at 4 °C before imaging.

### Confocal imaging

Slides were imaged on a DragonFly spinning-disk confocal microscope using a 63x glycerol-immersion objective (NA 1.3; 200 nm/pixel), controlled by Fusion software. Each gut was imaged in ∼35 overlapping tiles along the long axis and ∼5 tiles across the short axis, spanning ∼125 z-slices at 1.6 µm intervals. DAPI and Alexa Fluor 488 channels were acquired. Image volumes were assembled using the Fusion stitching function.

### Image processing and quantitative analysis

Stitched image volumes were maximum-projected using a custom MATLAB pipeline (Supplementary Code 1). Quantification was performed using a second MATLAB package (Supplementary Code 2). For each midgut, anterior and posterior boundaries (cardia and hindgut junction) were marked manually, and gut length was calculated as the pixel distance between the defined limits. NucBlue^+^ nuclei and GFP^+^ EECs were segmented, with limited pre-processing to optimize segmentation. The spatial position of each feature was recorded. Bounded guts were computationally divided into ten equal-length deciles, and feature counts were grouped accordingly. “Total cells” represent all segmented DAPI^+^ nuclei across the midgut; “GFP^+^ cells” are EECs marked by each driver. Expected female cell counts for each decile were computed by scaling male decile averages by the female-to-male gut length ratio (∼1.4 across genotypes). Ratios of observed values to these expected values in females were plotted (Fig. 1f) to identify sex-dependent regional enrichment or depletion of EECs.

### Systematic RNAi screen

Candidate genes were selected from transcriptomic and proteomic resources, including FlyAtlas2^40^, FlyGutSeq^37^, and publicly available RNA-seq datasets, based on enrichment in the adult midgut and EECs, predicted extracellular localization (Gene Ontology and protein-sequence features), or prior evidence for hormonal signaling or nutrient regulation^41^. This process yielded 165 target genes encoding secreted factors, peptide-processing enzymes, receptors (including orphan GPCRs), and nutrient or metabolite transporters. The full list of screened RNAi lines is provided in Table S4. Crosses were maintained at 18 °C, and progeny were shifted to 29 °C on an adult-optimized diet at eclosion. Adult-specific RNAi was induced for 5 days before phenotypic assays were performed, with flies kept at 29 °C throughout the experiments. Stress assays were conducted using three biological replicates of 20 animals per condition, metabolite measurements were performed with four replicates per condition, feeding behavior was assessed using 12 animals per genotype, and sleep-wake behavior was analyzed using 32 animals per genotype, with one day on the fed diet followed by one day of starvation.

### Stress-resistance assays

For starvation assays, groups of 20 flies were transferred to vials containing 1% agar in water. For ionic stress, flies were maintained on adult diet supplemented with 4% NaCl. Oxidative stress was induced using 5% H_2_O_2_ in adult diet. Survival was recorded at 6–8 h intervals until all animals perished. Three independent biological replicates (20 flies per replicate/vial) were performed for each condition.

### Metabolite measurements

Whole-body triacylglyceride (TAG), glycogen, and glucose levels were quantified from adult flies. For each biological replicate, 2–4 flies (depending on sex and driver genotype) were homogenized in PBS containing 0.5% Tween-20 (Sigma #1379) using a TissueLyser LT bead mill (Qiagen) with 5 mm stainless-steel beads. Total protein concentration was determined from homogenate aliquots using a bicinchoninic acid (BCA) assay (Sigma #B9643, #C2284, #P0914). In the large-scale screen, all metabolite measurements (TAG, glycogen, and glucose) were normalized to protein content to minimize technical variability from homogenization and sample handling. However, because females consistently contain substantially more protein than males, protein-based normalization can obscure biologically meaningful sex differences. For follow-up validation experiments, values were therefore normalized to fly number, an approach that more accurately reflects the magnitude of sex-specific differences and is widely used in studies that compare male and female TAG levels^4,6,13^. Glycogen was assayed by enzymatic hydrolysis with amyloglucosidase (Sigma #A7420) to release glucose, which was quantified using a colorimetric glucose assay kit (Sigma #GAGO20). Whole-body glucose levels were measured directly with that latter kit. TAG was quantified by enzymatic cleavage with triglyceride reagent (Sigma #T2449) to release glycerol, followed by detection with Free Glycerol Reagent (Sigma #F6428). Absorbance of indicator compound was read at 540 nm on an Ensight multimode plate reader (PerkinElmer). Protein, TAG, and glycogen concentrations were calculated from BSA, glycerol, and glucose standard curves. For starved-state measurements, males were starved for 15 h and females for 24 h on 1% agar, a difference reflecting the greater energy reserves and starvation resistance of females. For fed-state measurements, flies were collected at the same time as starved flies.

### Quantitative RT-PCR

For each genotype, five to six biological replicates were prepared, with each replicate consisting of five animals. Total RNA was extracted using the NucleoSpin RNA kit (Macherey-Nagel, #740955). Tissues were placed in 2-mL Eppendorf tubes containing RA1 lysis buffer supplemented with 1% β-mercaptoethanol and homogenized using a TissueLyser LT bead mill (Qiagen) with 5 mm stainless-steel beads (Qiagen #69989). RNA was reverse-transcribed into cDNA using the High-Capacity cDNA Reverse Transcription Kit (Applied Biosystems, #4368814) following the manufacturer’s instructions. Quantitative PCR was performed using RealQ Plus 2× SYBR Green Master Mix (Ampliqon, #A324402) on a QuantStudio 5 real-time PCR system (Applied Biosystems). Gene expression values were normalized to the ribosomal reference gene *Rp49*. Primer sequences are provided in Table S5.

### Feeding assays

Food choice was assessed using a two-choice dye-based feeding assay^8,9^. Groups of 25 flies were briefly anaesthetized with CO_2_ and placed into 35-mm Petri dishes containing 20-µL diet spots arranged in a checkerboard pattern. The two diets consisted of either 100 g/L sucrose (dyed with 0.5% amaranth; Sigma #A1016) or 100 g/L yeast extract (dyed with 0.5% erioglaucine; Sigma #861146). Animals were allowed to feed for 2 hours in the dark. After feeding, flies were collected in groups of 1–2, homogenized in 100 µL phosphate buffer (pH 7.5) using a TissueLyser LT bead mill (Qiagen) with 5-mm stainless-steel beads, and centrifuged at 16,000 *g* for 5 minutes. Fifty microliters of cleared lysate were transferred to 384-well plates, and absorbance was measured at 520 nm (amaranth) and 629 nm (erioglaucine) using an Ensight multimode plate reader (PerkinElmer). Standard curves generated for each dye were used to calculate animals’ consumption of sucrose and yeast diets. Feeding behavior was also monitored using the Fly Liquid-Food Interaction Counter (FLIC; Sable Systems, USA)^82^. Flies were housed individually in chambers of the *Drosophila* Feeding Monitor, maintained at 29 °C and 70% relative humidity under a 12 h:12 h light–dark cycle. Feeding wells contained 10% sucrose solution, and flies were introduced in the afternoon to allow acclimation. Feeding interactions were recorded overnight and during the following morning meal, beginning at lights-on. Data were acquired using the manufacturer’s software and processed in *R* with the published analysis package (https://github.com/PletcherLab/FLIC_R_Code).

### Sleep and activity assays

Sleep and locomotor behavior were measured using the Drosophila Activity Monitoring System (DAMS; TriKinetics, Waltham, MA, USA). Flies aged 5 days at 29 °C were lightly anaesthetized with CO_2_ and loaded individually into glass monitoring tubes. Each tube was closed at one end with a foam plug and at the other with a removable 250-µL PCR tube containing 90 µL of food medium. The food consisted of either 5% or 0% sucrose in 1% agarose, supplemented with 0.5% propionic acid and 0.15% methyl-4-hydroxybenzoate. Recordings began at lights-on of a 12 h:12 h light–dark cycle, with 5%-sucrose end tubes. After 24 h, these were swapped out for replacements containing 1% agar for starvation assays, and recordings continued under the same light–dark conditions, all at 29 °C. Periods of at least 5 min without beam crossings were scored as sleep. Data were processed with a custom MATLAB pipeline (MathWorks, Natick, MA) to quantify total sleep, daytime and nighttime sleep, locomotor activity, and starvation-induced changes in behavior.

### Statistical analysis

All statistical analyses were performed in MATLAB (The MathWorks), RStudio (FLIC analyses), or the Prism package (GraphPad, Inc., for general statistics and chart creation). For comparisons of two groups, unpaired two-sided *t*-tests were used; if data were not normally distributed, non-parametric Mann–Whitney *U* tests were applied instead. For comparisons involving more than two groups, ANOVA was performed followed by either Holm–Šidák or Tukey’s *post-hoc* tests for multiple comparisons. When normality assumptions were not met, non-parametric Kruskal–Wallis tests were used. For screen-wide hit calling, phenotypes were converted to z-scores relative to sex-matched controls, and hits were defined as values outside ±1.5 s.d. (z-score ±1.5), except for oxidative-stress survival and nighttime sleep, where a relaxed cutoff of ±2 s.d. (z-score ±2) was used due to smaller trait variance. This approach was chosen instead of multiple-testing correction to allow uniform identification of extreme phenotypes across assays. For correlation analyses, slopes, coefficients of determination (*R^2^*), and *p*-values are reported from linear regression fits, with shaded bands indicating 95% confidence intervals. For hierarchical clustering of phenotypic relationships (Fig. 4), phenotypic differences with sex-matched controls were converted into z-scores and clustered using Spearman’s rank correlation with average linkage. To assess sex differences in trait magnitude (Fig. 5), clustering was instead performed on pooled male and female absolute phenotypic values. For principal-component analysis (PCA), z-score–normalized phenotypic values were used with singular value decomposition; distances between groups were calculated as Euclidean distances in PCA space. Significance thresholds and the statistical tests used for each figure are specified in the figure legends. The MATLAB code used for the initial processing of the images used for Figure 1 is included in Supplemental Code 1, and the code underlying the further quantification and analysis in that figure is included in Supplemental Code 2. The MATLAB scripts used for screen-wide analysis, hierarchical clustering, generation of Venn diagrams and heatmaps, principal-component analyses, and pairwise correlation testing are included in Supplemental Code 3, along with the necessary raw data.

## Data Availability

All data supporting the findings of this study are included in the article and its supplementary materials. Additional raw data underlying the results will be made freely available by the corresponding author (K.R.) upon reasonable request.

## Code Availability

All custom code developed for this study is provided with the paper. This includes Code 1 (custom Imaris viewer/manipulator), Code 2 (gut-length measurement and cell-counting pipeline), and Code 3 (scripts for image processing, clustering, statistical analysis, and figure generation).

## Acknowledgements

We greatly appreciate sharing data and insights with Elizabeth Rideout (University of British Columbia). We are grateful to the Bloomington and Vienna stock centers and to their respective funding agencies. This work was supported by the Novo Nordisk Foundation (grant NNF19OC0054632) and the Lundbeck Foundation (grant 2019-772) to K.R. The spinning-disk confocal microscope and the PerkinElmer EnSight plate reader were purchased with generous grants from the Carlsberg Foundation (grants CF23-1302 and CF17-0615, respectively) to K.R.

## Author contributions

O.K., A.M., S.N., and K.R. conceived and designed the RNAi screen. O.K., A.M., N.A., and S.N. performed the screen; O.K., A.M., and N.A. carried out the follow-up experiments. O.K., A.M., N.A., and S.N. analyzed the data, with additional computational analyses and MATLAB-based pipelines developed and implemented by M.J.T. O.K., A.M., N.A., and M.J.T. prepared figures. M.J.T. contributed to data interpretation and helped write the manuscript. K.R. supervised the project and wrote the manuscript.

## Supplemental Figures

**Figure S1.**
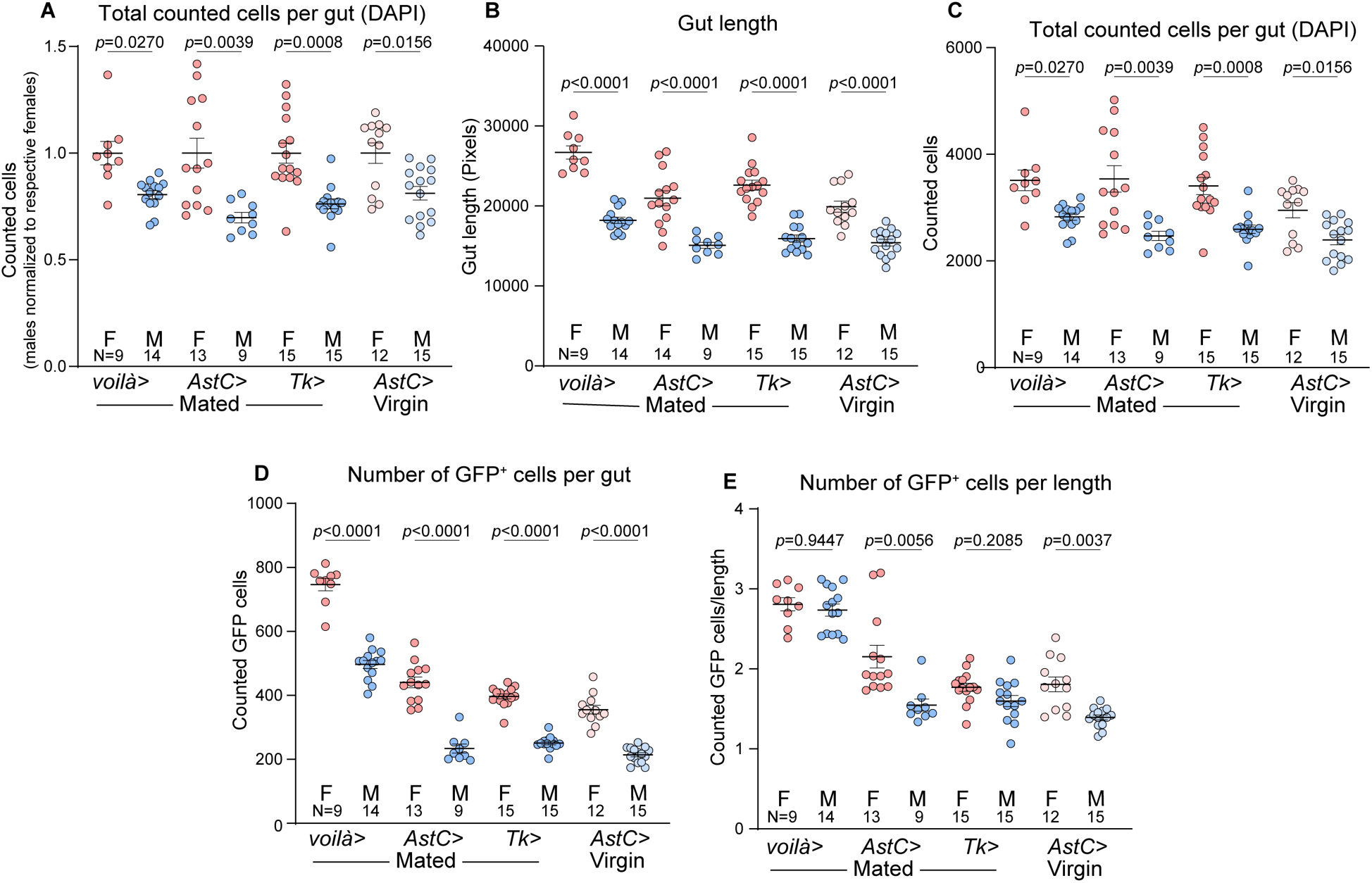
Gut length, total cell number, and EEC counts. (a) The total number of cells of all kinds per midgut (detected with DAPI staining). Values for each sex and genotype are normalized to the female average. (b) Absolute midgut length (measured in pixels in the confocal image) in males (M) and females (F) expressing GFP in defined enteroendocrine-cell (EEC) populations using *voilà-GAL4*, *AstC::T2A::GAL4* (*AstC>*), or *Tk::T2A::GAL4* (*Tk>*) drivers, or *AstC>* in virgins. (c) Total number of cells of all kinds per midgut (DAPI-stained nuclei). (d) Number of GFP-positive EECs per gut. (e) Number of GFP-positive EECs per unit length, shown without normalization to the female reference values. Data are presented as mean ± s.e.m. with individual data points. Statistical comparisons between sexes are indicated. Statistics: a-d: Welch’s ANOVA with Dunnett’s T3 multiple comparisons.

**Figure S2.**
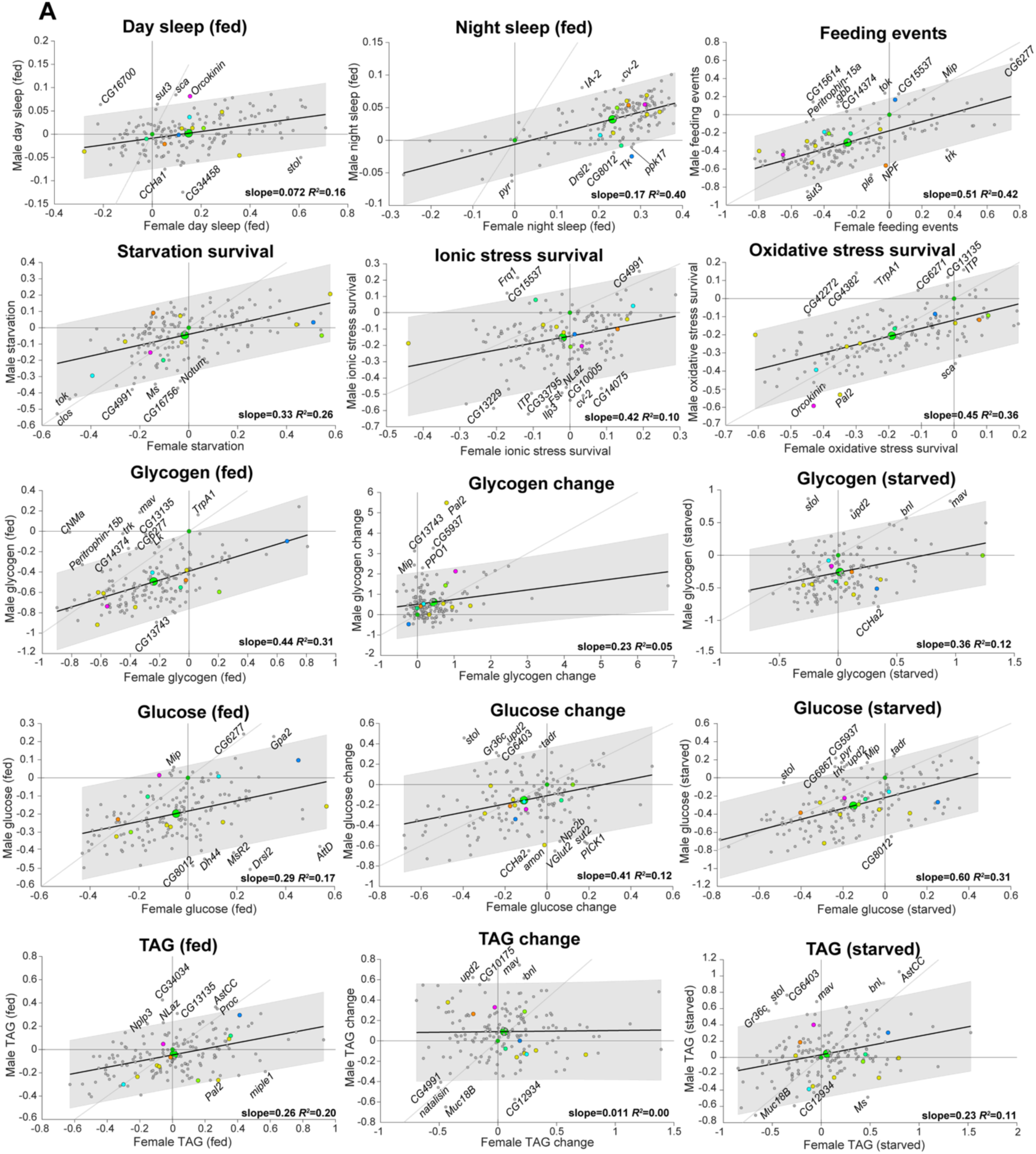
Comparative analysis of male and female responses across all physiological and behavioral assays. (a) Plots of each knockdown genotype’s effects in females (X axis) versus in males (Y axis). Each point represents an individual gene knockdown, with a fitted trend line and 95% confidence intervals shown in gray. The control genotype falls at position (0, 0) by definition, and the mean of all genotypes is indicated as a larger green dot. Points along the diagonal represent knockdowns with similar effects in both sexes, and distance from the diagonal indicates the degree of sex bias in effect size. Points in opposite quadrants, where the effect is positive in one sex but negative in the other, represent strong discordance, in which a given knockdown elicits opposing phenotypes in males and females. In the metabolite-loss charts, positive values indicate greater depletion during starvation than in the control, 0.0 indicates losses equal to those of the control, and negative values indicate smaller losses than in the control; a score of –1.0 indicates no loss during starvation (100% less than in the control).

**Figure S3.**
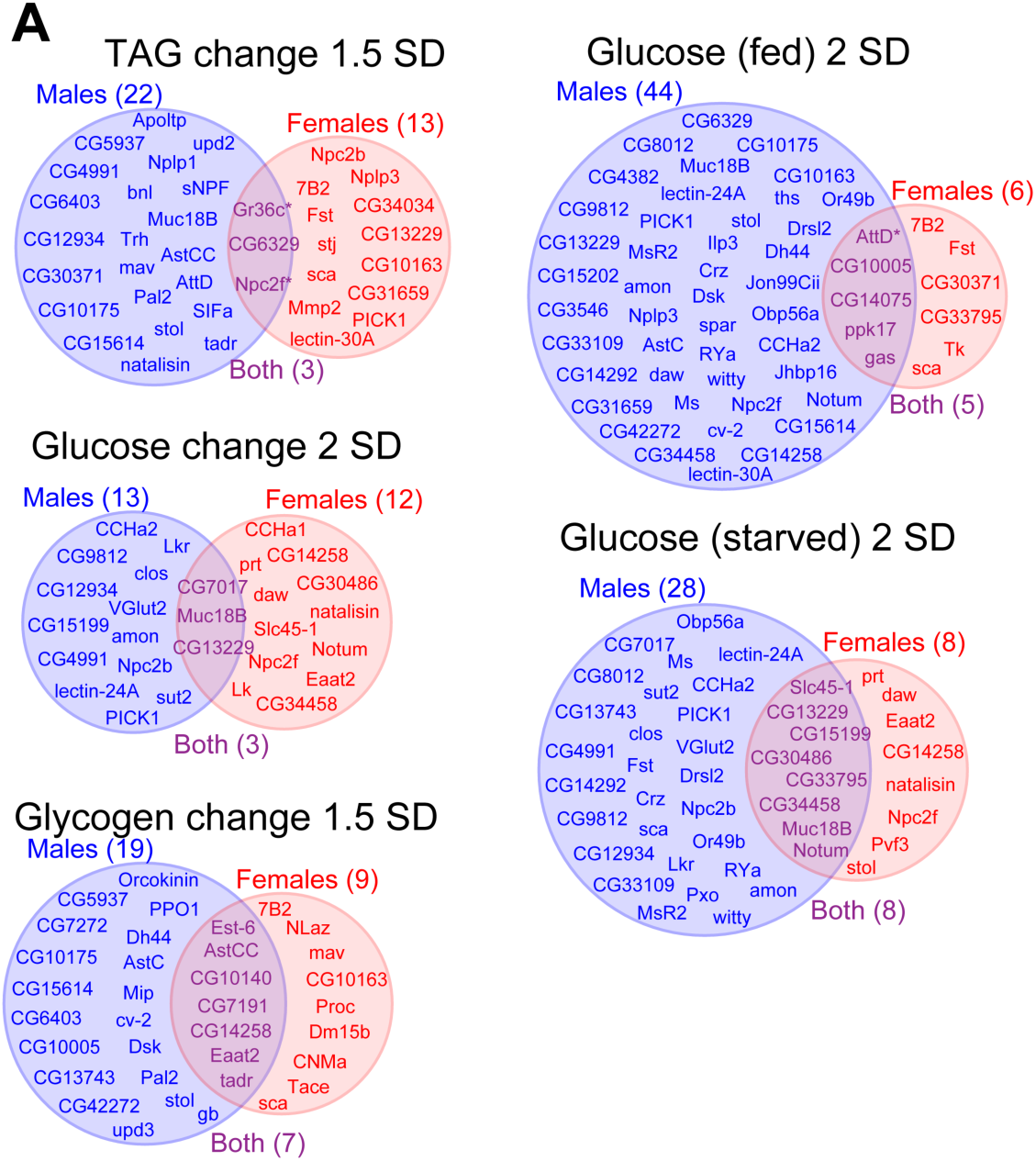
Sex-specific gut hormone knockdown phenotypes in additional metabolic assays. Hits were defined as EEC gene knockdowns with phenotypic values outside ±1.5 standard deviations (SD; z-score ±1.5) from the control distribution (±2 SD; z-score ±2 for glucose-related assays). This definition was applied consistently across all panels. (a) Venn diagrams show the distribution of hits into male-specific (blue, left), female-specific (red, right), and shared (purple, overlap) categories. Panels display hits for TAG loss during starvation, glucose levels in fed and starved animals, and glucose and glycogen loss during starvation.

**Figure S4.**
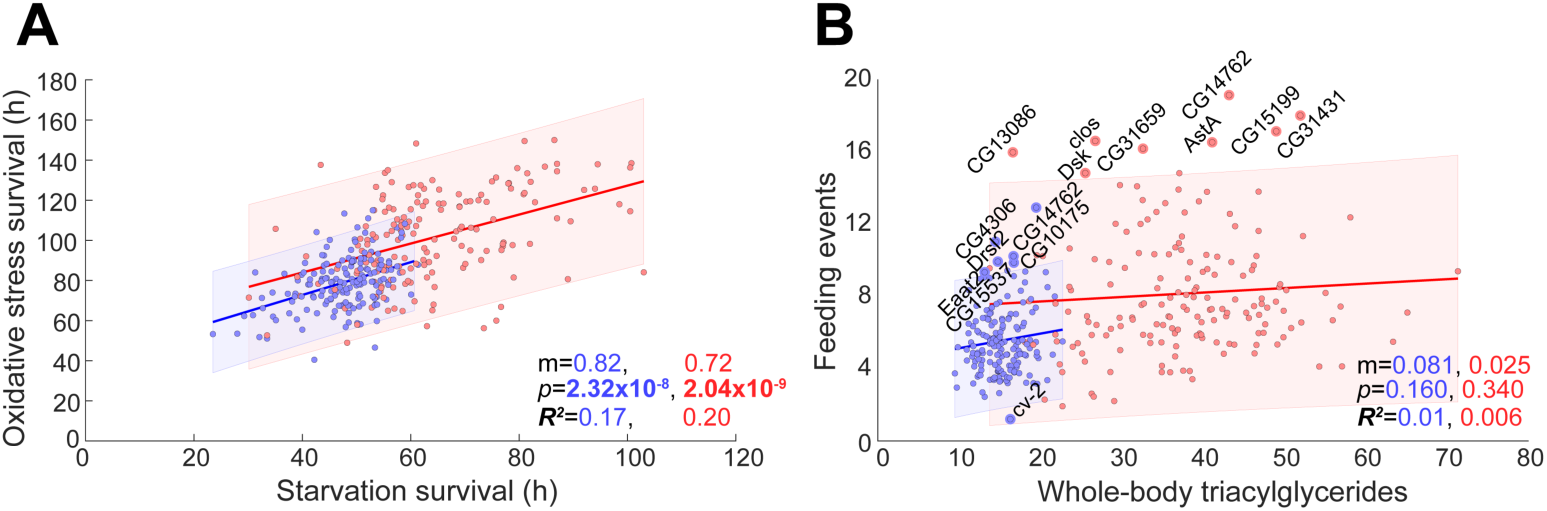
Additional trait correlations involving survival and feeding. (a) Starvation-survival duration plotted against oxidative-stress survival time across EEC-specific knockdown lines in females (red) and males (blue). (b) Whole-body triacylglyceride (TAG) levels plotted against number of feeding events. Each dot represents an individual knockdown line. Trend lines are shown with shaded 95% confidence intervals. Knockdowns falling outside this region are labeled. Reported values indicate each fit’s slope (m), p-value for “no correlation,” and coefficient of determination (*R^2^*) for males (blue, left) and females (red, right).

**Figure S5.**
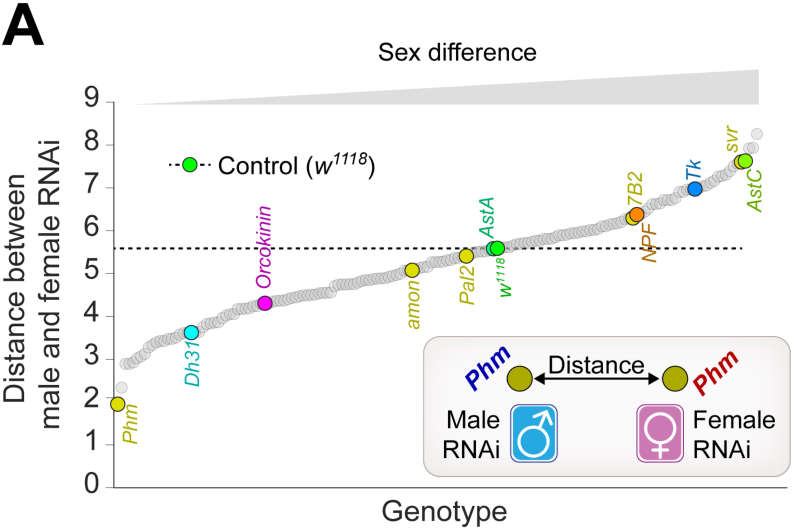
Sex differences in phenotypic space across EEC knockdowns. (a) Euclidean distance in the PCA1 x PCA2 plane between male and female RNAi animals of the same genotype. Genotypes are ordered along the x-axis, with the control (*voilà*> x *w^1118^*) indicated in green. Selected knockdowns are labeled. The schematic inset illustrates the distance calculation between male and female RNAi pairs. Statistics: Phenotypic values were converted into Z-scores, and PCA was performed with centering and the singular value decomposition method.

**Figure S6.**
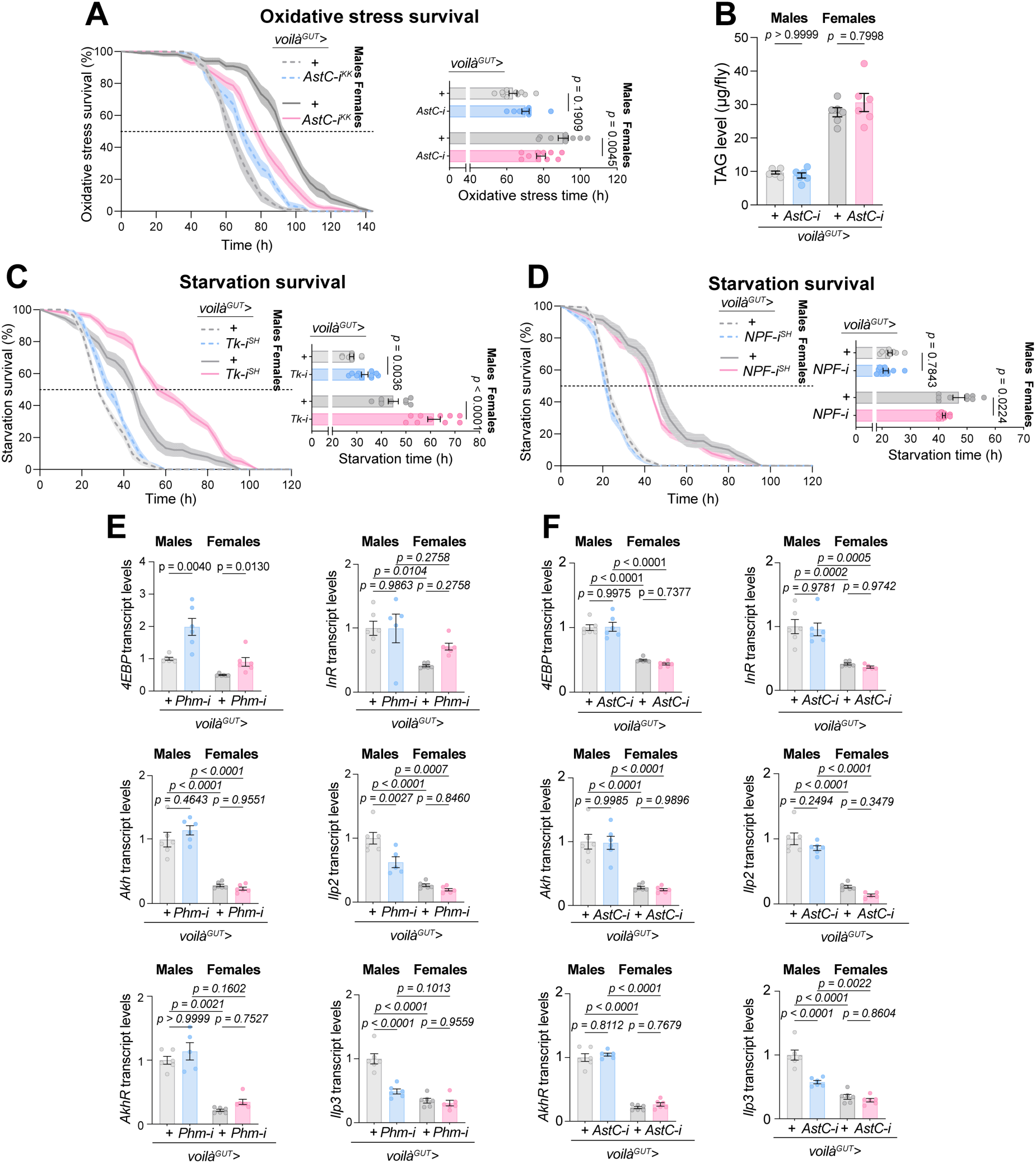
Additional effects of EEC-derived peptide hormone knockdown on stress survival, lipid storage, and systemic signaling. (a) Survival under oxidative stress upon EEC-specific knockdown of *AstC*. (b) Triacylglyceride (TAG) levels in males and females with EEC-specific *AstC* knockdown. (c,d) Survival under starvation upon EEC-specific knockdown of *Tk* (c) or *NPF* (d). (e, f) Expression of insulin- and AKH-pathway components (*Ilp2*, *Ilp3*, *Akh*, *AkhR*, *InR*, *4EBP*) in males and females upon EEC-specific knockdown of *Phm* (e) or *AstC* (f). Boxplots show median and interquartile range with individual data points; bar graphs show mean ± s.e.m. Statistical tests: survival assays (a, c, d) were analyzed by one-way ANOVA with Tukey’s post-hoc tests; TAG levels (b) by unpaired two-sided *t*-test; and gene-expression assays (e, f) by one-way ANOVA with Tukey’s post hoc tests or unpaired two-sided *t*-test (pairwise comparisons).

