## Supplemental code 1 for "Gut hormone signaling drives sex differences in metabolism and behavior": filedoer.m

```
clear all
fclose all
close all
warning off
global addnewline
addnewline=true;
global notelines
notelines=0;
global fourchannel
fourchannel=false;
global finished1
finished1=false;
global comboch1
comboch1=0;
global comboch2
comboch2=1;
global comboch3
comboch3=3;
global origbuttoncolor;
global kured;
global filesep
filesep='\';
if(ismac)
    filesep='/';
end
global skipexistingbutton
global extension
extension='.ims';
global filenum2
filenum2=0;
global interrupt
interrupt=false;
global maxfilechecks
maxfilechecks=100000000;
global maxfiles
maxfiles=20000;
global depth;
depth=0;
global doanything
doanything=false;
global filelist;
filelist={};
global dirlist;
dirlist={};
global filenum;
filenum=0;
global imsnum
imsnum=0;
global fulllist
fulllist={};
global tifflistbox
global tifflistfiles
tifflistfiles={};

global todofulllist
todofulllist={};
%global datasetnames
%global datasetnames2
global uif;
global combonames
global combinations
global combonames
global numtoprocess
numtoprocess=0;
global currentprocess
currentprocess=0;
global allfileslist
global realnames;
global numcombos
global combinations
global tifflistfiles
tifflistfiles={};
global combonames;
global includedfileslist;
global gogo
global uif;
global giantbutton
giantbutton={};
global grid
global blockedalready;
blockedalready=false;
gogo=false;
numcombos=0;
global vname
global vnames
global vloc
global numfiles
global realnames;
global master;
global datasetNs;
global vnamenoext;
global dataset
realnames={};
master=cell(1,1);
datasetNs=zeros(1,1);
includedfileslist={};
combonames={};
combinations={};
global quit
quit=false;
global hasstitchedfiles;
hasstitchedfiles=false;
warning off
scrdims=get(0,'ScreenSize');
scrheight=scrdims(4);
scrwidth=scrdims(3);
global prefix
prefix="";
colors={[0 0.5 0.5] [1 0 0] [0 1 0] [0 0 1] [0.9 0.9 0] [1 0 1] [0 .9 .9] [1 0.5 0] [0.5 0 1] };
winwidth=min(scrwidth,3100);
winheight=min(scrheight,800);
left=scrwidth/2-winwidth/2;
top=scrheight/2-winheight/2;
uif=uifigure;
uif.Name='Imaris reader and image processor';
grid=uigridlayout(uif);
grid.BackgroundColor='#999999';
%                1   2   3   4   5   6  7   8   9   10   11  12  13  14  15  16
grid.RowHeight={20, 40, 40, 40, 40, 40, 40, 40, 40, 40, 40, 40, 40, 40, 40, 40};
uif.Position=[left top winwidth+108 winheight-10];
diff=400;
grid.ColumnWidth={(winwidth-diff)/6,(winwidth-diff)/6,100,(winwidth-diff)/6,(winwidth-diff)/6,100,(winwidth-diff)/6,(winwidth-diff)/6,100,100};
allfileslist=uilistbox(grid,"Multiselect","on","Items",convertCharsToStrings(realnames));
allfileslist.Layout.Row=[2 16];
allfileslist.Layout.Column=[1 2];

allfileslabel=uilabel(grid);
allfileslabel.Layout.Row=1;
allfileslabel.Layout.Column=1;
allfileslabel.Text="Imaris files to choose from";

buttonLoad=uibutton(grid,'Text',"Add folder",'ButtonPushedFcn',@(src,event)buttonLoadevent(src,event));
buttonLoad.Tag='#f5f5f5';
buttonLoad.Layout.Row=2;
buttonLoad.Layout.Column=3;

kugray="#666666";
kured="#901A1E";
grid.BackgroundColor=kugray;

buttonLoad2=uibutton(grid,'Text',"Add files",'ButtonPushedFcn',@(src,event)buttonLoad2event(src,event));
buttonLoad2.Tag='#f5f5f5';
buttonLoad2.Layout.Row=3;
buttonLoad2.Layout.Column=3;

%
% 
% 
% 

buttonAdd=uibutton(grid,'Text',"Grab selected -->",'ButtonPushedFcn',@(src,event)buttonAddevent(src,event));
buttonAdd.Tag='#f5f5f5';
buttonAdd.Layout.Row=5;
buttonAdd.Layout.Column=3;

buttonAll=uibutton(grid,'Text',"Grab all -->",'ButtonPushedFcn',@(src,event)buttonAllevent(src,event));
buttonAll.Tag='#f5f5f5';
buttonAll.Layout.Row=6;
buttonAll.Layout.Column=3;

buttonStitched=uibutton(grid,'Text',"Grab stitched -->",'ButtonPushedFcn',@(src,event)buttonStitchedevent(src,event));
buttonStitched.Tag='#f5f5f5';
buttonStitched.Layout.Row=7;
buttonStitched.Layout.Column=3;

%
% 
% 
% 

buttonRemove=uibutton(grid,'Text',"<-- Remove",'ButtonPushedFcn',@(src,event)buttonRemoveevent(src,event));
buttonRemove.Tag='#f5f5f5';
buttonRemove.Layout.Row=8;
buttonRemove.Layout.Column=3;

selectedfileslabel=uilabel(grid);
selectedfileslabel.Layout.Row=1;
selectedfileslabel.Layout.Column=4;
selectedfileslabel.Text="Selected Imaris files to process";


includedfileslist=uilistbox(grid,"Multiselect","off","Items",convertCharsToStrings(combonames));
includedfileslist.Layout.Row=[2 16];
includedfileslist.Layout.Column=[4 5];

buttonProcess=uibutton(grid,'Text',"Process selected -->",'ButtonPushedFcn',@(src,event)buttonProcessevent(src,event));
buttonProcess.Tag='#f5f5f5';
buttonProcess.Layout.Row=2;
buttonProcess.Layout.Column=6;

buttonProcessAll=uibutton(grid,'Text',"Process all -->",'ButtonPushedFcn',@(src,event)buttonProcessAllevent(src,event));
buttonProcessAll.Tag='#f5f5f5';
buttonProcessAll.Layout.Row=3;
buttonProcessAll.Layout.Column=6;

buttonProperties=uibutton(grid,'Text',"Properties",'ButtonPushedFcn',@(src,event)buttonPropertiesevent(src,event));
buttonProperties.Tag='#f5f5f5';
buttonProperties.Layout.Row=4;
buttonProperties.Layout.Column=6;

buttonChannels=uibutton(grid,'Text',sprintf("Can't output\n4-channel TIFFs.\n\nArrange output\nchannels for\n4-channel files\n(%i, %i, %i)",comboch1,comboch2,comboch3),'ButtonPushedFcn',@(src,event)fourchannelsevent(src,event));
buttonChannels.Tag='#f5f5f5';
buttonChannels.Layout.Row=[5 7];
buttonChannels.Layout.Column=6;

tifffileslabel=uilabel(grid);
tifffileslabel.Layout.Row=1;
tifffileslabel.Layout.Column=7;
tifffileslabel.Text="TIFF files";

tifflistbox=uilistbox(grid,"Multiselect","on","Items",convertCharsToStrings(tifflistfiles));
tifflistbox.Layout.Row=[2 16];
tifflistbox.Layout.Column=[7 8];

buttonAddTiffFolder=uibutton(grid,'Text',"Add TIFF folder",'ButtonPushedFcn',@(src,event)buttonLoadTiffFolderevent(src,event));
buttonAddTiffFolder.Tag='#f5f5f5';
buttonAddTiffFolder.Layout.Row=2;
%buttonAddTiffFolder.Enable=false;
buttonAddTiffFolder.Layout.Column=9;

buttonAddTiffFiles=uibutton(grid,'Text',"Add TIFF files",'ButtonPushedFcn',@(src,event)buttonAddTiffFilesevent(src,event));
buttonAddTiffFiles.Tag='#f5f5f5';
buttonAddTiffFiles.Layout.Row=3;
buttonAddTiffFiles.Layout.Column=9;

buttonShow=uibutton(grid,'Text',"Show as is",'ButtonPushedFcn',@(src,event)buttonShowevent(src,event));
buttonShow.Tag='#f5f5f5';
buttonShow.Layout.Row=4;
buttonShow.Layout.Column=9;

buttonShow2=uibutton(grid,'Text',"Show bright",'ButtonPushedFcn',@(src,event)buttonShow2event(src,event));
buttonShow2.Tag='#f5f5f5';
buttonShow2.Layout.Row=5;
buttonShow2.Layout.Column=9;

buttonBrightenAll=uibutton(grid,'Text',sprintf("Brighten\nproportionally"),'ButtonPushedFcn',@(src,event)buttonBrightenAllevent(src,event));
buttonBrightenAll.Tag='#f5f5f5';
buttonBrightenAll.Layout.Row=6;
buttonBrightenAll.Layout.Column=9;
buttonBrightenAll.Enable=false;

buttonBrightenAll2=uibutton(grid,'Text',sprintf("Brighten\nindividually"),'ButtonPushedFcn',@(src,event)buttonBrightenAll2event(src,event));
buttonBrightenAll2.Tag='#f5f5f5';
buttonBrightenAll2.Layout.Row=7;
buttonBrightenAll2.Layout.Column=9;

buttonRename=uibutton(grid,'Text',"Rename",'ButtonPushedFcn',@(src,event)buttonRenameevent(src,event));
buttonRename.Tag='#f5f5f5';
buttonRename.Layout.Row=8;
buttonRename.Layout.Column=9;

buttonShrink=uibutton(grid,'Text',sprintf("Shrink"),'ButtonPushedFcn',@(src,event)buttonShrinkevent(src,event));
buttonShrink.Tag='#f5f5f5';
buttonShrink.Layout.Row=9;
buttonShrink.Layout.Column=9;

buttonFlipLR=uibutton(grid,'Text',sprintf("Flip L<>R"),'ButtonPushedFcn',@(src,event)buttonFlipLRevent(src,event));
buttonFlipLR.Tag='#f5f5f5';
buttonFlipLR.Layout.Row=10;
buttonFlipLR.Layout.Column=9;


buttonRechannel=uibutton(grid,'Text',sprintf("Rearrange\nchannels"),'ButtonPushedFcn',@(src,event)buttonRechannelevent(src,event));
buttonRechannel.Tag='#f5f5f5';
buttonRechannel.Layout.Row=11;
buttonRechannel.Layout.Column=9;


buttonPNG=uibutton(grid,'Text',sprintf("Save PNG"),'ButtonPushedFcn',@(src,event)buttonPNGevent(src,event));
buttonPNG.Tag='#f5f5f5';
buttonPNG.Layout.Row=12;
buttonPNG.Layout.Column=9;

buttonRemoveTIFF=uibutton(grid,'Text',sprintf("Remove\nfrom list"),'ButtonPushedFcn',@(src,event)buttonRemoveTIFFevent(src,event));
buttonRemoveTIFF.Tag='#f5f5f5';
buttonRemoveTIFF.Layout.Row=13;
buttonRemoveTIFF.Layout.Column=9;

buttonDeleteTIFF=uibutton(grid,'Text',sprintf("DELETE"),'ButtonPushedFcn',@(src,event)buttonDeleteTIFFevent(src,event));
buttonDeleteTIFF.Tag='#f5f5f5';
buttonDeleteTIFF.Layout.Row=13;
buttonDeleteTIFF.Layout.Column=10;

buttonSelectTIFFs=uibutton(grid,'Text','Select...','ButtonPushedFcn',@(src,event)buttonSelectTIFFsevent(src,event));
buttonSelectTIFFs.Tag='#f5f5f5';
buttonSelectTIFFs.Layout.Row=14;
buttonSelectTIFFs.Layout.Column=9;

buttonInvertTIFFs=uibutton(grid,'Text',sprintf("Invert\nselection"),'ButtonPushedFcn',@(src,event)buttonInvertTIFFsevent(src,event));
buttonInvertTIFFs.Tag='#f5f5f5';
buttonInvertTIFFs.Layout.Row=15;
buttonInvertTIFFs.Layout.Column=9;

buttonConsolidate=uibutton(grid,'Text',sprintf("Copy into\none folder"),'ButtonPushedFcn',@(src,event)buttonConsolidateevent(src,event));
buttonConsolidate.Tag='#f5f5f5';
buttonConsolidate.Layout.Row=5;
buttonConsolidate.Layout.Column=10;


buttonQuit=uibutton(grid,'Text',"Quit",'ButtonPushedFcn',@(src,event)buttonQuitevent(src,event));
buttonQuit.Tag='#f5f5f5';
buttonQuit.Layout.Row=16;
buttonQuit.Layout.Column=9;

skipexistingbutton=uibutton(grid,'state','Text','Skip existing',"ValueChangedFcn",@(src,event)updateskip(src,event));
skipexistingbutton.Tag='#f5f5f5';
skipexistingbutton.Layout.Row=2;
skipexistingbutton.Layout.Column=10;
skipexistingbutton.Value=true;
origbuttoncolor=skipexistingbutton.BackgroundColor;
buttonon(skipexistingbutton);


while quit==false
    buttonLoad.Enable=true;
    buttonAll.Enable=false;
    buttonProcess.Enable=false;
    buttonProcessAll.Enable=false;
    buttonRemove.Enable=false;
    buttonAdd.Enable=false;
    buttonProperties.Enable=false;
    buttonStitched.Enable=hasstitchedfiles;
    buttonChannels.Text=sprintf("Can't output\n4-channel TIFFs.\n\nArrange output\nchannels for\n4-channel files\n(%i, %i, %i)",comboch1,comboch2,comboch3);

    tlb= (length(tifflistbox.Value)>0);
buttonBrightenAll.Enable=tlb;
buttonBrightenAll2.Enable=tlb;
buttonShow.Enable=tlb;
buttonShow2.Enable=tlb;
buttonFlipLR.Enable=tlb;
buttonRemoveTIFF.Enable=tlb;
buttonShrink.Enable=tlb;

buttonRechannel.Enable=tlb;
buttonPNG.Enable=tlb;
buttonConsolidate.Enable=tlb;
buttonDeleteTIFF.Enable=tlb;
buttonRename.Enable=tlb;


    if length(allfileslist.Items)>0
%        buttonLoad.Enable=false;
        buttonAll.Enable=true;
    end
    if length(allfileslist.Value)>0
        buttonAdd.Enable=true;
    end
    if(length(includedfileslist.Items)>0)
        buttonProcessAll.Enable=true;
    end
    if(length(includedfileslist.Value)>0)
        buttonProcess.Enable=true;
        buttonProperties.Enable=true;
    end
    if(size(includedfileslist.Value,1)==1)
        buttonRemove.Enable=true;
    end
    pause(0.1)

    if quit==true
        close(uif);
        return
    end
end

function updateskip(src,event)
global skipexistingbutton
if skipexistingbutton.Value
    buttonon(skipexistingbutton);
else
    buttonoff(skipexistingbutton);
end
end

function buttonProcessevent(src, event)
    global includedfileslist
    global numtoprocess
    global currentprocess
    global interrupt
    global fourchannel
    global uif

    fourchannel=false;
%    numtoprocess=size(includedfileslist.Value,2);
%    for currentprocess=1:numtoprocess
currentprocess=1;
numtoprocess=1;
%        if(interrupt)
%            interrupt=false;
%            break
%        end
%        processFile(includedfileslist.Value{currentprocess});
        processFile(includedfileslist.Value);
%        includedfileslist.Value=includedfileslist.Items{currentprocess};
        buttonRemoveevent(1,2);
%    end

if fourchannel
    fourchannel=false;
    msg=sprintf("This image had four channels.\nMatLAB can only save four-channel images as multipage TIFFs.\nTo use one in FIJI, load it, then select\nImage-->Hyperstacks-->Stack to Hyperstack...\nThen: Order=default, Channels=4, Slices=1, Frames=1, Display Mode=Color, OK");
    uiconfirm(uif,msg,"Notice");
end
    showui();
end


function buttonProcessAllevent(src, event)
    global includedfileslist
    global numtoprocess
    global currentprocess
    global interrupt
    global uif
    global fourchannel

    fourchannel=false;
    numtoprocess=size(includedfileslist.Items,2);
    currentprocess=1;
    while size(includedfileslist.Items,2)>0
        if(interrupt)
            interrupt=false;
            break
        end
        processFile(includedfileslist.Items{1});
        includedfileslist.Value=includedfileslist.Items{1};
        buttonRemoveevent(1,2);
        currentprocess=currentprocess+1;
    end
    if fourchannel
        fourchannel=false;
        msg=sprintf("One or more of these images had four channels.\nMatLAB can only save four-channel images as multipage TIFFs.\nTo use one in FIJI, load it, then select\nImage-->Hyperstacks-->Stack to Hyperstack...\nThen: Order=default, Channels=4, Slices=1, Frames=1, Display Mode=Color, OK");
        uiconfirm(uif,msg,"Notice");
    end
    showui();
end

function buttonShowevent(src,event)
    global tifflistbox
    blockui("Loading image...");
    num=length(tifflistbox.Value);
for t=1:num
    i=imread(tifflistbox.Value{t});
    %fprintf("%i %i %i",size(i,1),size(i,2),size(i,3));
    figure('name',tifflistbox.Value{t})
    imshow(i);
end
    showui();
end


function buttonShow2event(src,event)
    global tifflistbox
    num=length(tifflistbox.Value);
    tp=99.8;
    tp=str2double(inputdlg("Input percentile to set as max","Input percentile",[1 45],{num2str(tp)}));
    m=10;
    m=str2double(inputdlg("Input percentile to set as zero","Input percentile",[1 45],{num2str(m)}));
    if isnan(tp)
        tp=99.8;
    end
    if isnan(m)
        m=10;
    end

    blockui(sprintf("Normalizing brightnesses individually to %f percentile=MAX",tp));
    for t=1:num
        name=tifflistbox.Value{t};
        appendtonote(sprintf("Reading image %s",name));
        i=im2double(imread(tifflistbox.Value{t}));
        ch=size(i,3);
        if ch==1
%            zmax=max(max(i(:,:)));
            i2=reshape(i(:,:),[],1);
            mm=prctile(i2,m);
            i=i-m;
            i2=i2-m;
            zmax=prctile(i2,tp);
            i=i/zmax;
        end
        if(ch==3)
            z1=i(:,:,1);
            i2=reshape(z1(:,:),[],1);
            mm=prctile(i2,m);
            z1=z1-mm;
            i2=i2-mm;
            z1max=prctile(i2,tp);
            z1=z1/z1max;
 
            z2=i(:,:,2);
            i2=reshape(z2(:,:),[],1);
            mm=prctile(i2,m);
            z2=z2-mm;
            i2=i2-mm;
            z2max=prctile(i2,tp);
            z2=z2/z2max;

            z3=i(:,:,3);
            i2=reshape(z3(:,:),[],1);
            mm=prctile(i2,m);
            z3=z3-mm;
            i2=i2-mm;
            z3max=prctile(i2,tp);
            z3=z3/z3max;

            i=cat(3,z1,z2,z3);
        end
        if(ch==2)
            z1=i(:,:,1);
            i2=reshape(z1(:,:),[],1);
            mm=prctile(i2,m);
            z1=z1-m;
            i2=i2-m;
            z1max=prctile(i2,tp);
            z1=z1/z1max;

            z2=i(:,:,2);
            i2=reshape(z2(:,:),[],1);
            mm=prctile(i2,m);
            z2=z2-m;
            i2=i2-m;
            z2max=prctile(i2,tp);
            z2=z2/z2max;

            i=cat(2,z1,z2);
        end
    figure('name',strcat(tifflistbox.Value{t},' brightened (',num2str(m),'-',num2str(tp),')'));
    imshow(i);
    end
showui();
end


function buttonAllevent(src,event)
    global allfileslist
    nn=length(allfileslist.Items);
     for t=1:nn
         allfileslist.Value=allfileslist.Items{1};
         buttonAddevent(src,event)
     end
end

function buttonStitchedevent(src,event)
    global allfileslist
    proceed=true;
    t=0;
    while proceed==true
        t=t+1;
        if(t>length(allfileslist.Items))
            proceed=false;
            break
        end
        x=strfind(allfileslist.Items{t},'FusionStitcher');
        if size(x,1)>0
            allfileslist.Value=allfileslist.Items{t};
            buttonAddevent(src,event)
            t=0;
        end
    end 
end

function buttonRemoveevent(src,event)
    global allfileslist
    global includedfileslist

    num=size(includedfileslist.Value,2);
    newitems={};
    allfileslist.Items{length(allfileslist.Items)+1}=includedfileslist.Value;
    n=0;
    for k=1:length(includedfileslist.Items)
        remove=false;
%        for v=1:num
%            if(strcmp(includedfileslist.Value{v},includedfileslist.Items{k}))
            if(strcmp(includedfileslist.Value,includedfileslist.Items{k}))
                remove=true;
            end
%        end
        if(remove==false)   
            n=n+1;
            newitems{n}=includedfileslist.Items{k};
        end
    end
    includedfileslist.Items=convertStringsToChars(newitems);
    drawnow
end

function buttonRemove2event(src,event)
    global includedfileslist
    num=size(includedfileslist.Value,2);
    newitems={};
    n=0;
    for k=1:length(includedfileslist.Items)
        remove=false;
%        for v=1:num
%            if(strcmp(includedfileslist.Value{v},includedfileslist.Items{k}))
            if(strcmp(includedfileslist.Value,includedfileslist.Items{k}))
                remove=true;
            end
%        end
        if(remove==false)   
            n=n+1;
            newitems{n}=includedfileslist.Items{k};
        end
    end
    includedfileslist.Items=convertStringsToChars(newitems);
    drawnow
end

function buttonRemoveTIFFevent(src,event)
    global tifflistbox
    fprintf("Note that this does not delete anything\n");
    num=length(tifflistbox.Value);
    newitems={};
    n=0;
    for k=1:length(tifflistbox.Items)
        remove=false;
        for v=1:num
            if(strcmp(tifflistbox.Value{v},tifflistbox.Items{k}))
                remove=true;
            end
        end
        if(remove==false)   
            n=n+1;
            newitems{n}=tifflistbox.Items{k};
        end
    end
    tifflistbox.Items=convertStringsToChars(newitems);
    drawnow
end


function x=findthechars(x)
     z=0;
     if(isa(x,'numeric'))
         x=num2str(x);
     end
     while(~ischar(x) && z<100)
         z=z+1;
         disp("+");
         if(isstring(x))
             x=convertStringsToChars(x);
         elseif iscell(x)
             x=x{1};
         end
     end
end

function buttonAddevent(src,event)
    global allfileslist
    global includedfileslist
    global numcombos
    global combonames;
    num=size(allfileslist.Value,2);
        for v=1:num
            numcombos=numcombos+1;
            combonames{numcombos}=allfileslist.Value{v};
        end
        newitems={};
        n=0;
        for k=1:length(allfileslist.Items)
            remove=false;
            for v=1:num
                if(strcmp(allfileslist.Value{v},allfileslist.Items{k}))
                    remove=true;
                end
            end
            if(remove==false)   
                n=n+1;
                newitems{n}=allfileslist.Items{k};
            end
       end
        allfileslist.Items=newitems;
        includedfileslist.Items=combonames;
        drawnow
end

function buttonQuitevent(src, event)
    global quit
    global uif
    x=uiconfirm(uif,'Quit?','Quit','Icon','warning','Options',{'Yes, quit', 'No, stay'},'DefaultOption',2,'CancelOption',2);
    if(strcmp(x,'Yes, quit'))
        quit=1;
    end
end

function buttonLoadevent(src, event)
    global allfileslist
    global filelist;
    global dirlist;
    global filenum;
    global fulllist
    global imsnum
    global extension
    global filenum2
    global uif
    global filesep
    global depth
    global hasstitchedfiles;
    filelist={};
    dirlist={};
    fulllist={};
    filenum=0;
    filenum2=0;
    depth=0;
    imsnum=0;
    blockui("Scanning...");
    dir2=uigetdir("c:\");
    focus(uif);
    exploredir(dir2);
    imsnum=0;
    for t=1:filenum
        n=filelist{t};
        if(length(n)>5)
            if strcmp(     n(    (length(n)-3)  : length(n)     )    ,extension)
                imsnum=imsnum+1;
                dlt=dirlist{t};
                if(dlt(length(dlt))~=filesep)
                    dlt(length(dlt)+1)=filesep;
                end
                fulllist{imsnum}=strcat(dlt,filelist{t});
                x=strfind(fulllist{t},'FusionStitcher');
                if size(x,1)>0
                    hasstitchedfiles=true;
                end

            end
        end
    end
for t=1:imsnum
    allfileslist.Items{length(allfileslist.Items)+1}=fulllist{t};
end
    showui();
    drawnow
end

function buttonLoadTiffFolderevent(src, event)
    global allfileslist
    global filelist;
    global dirlist;
    global filenum;
    global fulllist
    global tifflistbox
    global imsnum
    global extension
    global filenum2
    global uif
    global depth
    global filesep;
    global hasstitchedfiles;
    filelist={};
    dirlist={};
    fulllist={};
    filenum=0;
    filenum2=0;
    depth=0;
    imsnum=0;
    blockui("Scanning...");
    dir2=uigetdir("c:\");
    focus(uif);
    exploredir2(dir2);
    imsnum=0;
    for t=1:filenum
        n=filelist{t};
        if(length(n)>5)
            if strcmp( upper(     n(    (length(n)-4)  : length(n)     ) )   ,'.TIFF')
                imsnum=imsnum+1;
                dlt=dirlist{t};
                if(dlt(length(dlt))~=filesep)
                    dlt(length(dlt)+1)=filesep;
                end
                fulllist{imsnum}=strcat(dlt,filelist{t});
            end
        end
    end
for t=1:imsnum
    tifflistbox.Items{length(tifflistbox.Items)+1}=fulllist{t};
end
    showui();
    drawnow
end

function buttonLoad2event(src, event)
    global vname
    global vnames
    global vloc
    global allfileslist
    global filelist;
    global dirlist;
    global filenum;
    global fulllist
    global imsnum
    global extension
    global filenum2
    global depth
    global filesep
    global hasstitchedfiles
    filelist={};
    dirlist={};
    fulllist={};
    filenum=0;
    filenum2=0;
    depth=0;
    imsnum=0;
    global uif
    blockui("Scanning...");
    [vnames,vloc]=uigetfile("*.ims","Pick one or more IMS files",vname,'MultiSelect','on');
    focus(uif);

    if(vloc~=0)
        if(vloc(length(vloc))~=filesep)
            vloc(length(vloc)+1)=filesep;
        end
        if(~iscellstr(vnames))
            n=vnames;
            if(length(n)>5)
                if strcmp(     n(    (length(n)-3)  : length(n)     )    ,extension)
                    imsnum=1;
                    filelist{1}=vnames;
                    fulllist{1}=strcat(vloc,vnames);
                    filenum=1;
                    x=strfind(fulllist{1},'FusionStitcher');
                    if size(x,1)>0
                        hasstitchedfiles=true;
                    end
                end
            end
        else
            filenum=size(vnames,2);
            for t=1:filenum
                n=vnames{t};
                if(length(n)>5)
                    if strcmp(     n(    (length(n)-3)  : length(n)     )    ,extension)
                        imsnum=imsnum+1;
                        filelist{imsnum}=n;
                        fulllist{imsnum}=strcat(vloc,n);
                        x=strfind(fulllist{imsnum},'FusionStitcher');
                        if size(x,1)>0
                            hasstitchedfiles=true;
                        end
                    end
                end
            end
        end
    end
    for t=1:imsnum
        allfileslist.Items{length(allfileslist.Items)+1}=fulllist{t};
    end
    showui();
    drawnow
end

function blockui(txt2)
    global giantbutton
    global grid
    global blockedalready;
    if blockedalready
        giantbutton.Text=sprintf('%s\n%s',giantbutton.Text,txt2);
    else
        giantbutton=uibutton(grid,'Text',txt2,'WordWrap','on','ButtonPushedFcn',@(src,event)buttonGiantbuttonevent(src,event));
        giantbutton.Tag='#f5f5f5';
        giantbutton.Layout.Row=[1 16];
        giantbutton.Layout.Column=[1 10];
        blockedalready=true;
    end
    drawnow
end

function buttonGiantbuttonevent(src,event)
global interrupt
global uif
global doanything
if doanything
    x=uiconfirm(uif,'Stop?','Stop?','Options',{'Yes', 'No'},'DefaultOption',2,'CancelOption',2);
    interrupt=false;
    if(strcmp(x,'Yes'))
        interrupt=true;
    end
end
end

function showui
    global giantbutton
    global blockedalready;
    global notelines

    notelines=1;
    blockedalready=false;
    try
        delete(giantbutton);
    catch SDFGSDF
        %was already shown
    end
    drawnow
end

function appendtonote(txt)
    global giantbutton
    global addnewline
global notelines
    
    try
        cc='';
        if addnewline
            cc=sprintf('\n');
            notelines=notelines+1;
        end
        if(notelines>30)
            notelines=1;
            giantbutton.Text=sprintf("%s",txt);
        else
            giantbutton.Text=sprintf('%s%s%s',giantbutton.Text,cc,txt);
        end
        catch asdfasd
        blockui(txt);
    end
    fprintf('%s\n',txt);
    addnewline=true;
    drawnow
end

function appendtonote2(txt)
    global giantbutton
    global addnewline
global notelines
    try
        cc='';
        if addnewline
            cc=sprintf('\n');
            notelines=notelines+1;
        end
        if notelines>30
            notelines=1;
            giantbutton.Text=sprintf("%s",txt);
        else
            giantbutton.Text=sprintf('%s%s%s',giantbutton.Text,cc,txt);
        end
    catch asdfasd
        blockui(txt);
    end
    addnewline=false;
    fprintf('%s\n',txt);
    drawnow
end


function textboxevent(src,event)
end

function exploredir(dir1)
    files=dir(dir1);
    global filelist
    global dirlist
    global filenum
    global filenum2
    global extension
    global imsnum
    global filesep;
    global maxfilechecks
    global maxfiles
    global depth

    fprintf("%i %i %i %s\n ",depth, filenum2, filenum, dir1);
    if (filenum2>maxfilechecks) || (filenum>maxfiles) || (depth>10)
        return
    end
    for t=3:size(files,1)
        filenum2=filenum2+1;
        if(~files(t).isdir)
            n=files(t).name;
            if(length(n)>5)
                if strcmp(     n(    (length(n)-3)  : length(n)     )    ,extension)
                    imsnum=imsnum+1;
                    filenum=filenum+1;
                    filelist{filenum}=files(t).name;
                    dirlist{filenum}=files(t).folder;
                    dlt=dirlist{filenum};
                    if(dlt(length(dlt))~=filesep)
                        dlt(length(dlt)+1)=filesep;
                    end
                    dirlist{filenum}=dlt;
                    fprintf("%s\n",strcat(dlt,files(t).name));
                    fulllist{filenum}=strcat(dlt,filelist{filenum});
                end
            end
        end
    end
    for t=3:size(files,1)
        if(files(t).isdir)
            filenum2=filenum2+1;
            if(~strcmp(files(t).name,'.') && ~strcmp(files(t).name,'..'))
                depth=depth+1;
                flt=files(t).folder;
                if(flt(length(flt))~=filesep)
                    flt(length(flt)+1)=filesep;
                end
                exploredir(strcat(flt,files(t).name));
                depth=depth-1;
            end
        end
    end
end

function exploredir2(dir1)
    files=dir(dir1);
    global filelist
    global dirlist
    global filenum
    global filenum2
    global extension
    global filesep
    global imsnum
    global maxfilechecks
    global maxfiles
    global depth

    fprintf("%i %i %i %s\n ",depth, filenum2, filenum, dir1);
    if (filenum2>maxfilechecks) || (filenum>maxfiles) || (depth>10)
        return
    end
    for t=3:size(files,1)
        filenum2=filenum2+1;
        if(~files(t).isdir)
            n=files(t).name;
            if(length(n)>5)
                if strcmp(   upper(  n(    (length(n)-4)  : length(n)     ))    ,'.TIFF')
                    imsnum=imsnum+1;
                    filenum=filenum+1;
                    filelist{filenum}=files(t).name;
                    dirlist{filenum}=files(t).folder;
                    dlt=dirlist{filenum};
                    if(dlt(length(dlt))~=filesep)
                        dlt(length(dlt)+1)=filesep;
                    end
                    dirlist{filenum}=dlt;
                    fprintf("%s\n",strcat(dlt,files(t).name));
                    fulllist{filenum}=strcat(dlt,filelist{filenum});
                end
            end
        end
    end
    for t=3:size(files,1)
        if(files(t).isdir)
            filenum2=filenum2+1;
            if(~strcmp(files(t).name,'.') && ~strcmp(files(t).name,'..'))
                depth=depth+1;
                flt=files(t).folder;
                if(flt(length(flt))~=filesep)
                    flt(length(flt)+1)=filesep;
                end
                exploredir2(strcat(flt,files(t).name));
                depth=depth-1;
            end
        end
    end
end


function processFile(pathname)
    global currentprocess
    global numtoprocess
    global interrupt
    global doanything
    global comboch1
    global comboch2
    global comboch3
    global fourchannel
    global skipexistingbutton

    doanything=true;
    if(interrupt)
        showui();
        return
    end
    blockui("Click this area to cancel (eventually)");
    sss=dir(pathname);
    appendtonote(sprintf("Processing file %i of %i (%i MB)",currentprocess,numtoprocess,floor(sss.bytes/1024/1024)));
    appendtonote(pathname);
    flen=length(pathname);
    if flen<5
        appendtonote("File name not correct.")
        return
    end
    if ~strcmp(".ims",pathname((flen-3):flen))
        appendtonote("Not an .IMS file.")
        return
    end
    tic

    try
        z=h5readatt(pathname,'/DataSetInfo/Channel 0/','Name');
        ch0txt='';
        for t=1:size(z,1)
            ch0txt(t)=z{t};
        end
        newname=strcat(pathname(1:(flen-4)),"-maxCh0-",ch0txt(1:3),".TIFF");
        if ~skipexistingbutton.Value || ~isfile(newname)
            appendtonote("Reading channel 0... ")
            chans=0;
            irdata=h5read(pathname,'/DataSet/ResolutionLevel 0/TimePoint 0/Channel 0/Data');
            appendtonote("Max-projecting... ")
            irmax0=irdata(:,:,1);
            for irt2=2:size(irdata,3)
                irmax0=max(irmax0,irdata(:,:,irt2));
            end
            appendtonote("Saving image... ")
            %a=prctile(irmax0(:),90);
            %b=prctile(irmax0(:),99);
            %c=prctile(irmax0(:),99.9);
            %    newname=strcat(pathname(1:(flen-4)),"-maxCh0 (90%=",num2str(a),", 99%=", num2str(b),", 99.9%=",num2str(c),").TIFF");
            addtotifflist(newname);
            imwrite(irmax0,newname);
            chans=1;
    
            appendtonote("Reading channel 1... ")
            if(interrupt)
                showui();
                return
            end
    
            try
                z=h5readatt(pathname,'/DataSetInfo/Channel 1/','Name');
                ch1txt='';
                for t=1:size(z,1)
                    ch1txt(t)=z{t};
                end
                irdata=h5read(pathname,'/DataSet/ResolutionLevel 0/TimePoint 0/Channel 1/Data');
                irmax1=irdata(:,:,1);
                appendtonote("Max-projecting... ")
                for irt2=2:size(irdata,3)
                    irmax1=max(irmax1,irdata(:,:,irt2));
                end
                appendtonote("Saving image... ")
                %   a=prctile(irmax1(:),90);
                %   b=prctile(irmax1(:),99);
                %   c=prctile(irmax1(:),99.9);
                %        newname=strcat(pathname(1:(flen-4)),"-maxCh1 (90%=",num2str(a),", 99%=", num2str(b),", 99.9%=",num2str(c),").TIFF");
                newname=strcat(pathname(1:(flen-4)),"-maxCh1-",ch1txt(1:3),".TIFF");
                addtotifflist(newname);
                imwrite(irmax1,newname);
                chans=2;
            catch ERRR
                appendtonote("No channel 1.");
            end
    
            appendtonote("Reading channel 2... ")
            if(interrupt)
                showui();
                return
            end
            try
                z=h5readatt(pathname,'/DataSetInfo/Channel 2/','Name');
                ch2txt='';
                for t=1:size(z,1)
                    ch2txt(t)=z{t};
                end
                irdata=h5read(pathname,'/DataSet/ResolutionLevel 0/TimePoint 0/Channel 2/Data');
                irmax2=irdata(:,:,1);
                appendtonote("Max-projecting... ")
                for irt2=2:size(irdata,3)
                    irmax2=max(irmax2,irdata(:,:,irt2));
                end
                appendtonote("Saving image... ")
                %  a=prctile(irmax2(:),90);
                %  b=prctile(irmax2(:),99);
                %  c=prctile(irmax2(:),99.9);
                newname=strcat(pathname(1:(flen-4)),"-maxCh2-",ch2txt(1:3),".TIFF");
    
                %        newname=strcat(pathname(1:(flen-4)),"-maxCh2 (90%=",num2str(a),", 99%=", num2str(b),", 99.9%=",num2str(c),").TIFF");
                addtotifflist(newname);
    
                imwrite(irmax2,newname);
                chans=3;
            catch ERRR
                appendtonote("No channel 2.");
            end
    
            appendtonote("Reading channel 3... ")
            if(interrupt)
                showui();
                return
            end
    
            try
                z=h5readatt(pathname,'/DataSetInfo/Channel 3/','Name');
                ch3txt='';
                for t=1:size(z,1)
                    ch3txt(t)=z{t};
                end
                irdata=h5read(pathname,'/DataSet/ResolutionLevel 0/TimePoint 0/Channel 3/Data');
                irmax3=irdata(:,:,1);
                appendtonote("Max-projecting... ")
                for irt2=2:size(irdata,3)
                    irmax3=max(irmax3,irdata(:,:,irt2));
                end
                appendtonote("Saving image... ")
                %   a=prctile(irmax3(:),90);
                %   b=prctile(irmax3(:),99);
                %   c=prctile(irmax3(:),99.9);
                newname=strcat(pathname(1:(flen-4)),"-maxCh3-",ch3txt(1:3),".TIFF");
                addtotifflist(newname);
                imwrite(irmax3,newname);
                chans=4;
            catch ERRR
                appendtonote("No channel 3.");
            end
            if(interrupt)
                showui();
                return
            end
    
            appendtonote(sprintf("Max projection %i x %i x %i slices... ",size(irdata,2),size(irdata,1),size(irdata,3)));
            if chans==3
                %        newname=strcat(pathname(1:(flen-4)),"-maxCh0,1,2.TIFF");
                newname=strcat(pathname(1:(flen-4)),"-maxCh0(",ch0txt(1:3),"),1(",ch1txt(1:3),"),2(",ch2txt(1:3),").TIFF");
                addtotifflist(newname);
                imwrite(cat(3,irmax0,irmax1,irmax2),newname);
            end
            if chans==2
                newname=strcat(pathname(1:(flen-4)),"-maxCh0(",ch0txt(1:3),"),1(",ch1txt(1:3),"),0(",ch0txt(1:3),").TIFF");
                addtotifflist(newname);
                imwrite(cat(3,irmax0,irmax1,irmax0),newname);
            end
            if chans==4
                switch comboch1
                    case 0
                        fch1=irmax0;
                        ch1str=strcat('0(',ch0txt(1:3));
                    case 1
                        fch1=irmax1;
                        ch1str=strcat('1(',ch1txt(1:3));
                    case 2
                        fch1=irmax2;
                        ch1str=strcat('2(',ch2txt(1:3));
                    case 3
                        fch1=irmax3;
                        ch1str=strcat('3(',ch3txt(1:3));
                end
                switch comboch2
                    case 0
                        fch2=irmax0;
                        ch2str=strcat('),0(',ch0txt(1:3));
                    case 1
                        fch2=irmax1;
                        ch2str=strcat('),1(',ch1txt(1:3));
                    case 2
                        fch2=irmax2;
                        ch2str=strcat('),2(',ch2txt(1:3));
                    case 3
                        fch2=irmax3;
                        ch2str=strcat('),3(',ch3txt(1:3));
                end
                switch comboch3
                    case 0
                        fch3=irmax0;
                        ch3str=strcat('),0(',ch0txt(1:3),').TIFF');
                    case 1
                        fch3=irmax1;
                        ch3str=strcat('),1(',ch1txt(1:3),').TIFF');
                    case 2
                        fch3=irmax2;
                        ch3str=strcat('),2(',ch2txt(1:3),').TIFF');
                    case 3
                        fch3=irmax3;
                        ch3str=strcat('),3(',ch3txt(1:3),').TIFF');
                end
                %        newname=strcat(pathname(1:(flen-4)),"-maxCh0,2,3.TIFF");
                %        newname=strcat(pathname(1:(flen-4)),"-maxCh",num2str(comboch1),",",num2str(comboch2),",",num2str(comboch3),".TIFF");
                newname=strcat(pathname(1:(flen-4)),"-maxCh",ch1str,ch2str,ch3str);
                addtotifflist(newname);
                imwrite(cat(3, fch1, fch2, fch3),newname);
                newname=strcat(pathname(1:(flen-4)),"-maxCh0(",ch0txt(1:3),"),1(",ch1txt(1:3),"),2(",ch2txt(1:3),"),3(",ch3txt(1:3),")-multipage.TIFF");
                imwrite(irmax0,newname);
                imwrite(irmax1,newname,"WriteMode","append");
                imwrite(irmax2,newname,"WriteMode","append");
                imwrite(irmax3,newname,"WriteMode","append");
                fourchannel=true;
            end
            b=toc;
            appendtonote(sprintf("Took %f seconds (%i MB/sec).",b,floor(sss.bytes/b/1024/1024)));
        else
            appendtonote("Ch0 max TIFF exists -- skipping file.");
        end
    catch NOFILEERR
        appendtonote("Can't open file. Skipping.");
    end
    showui();
    doanything=false;
end

function addtotifflist(name)
global tifflistbox
global tifflistfiles
tifflistbox.Items{length(tifflistbox.Items)+1}=convertStringsToChars(name);
tifflistfiles{length(tifflistfiles)+1}=convertStringsToChars(name);
drawnow
end


function buttonBrightenAllevent(src,event)
    global tifflistbox
    num=length(tifflistbox.Value);
    tp=99.8;
    tp=str2double(inputdlg("Input percentile to set as max","Input percentile",[1 45],{num2str(tp)}));
    if isnan(tp)
        tp=99.8;
    end
    maxes=zeros(num,4,'double');
    maxmaxes=zeros(4,1,'double');
    chans=zeros(num);
    images={};
    code=char(floor(26*rand(1, 4)) + 65);
    blockui(sprintf("Normalizing brightnesses in group %s",code));
    for t=1:num
        appendtonote(sprintf("Reading image %i",t));
        images{t}=im2double(imread(tifflistbox.Value{t}));
        i=images{t};
        chans(t)=size(i,3);
        for j=1:chans(t)
            i3=i(:,:,j);
            i2=reshape(i3(:,:),[],1);
            mm=prctile(i2,tp);
            maxes(t,j)=mm;
            maxmaxes(j)=max(maxmaxes(j),mm);
        end
    end
    appendtonote(sprintf("Max brightnesses: %f, %f, %f, %f",maxmaxes(1),maxmaxes(2), maxmaxes(3), maxmaxes(4)));
    for t=1:num
        appendtonote(sprintf("Writing image %i",t));
        i=images{t};
        i2=cell(3,1);
        for c=1:chans(t)
                i2{c}=i(:,:,c);
                i2{c}=i2{c}/maxmaxes(c);
        end
        switch c
            case 1
                i=i2{1};
            case 2
                i=cat(2,i2{1},i2{2});
            case 3
                i=cat(3,i2{1},i2{2},i2{3});
        end
        name=tifflistbox.Value{t};
        name=strcat(name,' (brightened ',code,').TIFF');
        imwrite(im2uint16(i),name);
        addtotifflist(name);
    end
    showui();
end

function buttonBrightenAll2event(src,event)
    global tifflistbox
    num=length(tifflistbox.Value);
    tp=99.8;
    tp=str2double(inputdlg("Input percentile to set as max","Input percentile",[1 45],{num2str(tp)}));
    m=10;
    m=str2double(inputdlg("Input percentile to set as zero","Input percentile",[1 45],{num2str(m)}));
    if isnan(tp)
        tp=99.8;
    end
    if isnan(m)
        m=10;
    end

    blockui(sprintf("Normalizing brightnesses individually to %f percentile=MAX",tp));
    for t=1:num
        name=tifflistbox.Value{t};
        appendtonote2(sprintf("Image %s: Reading...",name));
        i=im2double(imread(tifflistbox.Value{t}));
        ch=size(i,3);
        appendtonote2(" Math...");
        if ch==1
%            zmax=max(max(i(:,:)));
            i2=reshape(i(:,:),[],1);
            mm=prctile(i2,m);
            i=i-m;
            i2=i2-m;
            zmax=prctile(i2,tp);
            i=i/zmax;
        end
        if(ch==3)
            z1=i(:,:,1);
            i2=reshape(z1(:,:),[],1);
            mm=prctile(i2,m);
            z1=z1-mm;
            i2=i2-mm;
            z1max=prctile(i2,tp);
            z1=z1/z1max;
 
            z2=i(:,:,2);
            i2=reshape(z2(:,:),[],1);
            mm=prctile(i2,m);
            z2=z2-mm;
            i2=i2-mm;
            z2max=prctile(i2,tp);
            z2=z2/z2max;

            z3=i(:,:,3);
            i2=reshape(z3(:,:),[],1);
            mm=prctile(i2,m);
            z3=z3-mm;
            i2=i2-mm;
            z3max=prctile(i2,tp);
            z3=z3/z3max;

            i=cat(3,z1,z2,z3);
        end
        if(ch==2)
            z1=i(:,:,1);
            i2=reshape(z1(:,:),[],1);
            mm=prctile(i2,m);
            z1=z1-m;
            i2=i2-m;
            z1max=prctile(i2,tp);
            z1=z1/z1max;

            z2=i(:,:,2);
            i2=reshape(z2(:,:),[],1);
            mm=prctile(i2,m);
            z2=z2-m;
            i2=i2-m;
            z2max=prctile(i2,tp);
            z2=z2/z2max;

            i=cat(2,z1,z2);
        end
        if(ch<4)
            appendtonote(" Saving.");
            name2=strcat(name(1:length(name)-5),' (brightened ',num2str(m),'-',num2str(tp),').TIFF');
            imwrite(im2uint16(i),name2);
            addtotifflist(name2);
        end
    end
    showui();
end


function buttonShrinkevent(src,event)
    global tifflistbox
    sf=20;
    sf=str2double(inputdlg("Input final size as percentage of original (each dim)","Input size",[1 45],{num2str(sf)}));
    if isnan(sf)
        sf=20;
    end
    sf=sf/100;
    num=length(tifflistbox.Value);
    blockui("Shrinking images");
    for t=1:num
        name=tifflistbox.Value{t};
        name2=strcat(name(1:length(name)-5),' small',num2str(sf*100),'.TIFF');
        appendtonote2(sprintf("%s: Reading...",name));
        i=imread(tifflistbox.Value{t});
        appendtonote2(" Resizing...");
        i2=imresize(i,sf);
        appendtonote(" Saving.");
        imwrite(i2,name2);
        addtotifflist(name2);
    end
    showui();
end

function buttonFlipLRevent(src,event)
    global tifflistbox
    num=length(tifflistbox.Value);
    blockui("Flipping images LR");
    for t=1:num
        name=tifflistbox.Value{t};
        name2=strcat(name(1:length(name)-5),' flipLR.TIFF');
        appendtonote(sprintf("Doing image %s",name));
        imwrite(flipdim(imread(tifflistbox.Value{t}),2),name2);
        addtotifflist(name2);
    end
    showui();
end

function fourchannelsevent(src, event)
global comboch1
global comboch2
global comboch3
global finished1

finished1=false;
blockui("Select channels");
uif1=uifigure;
grid1=uigridlayout(uif1);
grid1.BackgroundColor='#999999';
%                1   2   3   4   5   6  7   8   9   10   11
grid1.RowHeight={16, 32, 32, 32, 32, 40, 40, 40, 40, 40};
uif1.Position(4)=200;
%diff=400;
label1=uilabel(grid1);
label1.Layout.Row=1;
label1.Layout.Column=1;
label1.Text="Output Ch1:";
label2=uilabel(grid1);
label2.Layout.Row=1;
label2.Layout.Column=2;
label2.Text="Output Ch2:";
label3=uilabel(grid1);
label3.Layout.Row=1;
label3.Layout.Column=3;
label3.Text="Output Ch3:";

ch1grp=uibuttongroup(grid1);
ch1grp.Layout.Column=1;
ch1grp.Layout.Row=[2 4];
ch1ch0=uiradiobutton(ch1grp,"Text","Channel 0");
ch1ch1=uiradiobutton(ch1grp,"Text","Channel 1");
ch1ch2=uiradiobutton(ch1grp,"Text","Channel 2");
ch1ch3=uiradiobutton(ch1grp,"Text","Channel 3");
ch1ch3.Position=[6 10 100 24];
ch1ch2.Position=[6 34 100 24];
ch1ch1.Position=[6 58 100 24];
ch1ch0.Position=[6 82 100 24];

ch2grp=uibuttongroup(grid1);
ch2ch0=uiradiobutton(ch2grp,"Text","Channel 0");
ch2ch1=uiradiobutton(ch2grp,"Text","Channel 1");
ch2ch2=uiradiobutton(ch2grp,"Text","Channel 2");
ch2ch3=uiradiobutton(ch2grp,"Text","Channel 3");
ch2grp.Layout.Column=2;
ch2grp.Layout.Row=[2 4];
ch2ch3.Position=[6 10 100 26];
ch2ch2.Position=[6 34 100 26];
ch2ch1.Position=[6 58 100 26];
ch2ch0.Position=[6 82 100 26];


ch3grp=uibuttongroup(grid1);
ch3ch0=uiradiobutton(ch3grp,"Text","Channel 0");
ch3ch1=uiradiobutton(ch3grp,"Text","Channel 1");
ch3ch2=uiradiobutton(ch3grp,"Text","Channel 2");
ch3ch3=uiradiobutton(ch3grp,"Text","Channel 3");
ch3grp.Layout.Column=3;
ch3grp.Layout.Row=[2 4];
ch3ch3.Position=[6 10 100 26];
ch3ch2.Position=[6 34 100 26];
ch3ch1.Position=[6 58 100 26];
ch3ch0.Position=[6 82 100 26];

buttonQuit1=uibutton(grid1,'Text',"OK",'ButtonPushedFcn',@(src,event)buttonQuit1event(src,event));
buttonQuit1.Tag='#f5f5f5';
buttonQuit1.Layout.Row=5;
buttonQuit1.Layout.Column=3;

switch comboch1
    case 0
        ch1ch0.Value=true;
    case 1
        ch1ch1.Value=true;
    case 2
        ch1ch2.Value=true;
    case 3
        ch1ch3.Value=true;
end
switch comboch2
    case 0
        ch2ch0.Value=true;
    case 1
        ch2ch1.Value=true;
    case 2
        ch2ch2.Value=true;
    case 3
        ch2ch3.Value=true;
end
switch comboch3
    case 0
        ch3ch0.Value=true;
    case 1
        ch3ch1.Value=true;
    case 2
        ch3ch2.Value=true;
    case 3
        ch3ch3.Value=true;
end

while finished1==false
    pause(0.1);
end
if ch1ch0.Value
    comboch1=0;
end
if ch1ch1.Value
    comboch1=1;
end
if ch1ch2.Value
    comboch1=2;
end
if ch1ch3.Value
    comboch1=3;
end

if ch2ch0.Value
    comboch2=0;
end
if ch2ch1.Value
    comboch2=1;
end
if ch2ch2.Value
    comboch2=2;
end
if ch2ch3.Value
    comboch2=3;
end

if ch3ch0.Value
    comboch3=0;
end
if ch3ch1.Value
    comboch3=1;
end
if ch3ch2.Value
    comboch3=2;
end
if ch3ch3.Value
    comboch3=3;
end
close(uif1);
showui();
end

function buttonQuit1event(src,event)
global finished1
finished1=true;
end

function buttonPropertiesevent(src,event)
global finished1
global allfileslist
global includedfileslist
global uif
msg='';
try
    z=h5readatt(includedfileslist.Value,'/DataSetInfo/Channel 0/','Name');
    txt='';
    for t=1:size(z,1)
    txt(t)=z{t};
    end
    msg=sprintf('%sChannel 0: %s\n',msg,txt);
catch DFGSDF
end

try
    z=h5readatt(includedfileslist.Value,'/DataSetInfo/Channel 1/','Name');
    txt='';
    for t=1:size(z,1)
    txt(t)=z{t};
    end
    msg=sprintf('%sChannel 1: %s\n',msg,txt);
catch DFGSDF
end

try
    z=h5readatt(includedfileslist.Value,'/DataSetInfo/Channel 2/','Name');
    txt='';
    for t=1:size(z,1)
    txt(t)=z{t};
    end
    msg=sprintf('%sChannel 2: %s\n',msg,txt);
catch DFGSDF
end

try
    z=h5readatt(includedfileslist.Value,'/DataSetInfo/Channel 3/','Name');
    txt='';
    for t=1:size(z,1)
    txt(t)=z{t};
    end
    msg=sprintf('%sChannel 3: %s\n',msg,txt);
catch DFGSDF
end
    uiconfirm(uif,msg,"Notice");

end

function buttonAddTiffFilesevent(src, event)
    global vname
    global vnames
    global vloc
    global allfileslist
    global filelist;
    global tifflistbox
    global dirlist;
    global filenum;
    global fulllist
    global imsnum
    global extension
    global filenum2
    global depth
    global filesep
    global hasstitchedfiles
    filelist={};
    dirlist={};
    fulllist={};
    filenum=0;
    filenum2=0;
    depth=0;
    imsnum=0;
    global uif
    blockui("Scanning...");
    [vnames,vloc]=uigetfile("*.tiff","Pick one or more TIFF files",vname,'MultiSelect','on');
    focus(uif);

    if(vloc~=0)
        if(vloc(length(vloc))~=filesep)
            vloc(length(vloc)+1)=filesep;
        end
        if(~iscellstr(vnames))
            n=vnames;
            if(length(n)>6)
                if strcmp(     upper( n(    (length(n)-4)  : length(n)     )   ) ,'.TIFF')
                    imsnum=1;
                    filelist{1}=vnames;
                    fulllist{1}=strcat(vloc,vnames);
                    filenum=1;
                end
            end
        else
            filenum=size(vnames,2);
            for t=1:filenum
                n=vnames{t};
                if(length(n)>5)
                    if strcmp(    upper( n(    (length(n)-4)  : length(n)     )  )  ,'.TIFF')
                        imsnum=imsnum+1;
                        filelist{imsnum}=n;
                        fulllist{imsnum}=strcat(vloc,n);
                    end
                end
            end
        end
    end
    for t=1:imsnum
        tifflistbox.Items{length(tifflistbox.Items)+1}=fulllist{t};
    end
    showui();
    drawnow
end

function buttonRechannelevent(src,event)
    global tifflistbox
    num=length(tifflistbox.Value);
    blockui("Rechanneling");
    rc='210';
    grc=inputdlg("Input new order of existing channels eg '210' or 'GBR'","Input channels",[1 45],{rc});
    rc=grc{1};
    if length(rc)~=3
        rc='123';
    end
    for t=1:length(rc)
        if(upper(rc(t))=='R')
            rc(t)='1';
        end
        if(upper(rc(t))=='G')
            rc(t)='2';
        end
        if(upper(rc(t))=='B')
            rc(t)='3';
        end
        if(rc(t)~='1' && rc(t)~='2' && rc(t)~='3')
            rc(t)='0';
        end
    end
    c1=min(3,max(0,str2num(rc(1))))
    c2=min(3,max(0,str2num(rc(2))))
    c3=min(3,max(0,str2num(rc(3))))
    for t=1:num
        name=tifflistbox.Value{t};
        name2=strcat(name(1:length(name)-5),' rechan123-',rc,'.TIFF');
        appendtonote(sprintf("Doing image %s",name));
        i=imread(tifflistbox.Value{t});
        chan=size(i,3);
        blank=zeros(size(i,1),size(i,2));
        chans=cell(4,1);
        chans{1}=blank;
        chans{2}=blank;
        chans{3}=blank;
        chans{4}=blank;
        for c=1:chan
            chans{c+1}=i(:,:,c);
        end
        i2=cat(3,chans{c1+1},chans{c2+1},chans{c3+1});
        imwrite(i2,name2);
        addtotifflist(name2);
    end
    showui();

end

function buttonRenameevent(src,event)
    global tifflistbox
    global filesep
    num=length(tifflistbox.Value);
    blockui("Renaming");
    txt=sprintf("New filenames start with...");
    rc='Shortname';
    prefix=inputdlg(txt,"Text",[1 45],{rc});

    rc='FusionStitcher_F';
    txt=sprintf("In all filenames, following this example:\n%s\nFind the sequence...",tifflistbox.Value{1});
    grc=inputdlg(txt,"Text",[1 45],{rc});

    rc='2';
    z=inputdlg("And take this many characters for the new name","How many",[1 45],{rc});
    z=z{1};
    chs=str2num(z);
    if isnan(chs)
        chs=2;
    end
    grc=grc{1};
    gl=length(grc);
    for t=1:num
        name=tifflistbox.Value{t};
        x=strfind(name,grc);
        z='';
        if size(x,2)>0
            x=x(1);
            slash=false;
            for l=length(name):-1:1
                if(name(l)==filesep)
                    slash=true;
                    break
                end
            end
            if(slash)
                for n=x+gl:x+gl+chs-1
                    z(length(z)+1)=name(n);
                end
                name2=strcat(name(1:l),prefix,z,'.TIFF');
    if isfile(name2{1})
        fprintf("BAD RENAME: file exists %s\n",findthechars(name2));
    end
    fprintf("%s --> %s\n",findthechars(name),findthechars(name2{1}));
    copyfile(name,findthechars(name2{1}));
    addtotifflist(findthechars(name2{1}));
            end
        end
    end
    showui();
end


function buttonPNGevent(src,event)
    global tifflistbox
    global filesep
    num=length(tifflistbox.Value);
    blockui("Exporting as PNG");
    for t=1:num
        name=tifflistbox.Value{t};
        appendtonote2(sprintf("Image %s: loading...",name));
        i=imread(name);
        appendtonote2(" and saving...");
try
    imwrite(i,strcat(name(1:length(name)-4),'PNG'));
        addtotifflist(strcat(name(1:length(name)-4),'PNG'));
catch
    fprintf("Can't save %s -- kill Windows Photo, etc\n");
end
    appendtonote(" done!");
    end
    showui();
end


function buttonSelectTIFFsevent(src,event)
    global tifflistbox
    global filesep
    num=length(tifflistbox.Value);
    blockui("Selecting");
    txt=sprintf("Select TIFF files containing...");
    rc='small';
    txt=inputdlg(txt,"Text",[1 45],{rc});
    txt=txt{1};
    for t=1:length(tifflistbox.Items)
        name=tifflistbox.Items{t};
        x=strfind(name,txt);
        if size(x,2)>0
            tifflistbox.Value{length(tifflistbox.Value)+1}=name;
        end
    end
    showui();
end


function buttonInvertTIFFsevent(src,event)
    global tifflistbox
    global filesep
    num=length(tifflistbox.Value);
    blockui("Inverting");
    oldselect=tifflistbox.Value;
    newselect={};
    for t=1:length(tifflistbox.Items)
        name=tifflistbox.Items{t};
        found=0;
        for k=1:length(oldselect)
            if(strcmp(name,oldselect{k}))
                %is selected
                found=1;
            end
        end
        if found==0
            newselect{length(newselect)+1}=name;
        end
    end
    tifflistbox.Value=newselect;
    showui();
end


function buttonDeleteTIFFevent(src,event)
    global tifflistbox
    blockui("Deleting");
    num=size(tifflistbox.Value,2);
    newitems={};
    n=0;
    for k=1:length(tifflistbox.Items)
        remove=false;
        for v=1:num
            if(strcmp(tifflistbox.Value{v},tifflistbox.Items{k}))
                remove=true;
                movefile(tifflistbox.Items{k},strcat(tifflistbox.Items{k},"DELETED"));
            end
        end
        if(remove==false)   
            n=n+1;
            newitems{n}=tifflistbox.Items{k};
        end
    end
    tifflistbox.Items=convertStringsToChars(newitems);
    showui();
    drawnow
end


function buttonConsolidateevent(src,event)
    global tifflistbox
    global uif
    global filesep
    blockui("Copy into folder...");
    dir2=uigetdir("c:\");
    focus(uif);
    if(dir2(length(dir2))~=filesep)
        dir2(length(dir2)+1)=filesep;
    end
    num=size(tifflistbox.Value,2);
    tlbv=tifflistbox.Value;
    newitems={};
    n=0;
    for k=1:length(tifflistbox.Items)
        remove=false;
        for v=1:num
            if(strcmp(tlbv{v},tifflistbox.Items{k}))
                remove=true;
                name=tifflistbox.Items{k};
                slash=false;
                for l=length(name):-1:1
                    if(name(l)==filesep)
                        slash=true;
                        break
                    end 
                end
                if(slash)
                    name2=strcat(dir2,name((l+1):(length(name))));
                    copyfile(name,name2);
                    tifflistbox.Items{k}=name2;
                end
            end
        end
    end
    showui();
    drawnow
end

function buttonoff(button)
    global origbuttoncolor;
    button.Value=false;
    if(button.Text(length(button.Text)) == char(10003))
       button.Text=button.Text(1:length(button.Text)-2); 
    end
    button.BackgroundColor=origbuttoncolor;
    button.FontColor="#000000";
end

function buttonon(button)
    global kured;
    button.Value=true;
    button.FontColor="#ffffff";
    if(button.Text(length(button.Text)) ~= char(10003))
        button.Text=strcat(button.Text," ", char(10003));
    end
    button.BackgroundColor=kured;
end
```
