## Supplemental code 2 for "Gut hormone signaling drives sex differences in metabolism and behavior": gutquant.m

```
global parentdir
parentdir="D:\Rewitz lab\max";
global swapchannels
swapchannels=true;
global validsuffix
validsuffix='-maxCh0,1,0.TIFF';

askaboutexisting=false;

    global alreadyblocked;
    alreadyblocked=false;
%global GFPchan
%GFPchan=2;
%global DAPIchan
%DAPIchan=1;

global fullfilename;
global justfilename;
global u;
global grid;
global cansave;
cansave=false;
global ch1s;
global andimg;
global ch2s;
global max1;
max1=1;
global min1;
global roibox;
global roiwidth;
global roiheight;
global roix;
global origbuttoncolor;
global kured;
global roiy;
global finishedbox;
finishedbox=false;
roibox=0;
roiwidth=500;
roix=100;
roiheight=500;
roiy=100;
global giantbutton;
global ch1p;
global ch2p;
ch1p=98;
ch2p=98;
global img;
img=0;
global max2;
max2=0;
global fullfilename;
global min2;
global ox1;
global ox2;
global ox3;
global ox4;
global ox5;
global ox6;
global minc1;
global minc2;
global imgname;
global origch1;
global origch2;
global origch3;
global ch1;
global butcount;
global zoomed;
global shownimg;
zoomed=6;
global ch2;
global ch3;
global c1;
global c2;
global image;
global counts;
global contin;
global zimg;
global newimg;
global grid;
global butch1;
global butch2;
global butshowand;
global butshowcounted;
global butreset;
global x1;
global x2;
global x3;
global x4;
global x5;
global x6;
global txt;
global w3;
global min1
global max1;
global max2;
global min2;
global maxch1;
global maxch2;
global line1;
global line2;
global line3;
global line4;
global line5;
global line6;
global butreverse;
global but9;
global w4;
global quit;
global reversed;
global butfull;
global butr1;
global butr2;
global butr3;
global butr4;
global butr5;

global stats1;
global stats2;
global stats3;
global stats4;
global stats5;
global index1;
global index2;
global index3;
global index4;
global index5;
global allstats;
global allindices;
allstats=cell(5,3);
allindices=cell(5,3);
stats1=cell(1,3);
stats2=cell(1,3);
stats3=cell(1,3);
stats4=cell(1,3);
stats5=cell(1,3);
index1=zeros(1,3);
index2=zeros(1,3);
index3=zeros(1,3);
index4=zeros(1,3);
index5=zeros(1,3);

global countedall;
global countedr;
countedr=zeros(6,1);
global countedimg;
reversed=false;
global newname;
warning off
quit=false;
txt="Nothing yet";
fs=dir(parentdir);
fn=size(fs,1);
fprintf("Number of things in the directory: %f\n",fn-2);
% 
% for t=3:fn
%     fullfilename=strcat(fs(t).folder,filesep,fs(t).name);
%     suffix=fullfilename( (size(fullfilename,2)-3)  :   size(fullfilename,2)    );
%     if(suffix=='.tif')
%         movefile(fullfilename, strcat(fullfilename,"f"));
%     end
% end


u=uifigure;
u.Position=[1000 100 2000 500];
grid=uigridlayout(u);
kugray="#666666";
kured="#901A1E";
grid.BackgroundColor=kugray;
grid.RowHeight={60,60,60,60,60,60};
grid.ColumnWidth={150,150,150,'2x',30,30,30,30,30};

butshowand=uibutton(grid,'state','Text','Show AND',"ValueChangedFcn",@(src,event)updatebutshowand(src,event));
butshowand.Layout.Row=2;
butshowand.Layout.Column=2;
origbuttoncolor=butshowand.BackgroundColor;
buttonoff(butshowand);


butc1decdec=uibutton(grid,'Text','--',"ButtonPushedFcn",@(src,event)updatebutc1decdec(src,event));
butc1decdec.Layout.Row=2;
butc1decdec.Layout.Column=5;
butc1dec=uibutton(grid,'Text','-',"ButtonPushedFcn",@(src,event)updatebutc1dec(src,event));
butc1dec.Layout.Row=2;
butc1dec.Layout.Column=6;
butc1reset=uibutton(grid,'Text','R',"ButtonPushedFcn",@(src,event)updatebutc1reset(src,event));
butc1reset.Layout.Row=2;
butc1reset.Layout.Column=7;
butc1inc=uibutton(grid,'Text','+',"ButtonPushedFcn",@(src,event)updatebutc1inc(src,event));
butc1inc.Layout.Row=2;
butc1inc.Layout.Column=8;
butc1incinc=uibutton(grid,'Text','++',"ButtonPushedFcn",@(src,event)updatebutc1incinc(src,event));
butc1incinc.Layout.Row=2;
butc1incinc.Layout.Column=9;

butc2decdec=uibutton(grid,'Text','--',"ButtonPushedFcn",@(src,event)updatebutc2decdec(src,event));
butc2decdec.Layout.Row=3;
butc2decdec.Layout.Column=5;
butc2dec=uibutton(grid,'Text','-',"ButtonPushedFcn",@(src,event)updatebutc2dec(src,event));
butc2dec.Layout.Row=3;
butc2dec.Layout.Column=6;
butc2reset=uibutton(grid,'Text','R',"ButtonPushedFcn",@(src,event)updatebutc2reset(src,event));
butc2reset.Layout.Row=3;
butc2reset.Layout.Column=7;
butc2inc=uibutton(grid,'Text','+',"ButtonPushedFcn",@(src,event)updatebutc2inc(src,event));
butc2inc.Layout.Row=3;
butc2inc.Layout.Column=8;
butc2incinc=uibutton(grid,'Text','++',"ButtonPushedFcn",@(src,event)updatebutc2incinc(src,event));
butc2incinc.Layout.Row=3;
butc2incinc.Layout.Column=9;


butch1=uibutton(grid,'state','Text','Show Ch1',"ValueChangedFcn",@(src,event)updateCh1b(src,event));
butch1.Layout.Row=2;
butch1.Layout.Column=3;
buttonon(butch1);

ch1s=uislider(grid, "ValueChangedFcn",@(src,event)updateCh1(src,event));
ch1s.Layout.Row=2;
ch1s.Layout.Column=4;

butch2=uibutton(grid,'state','Text','Show Ch2',"ValueChangedFcn",@(src,event)updateCh2b(src,event));
butch2.Layout.Row=3;
butch2.Layout.Column=3;
buttonon(butch2);

ch2s=uislider(grid, "ValueChangedFcn",@(src,event)updateCh2(src,event));
ch2s.Layout.Row=3;
ch2s.Layout.Column=4;

butreset=uibutton(grid,'Text','RESET',"ButtonPushedFcn",@(src,event)updatebutreset(src,event));
butreset.Layout.Row=1;
butreset.Layout.Column=3;

butcount=uibutton(grid,'Text','Count all and save',"ButtonPushedFcn",@(src,event)updatebutcount(src,event));
butcount.Layout.Row=4;
butcount.Layout.Column=3;

w3=uilabel(grid,"Text", txt);
w3.Layout.Row=4;
w3.Layout.Column=4;

butnext=uibutton(grid,'Text','NEXT!',"ButtonPushedFcn",@(src,event)updatebutnext(src,event));
butnext.Layout.Row=4;
butnext.Layout.Column=2;

butquit=uibutton(grid,'Text','Quit',"ButtonPushedFcn",@(src,event)updatebutquit(src,event));
butquit.Layout.Row=6;
butquit.Layout.Column=3;

w4=uilabel(grid,"Text", "LOADING...");
w4.Layout.Row=1;
w4.Layout.Column=4;

butreverse=uibutton(grid,'state','Text','Reverse',"ValueChangedFcn",@(src,event)updatebutreverse(src,event));
butreverse.Layout.Row=3;
butreverse.Layout.Column=2;

butfull=uibutton(grid,'state','Text','Show full',"ValueChangedFcn",@(src,event)updatebutfull(src,event));
butfull.Layout.Row=1;
butfull.Layout.Column=1;
buttonon(butch1);

butr1=uibutton(grid,'state','Text','Show zoom (R1)',"ValueChangedFcn",@(src,event)updatebutr1(src,event));
butr1.Layout.Row=2;
butr1.Layout.Column=1;
buttonoff(butr1);

butr2=uibutton(grid,'state','Text','Show zoom (R2)',"ValueChangedFcn",@(src,event)updatebutr2(src,event));
butr2.Layout.Row=3;
butr2.Layout.Column=1;
buttonoff(butr2);

butr3=uibutton(grid,'state','Text','Show zoom (R3)',"ValueChangedFcn",@(src,event)updatebutr3(src,event));
butr3.Layout.Row=4;
butr3.Layout.Column=1;
buttonoff(butr3);

butr4=uibutton(grid,'state','Text','Show zoom (R4)',"ValueChangedFcn",@(src,event)updatebutr4(src,event));
butr4.Layout.Row=5;
butr4.Layout.Column=1;
buttonoff(butr4);
butr5=uibutton(grid,'state','Text','Show zoom (R5)',"ValueChangedFcn",@(src,event)updatebutr5(src,event));
butr5.Layout.Row=6;
butr5.Layout.Column=1;
buttonoff(butr5);

butshowcounted=uibutton(grid,'state','Text','Show counted',"ValueChangedFcn",@(src,event)updatebutshowcounted(src,event));
butshowcounted.Layout.Row=1;
butshowcounted.Layout.Column=2;
buttonoff(butshowcounted);
butshowcounted.Visible='off';
global butgaussianblur;
butgaussianblur=uibutton(grid,'state','Text','Gaussian Blur',"ValueChangedFcn",@(src,event)updatebutgaussianblur(src,event));
butgaussianblur.Layout.Row=6;
butgaussianblur.Layout.Column=2;


global but10x;

but10x=uibutton(grid,'state','Text','Low brightness',"ValueChangedFcn",@(src,event)updatebut10x(src,event));
but10x.Layout.Row=5;
but10x.Layout.Column=3;
buttonoff(but10x);

butsetb=uibutton(grid,'Text','Set just above last click',"ButtonPushedFcn",@(src,event)updatebutsetb(src,event));
butsetb.Layout.Row=5;
butsetb.Layout.Column=2;

butbox=uibutton(grid,'Text','Erase...',"ButtonPushedFcn",@(src,event)updatebutbox(src,event));
butbox.Layout.Row=7;
butbox.Layout.Column=1;

butbox2=uibutton(grid,'Text','Crop...',"ButtonPushedFcn",@(src,event)updatebutbox2(src,event));
butbox2.Layout.Row=7;
butbox2.Layout.Column=2;
global butbox3;

butbox3=uibutton(grid,'Text','Save image data',"ButtonPushedFcn",@(src,event)updatebutbox3(src,event));
butbox3.Layout.Row=7;
butbox3.Layout.Column=3;
butbox3.Visible='off';

global comment;

comment=uitextarea(grid);
comment.Layout.Row=5;
comment.Layout.Column=4;
comment.Value="No comment yet";
drawnow

counts=zeros(5,3);
counts(:,:)=-1;
countedall=false;
countedr(1)=false;
countedr(2)=false;
countedr(3)=false;
countedr(4)=false;
countedr(5)=false;
countedr(6)=false;
buttonoff(butshowcounted);
butshowcounted.Visible='off';
butshowcounted.Text="Show counted";


for t=3:fn
    counts=zeros(5,3);
    counts(:,:)=-1;
    countedall=false;
    countedr(1)=false;
    countedr(2)=false;
    countedr(3)=false;
    countedr(4)=false;
    countedr(5)=false;
    countedr(6)=false;
    buttonon(butch2); 
    buttonon(butch1);
    buttonoff(butshowcounted); 
    butshowcounted.Visible='off';
    butshowcounted.Text="Show counted";
    buttonoff(butshowand);
    buttonon(butfull);
    buttonoff(butr1);
    buttonoff(butr2);
    buttonoff(butr3);
    buttonoff(butr4);
    buttonoff(butgaussianblur);
    buttonoff(butr5);
    butbox3.Visible='off';
    butreverse.Text="Reverse";
    reversed=0;
    comment.Value="No comment yet";
    counted=0;
    filesep='\';
    fullfilename=strcat(fs(t).folder,filesep,fs(t).name);
    justfilename=fs(t).name;
    skip=false;
    vslen=length(validsuffix);
    suffix=fullfilename( (size(fullfilename,2)-vslen+1)  :   size(fullfilename,2)    );
    if(~strcmp(suffix,validsuffix))
        skip=true;
    end
    preloaded=false;
    if skip==false
        minusfour=fullfilename(1:size(fullfilename,2)-4);
        newname=strcat(minusfour,'mat');
        imgname=fullfilename;
        if isfile(newname)
            fprintf("%s exists.\n",newname);
            skip=true;
            if(askaboutexisting)
                xx=questdlg(strcat(newname)," exists.",strcat(newname)," exists.","Load it and continue","Skip","Skip");
                if(xx=="Skip")
                    skip=true;
                else
                    skip=false;
                    load(newname);
                    preloaded=true;
                end
            end
        end
    end
    if skip==false
        fprintf("Loading %s.\n",imgname);
        blockui(sprintf("Please wait -- loading %s",imgname));
        img=double(imread(imgname));
        if(size(img,3)<3)
            skip=true;
        end
    end
    if skip==false
        if(swapchannels)
            origch1=img(:,:,2)/65535;
            origch2=img(:,:,1)/65535;
        else
            origch1=img(:,:,1)/65535;
            origch2=img(:,:,2)/65535;
        end
        ch3=img(:,:,3);
        ch3=ch3-ch3; %for 0,1,0 images
        countedimg=cat(3,ch3,ch3,ch3);
        andimg=cat(3,ch3,ch3,ch3);
        maxch1=max(max(origch1));
        maxch2=max(max(origch2));
        minc1=max(1/65535,mean(prctile(origch1,ch1p)));
        minc2=max(1/65535,mean(prctile(origch2,ch2p)));
        txt="Nothing yet";
        w3.Text=txt;
        if(minc1/maxch1<0.05 && minc2/maxch2<0.05)
            buttonon(but10x);%.Value=true;
        end
        ch1s.Limits=[0 maxch1];
        ch2s.Limits=[0 maxch2];
        if(but10x.Value)
            ch1s.Limits=[0 max(maxch1/10,minc1)];
            ch2s.Limits=[0 max(maxch2/10,minc2)];
        end
        ch1s.Value=minc1;
        ch2s.Value=minc2;
        buttonoff(butshowand);%.Value=false;
        buttonon(butch1);%.Value=true;
        buttonon(butch2);%.Value=true;
        drawnow
        ch1=im2bw(origch1,ch1s.Value(1));
        ch2=im2bw(origch2,ch2s.Value(1));
        sz=size(origch1,2)/7;
        x1=round(sz);
        x2=round(sz*2);
        x3=round(sz*3);
        x4=round(sz*4);
        x5=round(sz*5);
        x6=round(sz*6);
        ox1=x1;
        ox2=x2;
        ox3=x3;
        ox4=x4;
        ox5=x5;
        ox6=x6;
        zoomed=6;
        drawimg();
        contin=false;
        showui();
        while(contin==false && quit==false)
            y=round(image.Parent.CurrentPoint(3));
            x=round(image.Parent.CurrentPoint(1));
            switch zoomed
                case 1
                    x=x+x1;
                case 2
                    x=x+x2;
                case 3
                    x=x+x3;
                case 4
                    x=x+x4;
                case 5
                    x=x+x5;
            end
            minx=max(1,x-10);
            miny=max(1,y-10);
            maxx=min(x+10,size(origch1,2));
            maxy=min(y+10,size(origch1,1));
            c1=0;
            c2=0;
            n=0;
            for x=minx:maxx
                for y=miny:maxy
                    c1=c1+origch1(y,x);
                    c2=c2+origch2(y,x);
                    n=n+1;
                end
            end
            c1=c1/n;
            c2=c2/n;
            drawnow
            pause(0.1);
        end
        close all
        if quit
            close(u);
            return
        end
    end
end
xx=questdlg("Finished!","Finished!","NICE!","NICE!","NICE!");
close(u);
return

function updatebutbox(src,event)
    global but10x
    global ch1s
    global ch2s
    global maxch1;
    global maxch2;
    global image;
    
    global roibox;
    global roiwidth;
    global roiheight;
    global roix;
    global roiy;
    global x1;
    global x2;
    global zoomed;
    global x3;
    global x4;
    global x5;
    global finishedbox;
    finishedbox=false;
    blockui("Select region to erase. Double-click to accept the region; select box and press delete key to cancel.");
    roiy=round(image.Parent.XLim(1)+(image.Parent.XLim(2)-image.Parent.XLim(1))*0.4);
    roiheight=round((image.Parent.XLim(2)-image.Parent.XLim(1))*0.2);
    
    roix=round(image.Parent.YLim(1)+(image.Parent.YLim(2)-image.Parent.YLim(1))*0.4);
    roiwidth=round((image.Parent.YLim(2)-image.Parent.YLim(1))*0.2);
    
    roibox = images.roi.Rectangle(image.Parent,'Position',[roiy roix roiheight roiwidth], 'Deletable', true, 'InteractionsAllowed','all', 'MarkerSize', 10, 'Color',[0 1 0], 'Label', 'Adjust boundaries; double-click in box to clear');
    addlistener(roibox, 'MovingROI', @roifn);
    addlistener(roibox, 'ROIClicked', @roifn2);
    addlistener(roibox, 'DeletingROI', @roifn3);
    drawnow
    while finishedbox==false
        pause(0.1);
    end
    showui();
end

function roifn3(src,event)
    global finishedbox
    finishedbox=true;
end


function updatebutbox2(src,event)
    global but10x
    global ch1s
    global ch2s
    global maxch1;
    global maxch2;
    global image;
    
    global roibox;
    global roiwidth;
    global roiheight;
    global roix;
    global roiy;
    global x1;
    global x2;
    global zoomed;
    global x3;
    global x4;
    global x5;
    
    global finishedbox;
    finishedbox=false;
    blockui("Select region to crop. Double-click to accept the region; select box and press delete key to cancel.");
    
    roiy=round(image.Parent.XLim(1)+(image.Parent.XLim(2)-image.Parent.XLim(1))*0.4);
    roiheight=round((image.Parent.XLim(2)-image.Parent.XLim(1))*0.2);
    
    roix=round(image.Parent.YLim(1)+(image.Parent.YLim(2)-image.Parent.YLim(1))*0.4);
    roiwidth=round((image.Parent.YLim(2)-image.Parent.YLim(1))*0.2);
    
    roibox = images.roi.Rectangle(image.Parent,'Position',[roiy roix roiheight roiwidth], 'Deletable', true, 'InteractionsAllowed','all', 'MarkerSize', 10, 'Color',[0 1 0], 'Label', 'Adjust boundaries; double-click in box to crop');
    addlistener(roibox, 'MovingROI', @roifn);
    addlistener(roibox, 'ROIClicked', @roifn2a);
    addlistener(roibox, 'DeletingROI', @roifn3);
    drawnow
    while finishedbox==false
        pause(0.1);
    end
    showui();
end

function roifn(obj,event)
    global roiheight;
    global roix;
    global roiwidth;
    global roiy;
    roix=obj.Position(1);
    roiy=obj.Position(2);
    roiwidth=obj.Position(3);
    roiheight=obj.Position(4);
end

function roifn2(obj,event)
    global roiheight;
    global roix;
    global roiwidth;
    global roiy;
    global origch1;
    global origch2;
    global ch1;
    global ch2;
    global cansave;
    global butbox3;
    global zoomed;
    global x1;
    global x2;
    global x3;
    global x4;
    global x5;
    global finishedbox;
    global roibox;
    global giantbutton;
    if event.SelectionType=='double'
        finishedbox=true;
        roix=obj.Position(1);
        roiy=obj.Position(2);
        roiwidth=obj.Position(3);
        roiheight=obj.Position(4);
        giantbutton.Text="Please wait -- erasing";
        drawnow
        switch zoomed
            case 6
                roix=roix+0;
            case 1
                roix=roix+x1;
            case 2
                roix=roix+x2;
            case 3
                roix=roix+x3;
            case 4
                roix=roix+x4;
            case 5
                roix=roix+x5;
        end
        origch1(roiy:(roiy+roiheight),roix:(roix+roiwidth))=0;
        origch2(roiy:(roiy+roiheight),roix:(roix+roiwidth))=0;
        ch1(roiy:(roiy+roiheight),roix:(roix+roiwidth))=0;
        ch2(roiy:(roiy+roiheight),roix:(roix+roiwidth))=0;
        delete(roibox);
        drawimg();
        cansave=true;
        butbox3.Visible='on';
        drawnow
    end
end


function roifn2a(obj,event)
    global roiheight;
    global roix;
    global roiwidth;
    global roiy;
    global origch1;
    global origch2;
    global ch1;
    global ch2;
    global ch3;
    global zoomed;
    global countedimg;
    global andimg;
    global x1;
    global x2;
    global x3;
    global x4;
    global x5;
    global x6;
    global ox1;
    global ox6;
    global ox2;
    global ox3;
    global ox4;
    global ox5;
    global finishedbox;
    global giantbutton;
        global cansave;
    global butbox3;
    
    if event.SelectionType=='double'
        finishedbox=true;
        giantbutton.Text="Please wait -- cropping";
        drawnow
        roix=obj.Position(1);
        roiy=obj.Position(2);
        roiwidth=obj.Position(3);
        roiheight=obj.Position(4);
        switch zoomed
            case 6
                roix=roix+0;
            case 1
                roix=roix+x1;
            case 2
                roix=roix+x2;
            case 3
                roix=roix+x3;
            case 4
                roix=roix+x4;
            case 5
                roix=roix+x5;
        end
        y1=round(roiy);
        y2=round(roiy+roiheight);
        x1=round(roix);
        x2=round(roix+roiwidth);
        ch1=ch1(y1:y2,x1:x2);
        ch2=ch2(y1:y2,x1:x2);
        ch3=ch3(y1:y2,x1:x2);
        origch1=origch1(y1:y2,x1:x2);
        origch2=origch2(y1:y2,x1:x2);
        countedimg=countedimg(y1:y2,x1:x2,:);
        andimg=andimg(y1:y2,x1:x2,:);
        sz=size(origch1,2)/7;
        x1=round(sz);
        x2=round(sz*2);
        x3=round(sz*3);
        x4=round(sz*4);
        x5=round(sz*5);
        x6=round(sz*6);
        ox1=x1;
        ox2=x2;
        ox3=x3;
        ox4=x4;
        ox5=x5;
        ox6=x6;
    
        delete(obj);
        drawimg();
        cansave=true;
        butbox3.Visible='on';
        drawnow
    end
end

function updatebut10x(src,event)
    global but10x
    global ch1s
    global ch2s
    global maxch1;
    global maxch2;
    global origch1;
    global origch2;
    ch1s.Limits=[0 maxch1];
    ch2s.Limits=[0 maxch2];
    if(but10x.Value)
        ch1s.Limits=[0 max(maxch1/10,ch1s.Value)];
        ch2s.Limits=[0 max(maxch2/10,ch2s.Value)];
    end
    drawnow
end

function updateCh1(src,event)
    global ch1s
    global ch2s
    global origch1
    global ch1
    global w4
    global butch1;
    global butch2;
    global butshowand;
    global butreset;
    global image;
    
    blockui("Please wait");
    oldxaxes=image.Parent.XLim;
    oldyaxes=image.Parent.YLim;
    ch1s.Value=max(ch1s.Value,1/65535);
    ch1=im2bw(origch1,ch1s.Value(1));
    drawimg();
    image.Parent.XLim=oldxaxes;
    image.Parent.YLim=oldyaxes;
    showui();
end

function updateCh2(src,event)
    global ch2s
    global ch1s
    global origch2
    global ch2
    global w4;
    global butch1;
    global butch2;
    global butshowand;
    global butreset;
    global image;
    blockui("Please WAIT");
    oldxaxes=image.Parent.XLim;
    oldyaxes=image.Parent.YLim;
    ch2s.Value=max(ch2s.Value,1/65535);
    ch2=im2bw(origch2,ch2s.Value(1));
    drawimg();
    image.Parent.XLim=oldxaxes;
    image.Parent.YLim=oldyaxes;
    showui();
end

function updatebutsetb(src,event)
    global ch1
    global c1
    global c2
    global ch2s
    global ch1s
    global origch2
    global ch2
    global w4;
    global butch1;
    global butch2;
    global butshowand;
    global butreset;
    global but10x;
    global image;
    global maxch1;
    global maxch2;
    
    blockui("Please WAIT");
    oldxaxes=image.Parent.XLim;
    oldyaxes=image.Parent.YLim;
    ch1fac=1.3;
    ch2fac=1.3;
    d=1;
    if(but10x.Value)
        d=10;
    end
    n1=max(1/65535,min(ch1fac*c1,maxch1));
    n2=max(1/65535,min(ch2fac*c2,maxch2));
    ch1s.Limits=[0 max(maxch1/d,ch1fac*c1)];
    ch2s.Limits=[0 max(maxch2/d,ch2fac*c2)];
    ch1s.Value(1)=n1;
    ch2s.Value(1)=n2;
    ch2=im2bw(origch2,ch2s.Value(1));
    ch1=im2bw(origch2,ch1s.Value(1));
    drawimg();
    image.Parent.XLim=oldxaxes;
    image.Parent.YLim=oldyaxes;
    showui();
end


function updateCh1b(src,event)
    global butch1;
    global butch2;
    global butshowand;
    global butshowcounted;
    global w4;
    global image;
    
    blockui("Please WAIT");
    if(butch1.Value==true)
        buttonon(butch1);
        buttonoff(butshowand);%.Value=false;
        buttonoff(butshowcounted);%.Value=false;
    end
    if(butch1.Value==false)
        buttonoff(butch1);
        if(butch2.Value==false)
            buttonon(butch1);%.Value=true;
        end
    end
    drawnow
    oldxaxes=image.Parent.XLim;
    oldyaxes=image.Parent.YLim;
    drawimg();
    image.Parent.XLim=oldxaxes;
    image.Parent.YLim=oldyaxes;
    showui();
end

function updateCh2b(src,event)
    global butch1;
    global butch2;
    global butshowand;
    global butshowcounted;
    global w4;
    global image;
    
    blockui("Please WAIT");
    if(butch2.Value==true)
        buttonon(butch1);
        buttonoff(butshowand);%.Value=false;
        buttonoff(butshowcounted);%.Value=false;
    end
    if(butch2.Value==false)
        buttonoff(butch2);
        if(butch1.Value==false)
            buttonon(butch2);%.Value=true;
        end
    end
    drawnow
    oldxaxes=image.Parent.XLim;
    oldyaxes=image.Parent.YLim;
    drawimg();
    image.Parent.XLim=oldxaxes;
    image.Parent.YLim=oldyaxes;
    showui();
end

function updatebutshowand(src,event)
    global butshowand;
    global butshowcounted;
    global butch1;
    global butch2;
    global w4;
    global image;
    
    blockui("Please WAIT");
    
    if(butshowand.Value==true)
        buttonoff(butshowcounted);
        buttonoff(butch1);
        buttonoff(butch2);
    end
    if(butshowand.Value==false)
        buttonoff(butshowcounted);
        buttonon(butch1);
        buttonon(butch2);
    end
    drawnow
    oldxaxes=image.Parent.XLim;
    oldyaxes=image.Parent.YLim;
    drawimg();
    image.Parent.XLim=oldxaxes;
    image.Parent.YLim=oldyaxes;
    showui();
end

function updatebutshowcounted(src,event)
    global butshowand;
    global butshowcounted;
    global butch1;
    global butch2;
    global w4;
    global image;
    
    blockui("Please WAIT");
    
    if(butshowcounted.Value==true)
        buttonoff(butshowand);
        buttonoff(butch1);
        buttonoff(butch2);
    end
    if(butshowcounted.Value==false)
        buttonoff(butshowand);
        buttonon(butch1);
        buttonon(butch2);
    end
    drawnow
    oldxaxes=image.Parent.XLim;
    oldyaxes=image.Parent.YLim;
    drawimg();
    image.Parent.XLim=oldxaxes;
    image.Parent.YLim=oldyaxes;
    showui();
end


function updatebutreset(src,event)
    global butshowand;
    global ch1s;
    global ch2s;
    global ch1;
    global ch2;
    global origch1;
    global origch2;
    global w4;
    global maxch1;
    global maxch2;
    global butch1;
    global butch2;
    global butshowand;
    global butshowcounted;
    global butreset;
    global minc1;
    global minc2;
    global ch1p;
    global ch2p;
    global butgaussianblur;
    blockui("Please WAIT");
    
    if(butgaussianblur.Value)
        origch1=img(:,:,1)/65535;
        origch2=img(:,:,2)/65535;
        ch1=im2bw(origch1,ch1s.Value(1));
        ch2=im2bw(origch2,ch2s.Value(1));
        buttonoff(butgaussianblur);
    end
    %ch1s.Value=0.05*maxch1;
    %ch2s.Value=0.05*maxch2;
    maxch1=max(max(origch1));
    maxch2=max(max(origch2));
    minc1=mean(prctile(origch1,ch1p));
    minc2=mean(prctile(origch2,ch2p));
    if(minc1/maxch1<0.05 && minc2/maxch2<0.05)
        buttonon(but10x);
    else
        buttonoff(but10x);
    end
    ch1s.Limits=[0 maxch1];
    ch2s.Limits=[0 maxch2];
    if(but10x.Value)
        ch1s.Limits=[0 max(maxch1/10,minc1)];
        ch2s.Limits=[0 max(maxch2/10,minc2)];
    end
    minc1=max(1/65535,minc1);
    minc2=max(1/65535,minc2);
    ch1s.Value(1)=minc1;
    ch2s.Value(1)=minc2;
    buttonoff(butshowand);
    buttonoff(butshowcounted);
    buttonon(butch1);
    buttonon(butch2);
    drawnow
    ch2=im2bw(origch2,ch2s.Value(1));
    ch1=im2bw(origch1,ch1s.Value(1));
    drawimg();
    showui();
end

function lineMoving1(src,event)
    global line1;
    global line2;
    global line3;
    global line4;
    global line5;
    global line6;
    global x1;
    global x2;
    global x3;
    global x4;
    global x5;
    global x6;
    
    x1=round(line1.Position(1));
    x2=round(line2.Position(1));
    x3=round(line3.Position(1));
    x4=round(line4.Position(1));
    x5=round(line5.Position(1));
    x6=round(line6.Position(1));
end


function updatebutcount(src,event)
    global butshowand;
    global ch1s;
    global x1;
    global x2;
    global x3;
    global x4;
    global x5;
    global x6;
    global ox1;
    global ox2;
    global ox3;
    global ox4;
    global ox5;
    global ox6;
    global ch2s;
    global grid;
    global ch1;
    global ch2;
    global butch1;
    global butch2;
    global butshowand;
    global butshowcounted;
    global countedall;
    global countedimg;
    global zoomed;
    global countedr;
    global butreset;
    global counts;
    global txt;
    global newname;
    global w4;
    global stats1;
    global stats2;
    global stats3;
    global stats4;
    global stats5;
    global index1;
    global index2;
    global index3;
    global index4;
    global index5;
    global comment;
    global allstats;
    global allindices;
    global fullfilename;
    global parentdir;
    global justfilename;
    global w3;
    global andimg;
    global filesep
    filesep='\';
    if(x1==ox1 || x6==ox6)
        xx=questdlg("Boundaries have not been changed.","Boundaries have not been changed.","Count anyway","Go back","Go back");
        if xx=="Go back"
            return
        end
    end
    blockui("Please WAIT");
    
    drawnow
    andimg=and(ch1,ch2);
    txt=sprintf("Counted items:\tR1\tR2\tR3\tR4\tR5\n");
    
    for t=1:3
        if t==1
            i=ch1;
            prefx='DAPI';
        end
        if t==2
            i=ch2;
            prefx='GFP';
        end
        if t==3
            i=andimg;
            prefx='BOTH';
        end
    
        seg1=i(:,x1:(x2-1));
        seg2=i(:,x2:(x3-1));
        seg3=i(:,x3:(x4-1));
        seg4=i(:,x4:(x5-1));
        seg5=i(:,x5:(x6));
    
        minSize=150;
        maxSize=10000;
        maxLength=300;
        minEcc=0;
        maxEcc=0.99;
    
        if(zoomed==6 || zoomed==1)
            stats1=regionprops(seg1,{'Centroid','Area','Eccentricity','PixelList','MajorAxisLength'});
            index1 = find([stats1.Area] >= minSize & [stats1.Area] <= maxSize & [stats1.MajorAxisLength]<=maxLength & [stats1.Eccentricity]>=minEcc & [stats1.Eccentricity]<=maxEcc);
            p=1;
            allstats{p,t}=stats1;
            allindices{p,t}=index1;
            counts(1,t)=size(index1,2);
            countedr(1)=true;
            PixelList = cell(length(index1));
            for j=1:length(index1)
                PixelList{j} = stats1(index1(j)).PixelList;
                for k = 1:size(PixelList{j},1)
                    countedimg(round(PixelList{j}(k,2)),round(PixelList{j}(k,1)+x1-1),t)=1;
                end
            end
        end
        if(zoomed==6 || zoomed==2)
            stats2=regionprops(seg2,{'Centroid','Area','Eccentricity','PixelList','MajorAxisLength'});
            index2 = find([stats2.Area] >= minSize & [stats2.Area] <= maxSize & [stats2.MajorAxisLength]<=maxLength & [stats2.Eccentricity]>=minEcc & [stats2.Eccentricity]<=maxEcc);
            p=2;
            allstats{p,t}=stats2;
            allindices{p,t}=index2;
            counts(2,t)=size(index2,2);
            countedr(2)=true;
            PixelList = cell(length(index2));
            for j=1:length(index2)
                PixelList{j} = stats2(index2(j)).PixelList;
                for k = 1:size(PixelList{j},1)
                    countedimg(round(PixelList{j}(k,2)),round(PixelList{j}(k,1)+x2-1),t)=1;
                end
            end
        end
        if(zoomed==6 || zoomed==3)
            stats3=regionprops(seg3,{'Centroid','Area','Eccentricity','PixelList','MajorAxisLength'});
            index3 = find([stats3.Area] >= minSize & [stats3.Area] <= maxSize & [stats3.MajorAxisLength]<=maxLength & [stats3.Eccentricity]>=minEcc & [stats3.Eccentricity]<=maxEcc);
            p=3;
            allstats{p,t}=stats3;
            allindices{p,t}=index3;
            counts(3,t)=size(index3,2);
            countedr(3)=true;
            PixelList = cell(length(index3));
            for j=1:length(index3)
                PixelList{j} = stats3(index3(j)).PixelList;
                for k = 1:size(PixelList{j},1)
                    countedimg(round(PixelList{j}(k,2)),round(PixelList{j}(k,1)+x3-1),t)=1;
                end
            end
        end
        if(zoomed==6 || zoomed==4)
            stats4=regionprops(seg4,{'Centroid','Area','Eccentricity','PixelList','MajorAxisLength'});
            index4 = find([stats4.Area] >= minSize & [stats4.Area] <= maxSize & [stats4.MajorAxisLength]<=maxLength & [stats4.Eccentricity]>=minEcc & [stats4.Eccentricity]<=maxEcc);
            counts(4,t)=size(index4,2);
            countedr(4)=true;
            p=4;
            allstats{p,t}=stats4;
            allindices{p,t}=index4;
            PixelList = cell(length(index4));
            for j=1:length(index4)
                PixelList{j} = stats4(index4(j)).PixelList;
                for k = 1:size(PixelList{j},1)
                    countedimg(round(PixelList{j}(k,2)),round(PixelList{j}(k,1)+x4-1),t)=1;
                end
            end
        end
        if(zoomed==6 || zoomed==5)
            stats5=regionprops(seg5,{'Centroid','Area','Eccentricity','PixelList','MajorAxisLength'});
            index5 = find([stats5.Area] >= minSize & [stats5.Area] <= maxSize & [stats5.MajorAxisLength]<=maxLength & [stats5.Eccentricity]>=minEcc & [stats5.Eccentricity]<=maxEcc);
            p=5;
            allstats{p,t}=stats5;
            allindices{p,t}=index5;
            counts(5,t)=size(index5,2);
            countedr(5)=true;
            PixelList = cell(length(index5));
            for j=1:length(index5)
                PixelList{j} = stats5(index5(j)).PixelList;
                for k = 1:size(PixelList{j},1)
                    countedimg(round(PixelList{j}(k,2)),round(PixelList{j}(k,1)+x5-1),t)=1;
                end
            end
    
        end
        countedr(6)= countedr(1) && countedr(2) && countedr(3) && countedr(4) && countedr(5);
        txt=sprintf("%s%s: %.0f %.0f %.0f %.0f %.0f\n",txt,prefx,counts(1,t),counts(2,t),counts(3,t),counts(4,t),counts(5,t));
    end
    cm=comment.Value;
    c1v=ch1s.Value;
    c2v=ch2s.Value;
    save(newname,"allstats","allindices","x1","x2","x3","x4","x5","x6","minSize","maxSize","maxLength","minEcc","maxEcc","cm","counts","c1v","c2v"); %,"ch1s.Value(1)","ch2s.Value(2)");
    w3.Text=txt;
    if ~isdir(strcat(parentdir,filesep,"Counted"))
        mkdir(strcat(parentdir,filesep,"Counted"));
    end
    imwrite(countedimg, strcat(parentdir,filesep,"Counted",filesep,"COUNTED ",justfilename));
    showui();
    butshowcounted.Visible='on';
    drawnow
end


function updatebutnext(src,event)
    global contin;
    global countedr;
    global fullfilename;
    global parentdir
    global justfilename
    global imgname;
    global filesep;
    filesep='\';
    contin=false;
    if(~countedr(6))
        xx=questdlg("Counting is not complete.","Counting is not complete.","Skip this one","Count and proceed","Stay","Stay");
        if xx=="Stay"
            return
        end
        if xx=="Count and proceed"
            zoomed=6;
            updatebutcount();
        end
        if xx=="Skip this one"
            yy=questdlg("Move the file to skip always?","Move the file to skip always?","Yes","No","Yes");
            if yy=="Yes"
                if ~isdir(strcat(parentdir,filesep,"Skipped"))
                    mkdir(strcat(parentdir,filesep,"Skipped"));
                end
                movefile(fullfilename, strcat(parentdir,filesep,"Skipped",filesep,justfilename));
            end
        end
    end
    contin=true;
end

function updatebutbox3(src,event)
    global contin;
    global countedr;
    global fullfilename;
    global filesep
    global parentdir
    global justfilename
    global imgname;
    global origch1;
    global origch2;
    global cansave;
    global butbox3;
    global ch3;
    blockui(strcat("Saving image ",imgname));
    if ~isdir(strcat(parentdir,filesep,"Originals"))
        mkdir(strcat(parentdir,filesep,"Originals"));
    end
    if(~isfile(strcat(parentdir,filesep,"Originals",filesep,justfilename)))
        movefile(strcat(parentdir,filesep,justfilename), strcat(parentdir,filesep,"Originals",filesep,justfilename));
    end
    img=cat(3,uint16(origch1*65535),uint16(origch2*65535),uint16(ch3*65535));
    imwrite(img,imgname);
    butbox3.Visible='off';
    cansave=false;
    showui();
end


function updatebutquit(src,event)
    global quit;
    xx=questdlg("Quit?","Quit?","Yes","Oops, no","Oops, no");
    quit=false;
    if xx=="Yes"
        quit=true;
    end
end

function updatebutreverse(src,event)
    global reversed;
    global origch1;
    global origch2;
    global ch1;
    global ch2;
    global butreverse;
    global w4;
    global butbox3;
    global cansave;
    cansave=true;
    reversed=~reversed;
    blockui("Please WAIT");
    
    drawnow
    origch1=flipdim(origch1,2);
    origch2=flipdim(origch2,2);
    ch1=flipdim(ch1,2);
    ch2=flipdim(ch2,2);
    drawimg();
    if(reversed)
        buttonon(butreverse);
    else
        buttonoff(butreverse);
    end
    showui();
    butbox3.Visible='on';
    drawnow
end

function drawimg
    global ch2;
    global ch3;
    global ch1;
    global image;
    global newimg;
    global shownimg;
    global butch1;
    global butch2;
    global butshowand;
    global butshowcounted;
    global countedimg;
    global line1;
    global line2;
    global line3;
    global line4;
    global countedr;
    global line5;
    global line6;
    global x1;
    global x2;
    global x3;
    global x4;
    global x5;
    global x6;
    global zoomed;
    
    if(butshowand.Value==true)
        andimg=and(ch1,ch2);
        shownimg=andimg;
    end
    if(butshowcounted.Value==true)
        if(countedr(zoomed))
            shownimg=countedimg;
        else
            buttonoff(butshowcounted);
            buttonon(butch1);
            buttonon(butch2);
            drawnow
        end
    end
    if(butshowand.Value==false && butshowcounted.Value == false)
        if(butch1.Value==true && butch2.Value==true)
            newimg=cat(3,ch1,ch2,ch3);
            shownimg=newimg;
        end
        if(butch1.Value==true && butch2.Value==false)
            newimg=cat(3,ch1,ch3,ch3);
            shownimg=newimg;
        end
        if(butch1.Value==false && butch2.Value==false)
            newimg=cat(3,ch3,ch3,ch3);
            shownimg=newimg;
        end
        if(butch1.Value==false && butch2.Value==true)
            newimg=cat(3,ch3,ch2,ch3);
            shownimg=newimg;
        end
    end
    switch zoomed
        case 6
            image=imshow(shownimg);
            line1 = images.roi.Line(image.Parent, 'Position',[x1 x1; image.Parent.YLim(1) image.Parent.YLim(2)]','InteractionsAllowed','translate','MarkerSize',5,'Color',[1 0 0]);
            line2 = images.roi.Line(image.Parent, 'Position',[x2 x2; image.Parent.YLim(1) image.Parent.YLim(2)]','InteractionsAllowed','translate','MarkerSize',5,'Color',[1 0.5 0]);
            line3 = images.roi.Line(image.Parent, 'Position',[x3 x3; image.Parent.YLim(1) image.Parent.YLim(2)]','InteractionsAllowed','translate','MarkerSize',5,'Color',[1 1 0]);
            line4 = images.roi.Line(image.Parent, 'Position',[x4 x4; image.Parent.YLim(1) image.Parent.YLim(2)]','InteractionsAllowed','translate','MarkerSize',5,'Color',[0 1 0]);
            line5 = images.roi.Line(image.Parent, 'Position',[x5 x5; image.Parent.YLim(1) image.Parent.YLim(2)]','InteractionsAllowed','translate','MarkerSize',5,'Color',[0 1 1]);
            line6 = images.roi.Line(image.Parent, 'Position',[x6 x6; image.Parent.YLim(1) image.Parent.YLim(2)]','InteractionsAllowed','translate','MarkerSize',5,'Color',[1 0 1]);
            addlistener(line1, 'MovingROI', @lineMoving1);
            addlistener(line2, 'MovingROI', @lineMoving1);
            addlistener(line3, 'MovingROI', @lineMoving1);
            addlistener(line4, 'MovingROI', @lineMoving1);
            addlistener(line5, 'MovingROI', @lineMoving1);
            addlistener(line6, 'MovingROI', @lineMoving1);
        case 1
            image=imshow(shownimg(:,x1:(x2-1),:));
        case 2
            image=imshow(shownimg(:,x2:(x3-1),:));
        case 3
            image=imshow(shownimg(:,x3:(x4-1),:));
        case 4
            image=imshow(shownimg(:,x4:(x5-1),:));
        case 5
            image=imshow(shownimg(:,x5:(x6-1),:));
    end
end


function updatebutfull(src,event)
    global w4;
    global zoomed;
    global butfull;
    global butr1;
    global butr2;
    global butr3;
    global butr4;
    global butr5;
    global butcount;
    global butshowcounted;
    global countedr;
    global butch1;
    global butch2;
    blockui("Please WAIT");
    
    buttonon(butfull);
    buttonoff(butr1);
    buttonoff(butr2);
    buttonoff(butr3);
    buttonoff(butr4);
    buttonoff(butr5);
    butcount.Text="Count all and save";
    zoomed=6;
    butshowcounted.Visible='off';
    if countedr(zoomed)
        butshowcounted.Visible='on';
    end
    if(butshowcounted.Value==true && countedr(zoomed)==false)
        buttonoff(butshowcounted);
        butshowcounted.Visible='off';
        buttonon(butch1);
        buttonon(butch2);
    end
    drawnow
    drawimg();
    showui();
    drawnow
end

function updatebutr1(src,event)
    global w4;
    global zoomed;
    global butfull;
    global butr1;
    global butr2;
    global butr3;
    global butr4;
    global butr5;
    global butcount;
    global butshowcounted;
    global countedr;
    global butch1;
    global butch2;
    global image;
    blockui("Please WAIT");
    
    buttonoff(butfull);
    buttonon(butr1);
    buttonoff(butr2);
    buttonoff(butr3);
    buttonoff(butr4);
    buttonoff(butr5);
    zoomed=1;
    butshowcounted.Visible='off';
    if countedr(zoomed)
        butshowcounted.Visible='on';
    end
    if(butshowcounted.Value==true && countedr(zoomed)==false)
        buttonoff(butshowcounted);
        butshowcounted.Visible='off';
buttonon(butch1);
buttonon(butch2);
    end
    butcount.Text="Count R1 and save";
    drawnow
    drawimg();
    showui();
    drawnow
end

function updatebutr2(src,event)
    global w4;
    global zoomed;
    global butfull;
    global butr1;
    global butr2;
    global butr3;
    global butr4;
    global butr5;
    global butcount;
    global butshowcounted;
    global countedr;
    global butch1;
    global butch2;
    blockui("Please WAIT");
    
    buttonoff(butfull);
    buttonoff(butr1);
    buttonon(butr2);
    buttonoff(butr3);
    buttonoff(butr4);
    buttonoff(butr5);
    zoomed=2;
    butshowcounted.Visible='off';
    if countedr(zoomed)
        butshowcounted.Visible='on';
    end
    if(butshowcounted.Value==true && countedr(zoomed)==false)
        buttonoff(butshowcounted);
        butshowcounted.Visible='off';
buttonon(butch1);
buttonon(butch2);
    end
    
    butcount.Text="Count R2 and save";
    drawnow
    drawimg();
    showui();
    drawnow
end

function updatebutr3(src,event)
    global w4;
    global zoomed;
    global butfull;
    global butr1;
    global butr2;
    global butr3;
    global butr4;
    global butr5;
    global butcount;
    global butshowcounted;
    global countedr;
    global butch1;
    global butch2;
    blockui("Please WAIT");
    
    buttonoff(butfull);
    buttonoff(butr1);
    buttonoff(butr2);
    buttonon(butr3);
    buttonoff(butr4);
    buttonoff(butr5);
    zoomed=3;
    butshowcounted.Visible='off';
    if countedr(zoomed)
        butshowcounted.Visible='on';
    end
    if(butshowcounted.Value==true && countedr(zoomed)==false)
        buttonoff(butshowcounted);
        butshowcounted.Visible='off';
buttonon(butch1);
buttonon(butch2);
    end
    butcount.Text="Count R3 and save";
    drawnow
    drawimg();
    showui();
    drawnow
end

function updatebutr4(src,event)
    global w4;
    global zoomed;
    global butfull;
    global butr1;
    global butr2;
    global butr3;
    global butr4;
    global butr5;
    global butcount;
    global butshowcounted;
    global countedr;
    global butch1;
    global butch2;
    blockui("Please WAIT");
    buttonoff(butfull);
    buttonoff(butr1);
    buttonoff(butr2);
    buttonoff(butr3);
    buttonon(butr4);
    buttonoff(butr5);
    zoomed=4;
    butshowcounted.Visible='off';
    if countedr(zoomed)
        butshowcounted.Visible='on';
    end
    if(butshowcounted.Value==true && countedr(zoomed)==false)
        buttonoff(butshowcounted);
        butshowcounted.Visible='off';
buttonon(butch1);
buttonon(butch2);
    end
    butcount.Text="Count R4 and save";
    drawnow
    drawimg();
    showui();
    drawnow
end

function updatebutr5(src,event)
    global w4;
    global zoomed;
    global butfull;
    global butr1;
    global butr2;
    global butr3;
    global butr4;
    global butr5;
    global butcount;
    global butshowcounted;
    global countedr;
    global butch1;
    global butch2;
    blockui("Please WAIT");
    buttonoff(butfull);
    buttonoff(butr1);
    buttonoff(butr2);
    buttonoff(butr3);
    buttonoff(butr4);
    buttonon(butr5);
    zoomed=5;
    butshowcounted.Visible='off';
    if countedr(zoomed)
        butshowcounted.Visible='on';
    end
    if(butshowcounted.Value==true && countedr(zoomed)==false)
        buttonoff(butshowcounted);
        butshowcounted.Visible='off';
buttonon(butch1);
buttonon(butch2);
    end
    butcount.Text="Count R5 and save";
    drawnow
    drawimg();
    showui();
    drawnow
end


function updatebutgaussianblur(src,event)
    global w4;
    global origch1;
    global ch1s;
    global ch2s;
    global ch1;
    global ch2;
    global origch2;
    global image;
    global butgaussianblur;
    global imgname;
    global img;
    blockui("Please WAIT");   
    drawnow
    oldxaxes=image.Parent.XLim;
    oldyaxes=image.Parent.YLim;
    if(butgaussianblur.Value)
        origch1=imgaussfilt(origch1,1,"FilterSize",3);
        origch2=imgaussfilt(origch2,1,"FilterSize",3);
        ch1=im2bw(origch1,ch1s.Value(1));
        ch2=im2bw(origch2,ch2s.Value(1));
    else
        origch1=img(:,:,1)/65535;
        origch2=img(:,:,2)/65535;
        ch1=im2bw(origch1,ch1s.Value(1));
        ch2=im2bw(origch2,ch2s.Value(1));
    end
    drawimg();
    image.Parent.XLim=oldxaxes;
    image.Parent.YLim=oldyaxes;
    showui();
    drawnow
end


function updatebutc1decdec(src,event)
    global ch2s
    global ch1s
    global origch2
    global origch1;
    global ch1;
    global ch2
    global w4;
    global butch1;
    global butch2;
    global image;
    global minc1;
    global minc2;
    blockui("Please WAIT");
    drawnow
    oldxaxes=image.Parent.XLim;
    oldyaxes=image.Parent.YLim;
    ch1s.Value=0.8*ch1s.Value;
    ch1s.Value=max(ch1s.Value,1/65535);
    ch1=im2bw(origch1,ch1s.Value(1));
    drawimg();
    image.Parent.XLim=oldxaxes;
    image.Parent.YLim=oldyaxes;
    showui();
    drawnow
end

function updatebutc1dec(src,event)
    global ch2s
    global ch1s
    global origch2
    global origch1;
    global ch2;
    global ch1;
    global w4;
    global butch1;
    global butch2;
    global image;
    blockui("Please WAIT");
    drawnow
    oldxaxes=image.Parent.XLim;
    oldyaxes=image.Parent.YLim;
    ch1s.Value=0.98*ch1s.Value;
    ch1s.Value=max(ch1s.Value,1/65535);
    ch1=im2bw(origch1,ch1s.Value(1));
    drawimg();
    image.Parent.XLim=oldxaxes;
    image.Parent.YLim=oldyaxes;
    showui();
    drawnow
end

function updatebutc1incinc(src,event)
    global ch2s
    global ch1s
    global origch2
    global origch1
    global ch2
    global ch1;
    global w4;
    global butch1;
    global butch2;
    global image;
    global maxch1;
    global maxch2;
    global minc1;
    global minc2;
    blockui("Please WAIT");
    drawnow
    oldxaxes=image.Parent.XLim;
    oldyaxes=image.Parent.YLim;
    newval=min(maxch1,1.2*ch1s.Value);
    ch1s.Limits(2)=max(ch1s.Limits(2),newval);
    ch1s.Value=newval;
    ch1=im2bw(origch1,ch1s.Value(1));
    drawimg();
    image.Parent.XLim=oldxaxes;
    image.Parent.YLim=oldyaxes;
    showui();
    drawnow
end

function updatebutc1inc(src,event)
    global ch2s
    global ch1s
    global origch2
    global origch1
    global ch2
    global ch1;
    global w4;
    global butch1;
    global butch2;
    global image;
    global maxch1;
    global maxch2;
    blockui("Please WAIT");
    drawnow
    oldxaxes=image.Parent.XLim;
    oldyaxes=image.Parent.YLim;
    newval=min(maxch1,1.02*ch1s.Value);
    ch1s.Limits(2)=max(ch1s.Limits(2),newval);
    ch1s.Value=newval;
    ch1=im2bw(origch1,ch1s.Value(1));
    drawimg();
    image.Parent.XLim=oldxaxes;
    image.Parent.YLim=oldyaxes;
    showui();
    drawnow
end

function updatebutc1reset(src,event)
    global ch2s
    global ch1s
    global origch2
    global origch1
    global ch2
    global ch1;
    global w4;
    global butch1;
    global butch2;
    global image;
    global maxch1;
    global maxch2;
    global ch1p;
    global ch2p;
    global minc1;
    global minc2;
    blockui("Please WAIT");
    drawnow
    oldxaxes=image.Parent.XLim;
    oldyaxes=image.Parent.YLim;
    minc1=max(1/65535,mean(prctile(origch1,ch1p)));
    if(minc1/maxch1<0.05 && minc2/maxch2<0.05)
        buttonon(but10x);
    else
        buttonoff(but10x);
    end
    ch1s.Limits=[0 maxch1];
    ch2s.Limits=[0 maxch2];
    if(but10x.Value)
        ch1s.Limits=[0 max(maxch1/10,minc1)];
    end
    ch1s.Value=minc1;
    ch1=im2bw(origch1,ch1s.Value(1));
    drawimg();
    image.Parent.XLim=oldxaxes;
    image.Parent.YLim=oldyaxes;
    showui();
    drawnow
end

function updatebutc2decdec(src,event)
    global ch2s
    global ch1s
    global origch2
    global origch1;
    global ch1;
    global ch2
    global w4;
    global minc1;
    global minc2;
    global butch1;
    global butch2;
    global image;
    blockui("Please WAIT");
    drawnow
    oldxaxes=image.Parent.XLim;
    oldyaxes=image.Parent.YLim;
    ch2s.Value=0.8*ch2s.Value;
    ch2s.Value=max(ch2s.Value,1/65535);
    ch2=im2bw(origch2,ch2s.Value(1));
    drawimg();
    image.Parent.XLim=oldxaxes;
    image.Parent.YLim=oldyaxes;
    showui();
    drawnow
end

function updatebutc2dec(src,event)
    global ch2s
    global ch1s
    global origch2
    global origch1;
    global ch2;
    global ch1;
    global w4;
    global butch1;
    global butch2;
    global image;
    blockui("Please WAIT");
    drawnow
    oldxaxes=image.Parent.XLim;
    oldyaxes=image.Parent.YLim;
    ch2s.Value=0.98*ch2s.Value;
    ch2s.Value=max(ch2s.Value,1/65535);
    ch2=im2bw(origch2,ch2s.Value(1));
    drawimg();
    image.Parent.XLim=oldxaxes;
    image.Parent.YLim=oldyaxes;
    showui();
    drawnow
end

function updatebutc2incinc(src,event)
    global ch2s
    global ch1s
    global origch2
    global origch1
    global ch2
    global ch1;
    global w4;
    global butch1;
    global butch2;
    global image;
    global maxch1;
    global maxch2;
    global minc1;
    global minc2;
    blockui("Please WAIT");
    drawnow
    oldxaxes=image.Parent.XLim;
    oldyaxes=image.Parent.YLim;
    newval=min(maxch2,1.2*ch2s.Value);
    ch2s.Limits(2)=max(ch2s.Limits(2),newval);
    ch2s.Value=newval;
    ch2=im2bw(origch2,ch2s.Value(1));
    drawimg();
    image.Parent.XLim=oldxaxes;
    image.Parent.YLim=oldyaxes;
    showui();
drawnow
end

function updatebutc2inc(src,event)
    global ch2s
    global ch1s
    global origch2
    global origch1
    global ch2
    global ch1;
    global w4;
    global butch1;
    global butch2;
    global image;
    global maxch1;
    global maxch2;
    blockui("Please WAIT");
    drawnow
    oldxaxes=image.Parent.XLim;
    oldyaxes=image.Parent.YLim;
    newval=min(maxch2,1.02*ch2s.Value);
    ch2s.Limits(2)=max(ch2s.Limits(2),newval);
    ch2s.Value=newval;
    ch2=im2bw(origch2,ch2s.Value(1));
    drawimg();
    image.Parent.XLim=oldxaxes;
    image.Parent.YLim=oldyaxes;
    showui();
    drawnow
end

function updatebutc2reset(src,event)
    global ch2s
    global ch1s
    global origch2
    global origch1
    global ch2
    global ch1;
    global w4;
    global butch1;
    global butch2;
    global minc1;
    global minc2;
    global image;
    global maxch1;
    global maxch2;
    global ch1p;
    global ch2p;
    global minc1;
    global minc2;
    blockui("Please WAIT");
    drawnow
    oldxaxes=image.Parent.XLim;
    oldyaxes=image.Parent.YLim;
    minc2=max(1/65535,mean(prctile(origch2,ch2p)));
    ch2s.Limits(2)=max(ch2s.Limits(2),minc2);
    ch2s.Value=minc2;
    ch2=im2bw(origch2,ch2s.Value(1));
    drawimg();
    image.Parent.XLim=oldxaxes;
    image.Parent.YLim=oldyaxes;
    showui();
    drawnow
end

function blockui(txt2)
    global giantbutton
    global grid
    global alreadyblocked

    if(alreadyblocked)
        showui();
    end
    giantbutton=uibutton(grid,'Text',txt2);
    giantbutton.Layout.Row=[1 7];
    giantbutton.Layout.Column=[1 9];
    alreadyblocked=true;
    drawnow
end

function showui
    global w4
    global giantbutton
    global alreadyblocked;
    alreadyblocked=false;
    w4.Text="Ready";
    try
        delete(giantbutton);
    catch SDFGSDF
    end
    drawnow
end

function buttonoff(button)
    global origbuttoncolor;
    button.Value=false;
    if(button.Text(length(button.Text)) == char(10003))
       button.Text=button.Text(1:length(button.Text)-2); 
    end
    button.BackgroundColor=origbuttoncolor;
    button.FontColor="#000000";
end

function buttonon(button)
    global kured;
    button.Value=true;
    button.FontColor="#ffffff";
    if(button.Text(length(button.Text)) ~= char(10003))
        button.Text=strcat(button.Text," ", char(10003));
    end
    button.BackgroundColor=kured;
end
```
