## Supplemental code 2 for "Gut hormone signaling drives sex differences in metabolism and behavior": makecsv.m

```
clear all
close all
fclose all
global parentdir
parentdir="D:\Rewitz lab\max";
global filesep
filesep='\';
fs=dir(parentdir);
fn=size(fs,1);
fo=fopen(strcat(parentdir,filesep,"GutAnalysis1.csv"),"w");
fo2=fopen(strcat(parentdir,filesep,"GutAnalysis2 all positions.csv"),"w");
terms={"DAPI","GFP","Both"};
fprintf(fo,"Name (commas replaced by exclamation points),R1Start,R1Len,R2Start,R2Len,R3Start,R3Len,R4Start,R4Len,R5Start,R5Len,R5End,TotalLen,R1DAPI,R1GFP,R1Both,R2DAPI,R2GFP,R2Both,R3DAPI,R3GFP,R3Both,R4DAPI,R4GFP,R4Both,R5DAPI,R5GFP,R5Both,TotalDAPI,TotalGFP,TotalBoth,Cells/Len,GFPCells/Len,GFP/Cells,Comment,,");
for c=1:3
    for p=1:5
        fprintf(fo,"%s quintile %i,",terms{c},p);
    end
    fprintf(fo,",");
end

for c=1:3
    for p=1:10
        fprintf(fo,"%s decile %i,",terms{c},p);
    end
    fprintf(fo,",");
end

for c=1:3
    for p=1:100
        fprintf(fo,"%s pctile %i,",terms{c},p);
    end
    fprintf(fo,",");
end

    for p=1:5
        fprintf(fo,"GFP enr quintile %i,",p);
    end
    fprintf(fo,",");

    for p=1:10
        fprintf(fo,"GFP enr decile %i,",p);
    end
    fprintf(fo,",");

    for p=1:100
        fprintf(fo,"GFP enr percentile %i,",p);
    end
    fprintf(fo,",");


    fprintf(fo,"\n");
ndone=0;
offsets=zeros(5,1);
prev="";
for t=3:fn
    f=fs(t).name;
    skip=false;
    suffix=f( (size(f,2)-3)  :   size(f,2)    );
    if(suffix == ".mat")
        load(strcat(parentdir,filesep,f));
        ndone=ndone+1;
        prev2=prev;
        prev=f;
        if ndone==1
            prev2=f;
        end
        if(~strcmp(prev2(1:16),prev(1:16)))
            fprintf(fo,"\n\n");
        end
        offsets(1)=0;
        offsets(2)=x2-x1;
        offsets(3)=x3-x1;
        offsets(4)=x4-x1;
        offsets(5)=x5-x1;

        %name
        for nnn=1:length(f)
if(f(nnn)==',')
    f(nnn)='!';
end
        end
        fprintf(fo,"%s,",f(1:size(f,2)-4));

        %r1start r1len
        fprintf(fo,"%i,%i,",x1,x2-x1);

        %r2start r2len
        fprintf(fo,"%i,%i,",x2,x3-x2);

        %r3start r3len
        fprintf(fo,"%i,%i,",x3,x4-x3);

        %r4start r4len
        fprintf(fo,"%i,%i,",x4,x5-x4);

        %r5start r5len
        fprintf(fo,"%i,%i,",x5,x6-x5);

        %r5end totallen
        fprintf(fo,"%i,%i,",x6,x6-x1);

        %r1dapi r1gfp r1both
        fprintf(fo,"%i,%i,%i,",counts(1,1),counts(1,2),counts(1,3));

        %r2dapi r2gfp r2both
        fprintf(fo,"%i,%i,%i,",counts(2,1),counts(2,2),counts(2,3));

        %r3dapi r3gfp r3both
        fprintf(fo,"%i,%i,%i,",counts(3,1),counts(3,2),counts(3,3));

        %r4dapi r4gfp r4both
        fprintf(fo,"%i,%i,%i,",counts(4,1),counts(4,2),counts(4,3));

        %r5dapi r5gfp r5both
        fprintf(fo,"%i,%i,%i,",counts(5,1),counts(5,2),counts(5,3));

        totcells=counts(1,1)+counts(2,1)+counts(3,1)+counts(4,1)+counts(5,1);
        totgfp=counts(1,2)+counts(2,2)+counts(3,2)+counts(4,2)+counts(5,2);
        totboth=counts(1,3)+counts(2,3)+counts(3,3)+counts(4,3)+counts(5,3);

        %totalcells totalgfp totalboth
        fprintf(fo,"%i,%i,%i,",totcells, totgfp, totboth);

        %name
        fprintf(fo,"%.6f,%.6f,%.6f,%s,", totcells/(x6-x1),totgfp/(x6-x1),totgfp/totcells,cm{1});

        %name
        obj=zeros(1,1);
        xposns=zeros(10000,3);
        percentiles=zeros(100,3);
        nob=0;
        nob2=0;
        for(ch=1:3)
            nob=0;
            nob2=0;
            for(reg=1:5)
                indices=allindices(reg,ch);
                currstats=allstats(reg,ch);
                currstats2=currstats{1};
                nobj=counts(reg,ch);
                for obj=1:nobj
                    list1=indices{1};
                    nob=nob+1;
                    objx=currstats2(list1(obj));
                    posn=round(objx.Centroid(1)+offsets(reg));
                    xposns(nob,ch)=posn;
                    ptile=round(posn/(x6-x1)*100);
                    ptile=max(ptile,1);
                    ptile=min(ptile,100);
                    percentiles(ptile,ch)=percentiles(ptile,ch)+1/(counts(1,ch)+counts(2,ch)+counts(3,ch)+counts(4,ch)+counts(5,ch));
                end %obj
            end %reg
        end %ch
        fprintf(fo,",");

        for ch=1:3
            for p=1:5
                dd=sum(percentiles(      ((p-1)*20+1):(p*20),   ch));
                fprintf(fo,"%.6f,",dd);
            end
            fprintf(fo,",");
        end

        for ch=1:3
            for p=1:10
                dd=sum(percentiles(       ((p-1)*10+1):(p*10)     ,ch));
                fprintf(fo,"%.6f,",dd);
            end
            fprintf(fo,",");
        end

        for ch=1:3
            for p=1:100
                fprintf(fo,"%.6f,",percentiles(p,ch));
            end
            fprintf(fo,",");
        end

            for p=1:5
                dd=sum(percentiles(  ((p-1)*20+1):(p*20),   2));
                dd2=sum(percentiles( ((p-1)*20+1):(p*20),   1));
                rr=dd/dd2;
                if isnan(rr) || isinf(rr)
                    fprintf(fo,",");
                else
                    fprintf(fo,"%.6f,",rr);
                end
            end
            fprintf(fo,",");

            for p=1:10
                dd=sum(percentiles(   ((p-1)*10+1):(p*10)     ,2));
                dd2=sum(percentiles(  ((p-1)*10+1):(p*10)     ,1));
                rr=dd/dd2;
                if isnan(rr) || isinf(rr)
                    fprintf(fo,",");
                else
                    fprintf(fo,"%.6f,",rr);
                end
            end
            fprintf(fo,",");

            for p=1:100
                dd=percentiles(p, 2);
                dd2=percentiles(p, 1);
                rr=dd/dd2;
                if isnan(rr) || isinf(rr)
                    fprintf(fo,",");
                else
                    fprintf(fo,"%.6f,",rr);
                end
            end
            fprintf(fo,",");


        fprintf(fo,"\n");

        for ch=1:3
            fprintf(fo2,"%s,All %s positions,",f,terms{ch});
            for p=1:(counts(1,ch)+counts(2,ch)+counts(3,ch)+counts(4,ch)+counts(5,ch))
                fprintf(fo2,"%i,",xposns(p,ch));
            end
            fprintf(fo2,"\n");
        end
        fprintf(fo2,"\n");
    end
end
fclose(fo);
fclose(fo2);
fclose all;
close all;

return

%show all images
for s=1:5
j=allstats{s,2};
for n=1:size(j,1)
    if(j(n).Area>150)
    p=j(n).PixelList;
    x1=min(p(:,1));
    y1=min(p(:,2));
    w=max(p(:,1))-x1;
    h=max(p(:,2))-y1;
    img=zeros(w,h);
    for t=1:size(p,1)
        img(p(t,1)-x1+1,p(t,2)-y1+1)=1;
    end
    imshow(img);
    end
end
end
```
