## Supplemental code 3 for "Gut hormone signaling drives sex differences in metabolism and behavior": cg2.m

```
function obj=cg2(datatable,rows,columns, type)
    d=zscore(datatable{rows,columns});

    % 
    % n=0;
    % f=0;
    % for t=rows
    %     n=n+1;
    %     if(strcmp(datatable.Name{t},'W1118') || strcmp(datatable.Gene{t},'W1118'))
    %         f=1;
    %         disp('asdfsa')
    %         break
    %     end
    % end
    % if(f>0)
    %     c=1;
    %     r=1;
    %     for t=columns
    %         r=1;
    %         for t2=1:size(rows,1)
    %             d(r,c)=d(r,c)-d(n,c);
    %             r=r+1;  
    %         end 
    %         c=c+1;
    %     end
    % end
    % 
    columnnames=datatable.Properties.VariableNames(columns);
    rownames=strcat(datatable.Gene(rows),'-',datatable.Sex(rows));
    obj=clustergram(d,'ColumnLabels',columnnames,'RowLabels',rownames, 'ColumnPDist', type, 'RowPDist', type, 'ColorMap','redbluecmap');
end
% euclidean jaccard minkowski correlation spearman
```
