## Supplemental code 3 for "Gut hormone signaling drives sex differences in metabolism and behavior": cg2noz.m

```
function cg2noz(datatable,rows,columns)
    d=datatable{rows,columns};
    columnnames=datatable.Properties.VariableNames(columns);
    rownames=strcat(datatable.Gene(rows),'-',datatable.Sex(rows));
    clustergram(d,'ColumnLabels',columnnames,'RowLabels',rownames);
end
```
