## Supplemental code 3 for "Gut hormone signaling drives sex differences in metabolism and behavior": cg.m

```
function cg(typex,sizz,sexx,rows,cols)
global datatables;
d=datatables{typex,sizz,sexx}();
columnnames=codatatables{typex,sizz,sexx}.Properties.VariableNames(firstc:lastc);
clustergram(d,'ColumnLabels',columnnamess{typex,sizz,sexx},'RowLabels',rownamess{typex,sizz,sexx});
end
```
