## Supplemental code 3 for "Gut hormone signaling drives sex differences in metabolism and behavior": circleintersection.m

```
function area=circleintersection(ax,ay,ar,bx,by,br)
dist=sqrt((bx-ax)^2+(by-ay)^2);
if  dist>= ar+br
    area=0;
    return
end
if ar>br && dist <= (ar-br)
    area=3.141592*br*br;
    return
end
if br>ar && dist <= (br-ar)
    area=3.141592*ar*ar;
    return
end

alpha = acos((ar*ar+dist*dist-br*br) / (2*ar*dist)) * 2;
beta = acos((br*br + dist*dist-ar*ar) / (2*br*dist)) * 2;
a1=0.5*beta*br*br-0.5*br*br*sin(beta);
a2=0.5*alpha*ar*ar-0.5*ar*ar*sin(alpha);
area=a1+a2;

end
```
