## Supplemental code 3 for "Gut hormone signaling drives sex differences in metabolism and behavior": findcorrelation2.m

```
function findcorrelation2(dataset, rows, columns, name, name2, listname, sexname)
d2=dataset{rows,columns};
d3=table(dataset.ScreenID, dataset.Name, dataset.Gene, dataset.Sex,'VariableNames', ["ScreenID",        "Name",        "Gene",         "Sex"]);
d3=d3(rows,:);
d2=zscore(d2);
% 
% n=-1;
% for t=1:max(size(rows,1),size(rows,2))
%     if(strcmp(d3.Name{t},'W1118') || strcmp(d3.Gene{t},'W1118') || strcmp(d3.Gene{t},'W1118-M')|| strcmp(d3.Name{t},'W1118-M')|| strcmp(d3.Gene{t},'W1118-F')|| strcmp(d3.Name{t},'W1118-F'))
%         n=t;
%         disp('1234213')
%         break
%     end
% end
% if(n>0)
%     c=1;
%     r=1;
%     for t=columns
%         r=1;
%         for t2=rows
%             d2(r,c)=d2(r,c)-d2(n,c)
%             r=r+1;
%         end
%         c=c+1;
%     end
% end

d2=d2';
out=fopen(strcat("Trees\",name," ", name2," correlations.txt"),'a');
fprintf(out,"%s\n%s\n",upper(listname),upper(sexname));
[x,y]=corr(d2,'Type','Spearman');
f=findgene(d3,name,'f')
m=findgene(d3,name,'m')
f2=findgene(d3,name2,'f')
m2=findgene(d3,name2,'m')

if(f~=-1 && f2~=-1)
r3=x(f,f2);
p3=y(f,f2);
fprintf(out,"%s-f vs %s-f r=%f p=%f\n",name,name2,r3,p3);
end

if(m~=-1 && m2~=-1)
r4=x(m,m2);
p4=y(m,m2);
fprintf(out,"%s-m vs %s-m r=%f p=%f\n",name,name2,r4,p4);
end

if(m~=-1 && f~=-1 && m2 ~=-1 && f2 ~=-1)
r=x(f,m);
p=y(f,m);
r2=x(f2,m2);
p2=y(f2,m2);
fprintf(out,"%s-%s vs %s-%s r=%f p=%f\n",name,'f',name,'m',r,p);
fprintf(out,"%s-%s vs %s-%s r=%f p=%f\n",name2,'f',name2,'m',r2,p2);
end


if f~=-1
    [list, ranks]=sort(abs(x(f,:)));
    for t=length(list):-1:length(list)-10
        fprintf(out,"Correlation %f between %s-F and %s-%s (p=%1.2e)\n",x(f,ranks(t)),name,d3.Gene{ranks(t)},d3.Sex{ranks(t)},y(f,ranks(t)));
    end
end

if(m~=-1)
    [list, ranks]=sort(abs(x(m,:)));
for t=length(list):-1:length(list)-10
fprintf(out,"Correlation %f between %s-M and %s-%s (p=%1.2e)\n",x(m,ranks(t)),name,d3.Gene{ranks(t)},d3.Sex{ranks(t)},y(m,ranks(t)));
end
end

if(f2~=-1)
    [list, ranks]=sort(abs(x(f2,:)));
for t=length(list):-1:length(list)-10
fprintf(out,"Correlation %f between %s-F and %s-%s (p=%1.2e)\n",x(f2,ranks(t)),name2,d3.Gene{ranks(t)},d3.Sex{ranks(t)},y(f2,ranks(t)));
end
end

if m2~=-1
[list, ranks]=sort(abs(x(m2,:)));
for t=length(list):-1:length(list)-10
fprintf(out,"Correlation %f between %s-M and %s-%s (p=%1.2e)\n",x(m2,ranks(t)),name2,d3.Gene{ranks(t)},d3.Sex{ranks(t)},y(m2,ranks(t)));
end
end
fclose(out);

end
```
