## Supplemental code 3 for "Gut hormone signaling drives sex differences in metabolism and behavior": findcorrelation3.m

```
function findcorrelation3(dataset, rows, columns, name, name2, listname, sexname)
d2=dataset{rows,columns};
d3=table(dataset.ScreenID, dataset.Name, dataset.Gene, dataset.Sex,'VariableNames', ["ScreenID",        "Name",        "Gene",         "Sex"]);
d3=d3(rows,:);
d2=zscore(d2);
% 
% n=-1;
% for t=1:max(size(rows,1),size(rows,2))
%     if(strcmp(d3.Name{t},'W1118') || strcmp(d3.Gene{t},'W1118') || strcmp(d3.Gene{t},'W1118-M')|| strcmp(d3.Name{t},'W1118-M')|| strcmp(d3.Gene{t},'W1118-F')|| strcmp(d3.Name{t},'W1118-F'))
%         n=t;
%         disp('1234213')
%         break
%     end
% end
% if(n>0)
%     c=1;
%     r=1;
%     for t=columns
%         r=1;
%         for t2=rows
%             d2(r,c)=d2(r,c)-d2(n,c)
%             r=r+1;
%         end
%         c=c+1;
%     end
% end

d2=d2';
out=fopen(strcat("Trees\all correlations.txt"),'a');
% fprintf(out,"%s\n%s\n",upper(listname),upper(sexname));
[x,y]=corr(d2,'Type','Spearman');
f=findgene(d3,name,'f')
m=findgene(d3,name,'m')
% f2=findgene(d3,name2,'f')
% m2=findgene(d3,name2,'m')


if f~=-1
    [list, ranks]=sort(abs(x(f,:)));
    for t=1:length(list)
        fprintf(out,"%s,%s,%f,%s-F,%s-%s,%1.2e\n",listname, sexname, x(f,ranks(t)),name,d3.Gene{ranks(t)},d3.Sex{ranks(t)},y(f,ranks(t)));
    end
end

if(m~=-1)
    [list, ranks]=sort(abs(x(m,:)));
    for t=1:length(list)
        fprintf(out,"%s,%s,%f,%s-M,%s-%s,%1.2e\n",listname, sexname, x(m,ranks(t)),name,d3.Gene{ranks(t)},d3.Sex{ranks(t)},y(m,ranks(t)));
    end
end

% if(f2~=-1)
%     [list, ranks]=sort(abs(x(f2,:)));
% for t=length(list):-1:length(list)-10
% fprintf(out,"%s,%s,%f,%s-F,%s-%s,%1.2e\n",listname, sexname, x(f2,ranks(t)),name2,d3.Gene{ranks(t)},d3.Sex{ranks(t)},y(f2,ranks(t)));
% end
% end
% 
% if m2~=-1
% [list, ranks]=sort(abs(x(m2,:)));
% for t=length(list):-1:length(list)-10
% fprintf(out,"%s,%s,%f,%s-M,%s-%s,%1.2e\n",listname, sexname, x(m2,ranks(t)),name2,d3.Gene{ranks(t)},d3.Sex{ranks(t)},y(m2,ranks(t)));
% end
% end
fclose(out);

end
```
