## Supplemental code 3 for "Gut hormone signaling drives sex differences in metabolism and behavior": findcorrelationMFs.m

```
function findcorrelationMFs(dataset, name)
d2=zscore(dataset{:,5:22});
d2=d2';
[x,y]=corr(d2,'Type','Spearman');
r=zeros(166,1);
p=zeros(166,1);
for t=1:166
    r(t)=x(t*2-1,2*t);
    p(t)=y(t*2-1,2*t);
end

[r2,ranks]=sort(r);
p2=p(ranks);
for t=1:166
fprintf('Sorted by correlation: Male-female correlation %f (p=%fe) for %s\n',r2(t),p2(t),dataset.Gene{ranks(t)*2});
end


[p2,ranks]=sort(p);
r2=r(ranks);
for t=1:166
fprintf('Sorted by p value: Male-female correlation %f (p=%f) for %s\n',r2(t),p2(t),dataset.Gene{ranks(t)*2});
end


end
```
