## Supplemental code 3 for "Gut hormone signaling drives sex differences in metabolism and behavior": findgene2.m

```
function x=findgene2(dataset,name)
target=upper(name);
ll=size(dataset,1);
for x=1:ll
    if strcmp(upper(strcat(dataset.Gene{x},dataset.Sex{x})),target) || strcmp(upper(strcat(dataset.Name{x},dataset.Sex{x})),target)
        return
    end
end
x=-1;
return 
end
```
