## Supplemental code 3 for "Gut hormone signaling drives sex differences in metabolism and behavior": findintree.m

```
function findintree(tree,gene)
rlc.Labels=tree.RowLabels;
colors=cell(1,1);

for t=1:length(tree.RowLabels)
colors{t}='#ffaaaa';
x=tree.RowLabels{t};
if(x(length(x))=='M')
    colors{t}='#aaaaff';
end
if strcmp(upper(tree.RowLabels{t}),upper(strcat(gene,'-F')))
    colors{t}='red';
end
if strcmp(upper(tree.RowLabels{t}),upper(strcat(gene,'-M')))
    colors{t}='blue';
end
end
rlc.Colors=colors';
tree.RowLabelsColor=rlc;
tree.LabelsWithMarkers=true;
end
```
