## Supplemental code 3 for "Gut hormone signaling drives sex differences in metabolism and behavior": MainAnalysisScript.m

```
global malelm;
global femalelm;
global malex
global femalex
global malefit
global femalefit
global femalefitS
global malefitS
global maley
global femaley


for t=5 %5:24
%    for v=(t+1):24
    for v=5:24
        titletxt=strcat("Abs ",datasetabsolutesmallboth.Properties.VariableNames{t}," x ",datasetabsolutesmallboth.Properties.VariableNames{v});
        fprintf("%f %f %s\n",t,v,titletxt);
        %make2dplotnozeros([datasetabsolutesmallboth{:,t}],   [datasetabsolutesmallboth{:,v}],     datasetabsolutesmallboth.Gene,   datasetabsolutesmallboth.Sex, titletxt, datasetabsolutesmallboth.Properties.VariableNames{t}, datasetabsolutesmallboth.Properties.VariableNames{v});
        titletxt=strcat("Abs ",datasetabsolutesmallboth.Properties.VariableNames{t}," x ",datasetabsolutesmallboth.Properties.VariableNames{v},", Cleaner");
        make2dplotnozeros5([datasetabsolutesmallboth{:,t}],   [datasetabsolutesmallboth{:,v}],     datasetabsolutesmallboth.Gene,   datasetabsolutesmallboth.Sex, titletxt, datasetabsolutesmallboth.Properties.VariableNames{t}, datasetabsolutesmallboth.Properties.VariableNames{v});
        close all
    end
end


for t=5   %5:24
%    for v=(t+1):24
    for v=5:24
        titletxt=strcat("Delta ",datasetdeltasmallboth.Properties.VariableNames{t}," x ",datasetdeltasmallboth.Properties.VariableNames{v});
        fprintf("%f %f %s\n",t,v,titletxt);
        %make2dplotnozeros([datasetdeltasmallboth{:,t}],   [datasetdeltasmallboth{:,v}],     datasetdeltasmallboth.Gene,   datasetdeltasmallboth.Sex, titletxt, datasetdeltasmallboth.Properties.VariableNames{t},datasetdeltasmallboth.Properties.VariableNames{v}  );
        titletxt=strcat("Delta ",datasetdeltasmallboth.Properties.VariableNames{t}," x ",datasetdeltasmallboth.Properties.VariableNames{v},", Cleaner");
        make2dplotnozeros5([datasetdeltasmallboth{:,t}],   [datasetdeltasmallboth{:,v}],     datasetdeltasmallboth.Gene,   datasetdeltasmallboth.Sex, titletxt, datasetdeltasmallboth.Properties.VariableNames{t},datasetdeltasmallboth.Properties.VariableNames{v}  );
        close all
    end
end

return


%
%load("datasets.mat");

% 
% 
% datasetzsmallboth=datasetdeltasmallboth;
%  %return
% 
% 
% 
% zlim=1.5;
% for t=5:22
%     msd=std(datasetzsmallboth{2:2:332,t});
%     fsd=std(datasetzsmallboth{1:2:332,t});
%     mlist=cell(10,2);
%     flist=cell(10,2);
%     mlen=0;
%     flen=0;
%     for n=1:166
%         datasetzsmallboth{2*n,t}=datasetzsmallboth{2*n,t}/msd;
%         datasetzsmallboth{2*n-1,t}=datasetzsmallboth{2*n-1,t}/fsd;
%         if(abs(datasetzsmallboth{2*n,t})>=zlim)
%             mlen=mlen+1;
%             mlist{mlen,1}=datasetzsmallboth{2*n,3};
%             mlist{mlen,2}=datasetzsmallboth{2*n,t};
% %            fprintf("%s: %s %f\n",datasetzsmallboth.Properties.VariableNames{t},mlist{mlen,1}{1},mlist{mlen,2});
%         end
%         if(abs(datasetzsmallboth{2*n-1,t})>=zlim)
%             flen=flen+1;
%             flist{flen,1}=datasetzsmallboth{2*n-1,3};
%             flist{flen,2}=datasetzsmallboth{2*n-1,t};
% %            fprintf("%s: %s %f\n",datasetzsmallboth.Properties.VariableNames{t},flist{flen,1}{1},flist{flen,2});
%         end
%     end
%     mlen2=mlen;
%     flen2=flen;
%     bothlen=0;
%     bothlist=cell(10,3);
%     matched=0;
%     for n=1:mlen
%         for o=1:flen
%             if(strcmp(mlist{n,1}{1},flist{o,1}{1}))
%                 mlen2=mlen2-1;
%                 flen2=flen2-1;
%                 bothlen=bothlen+1;
%                 bothlist{bothlen,1}=mlist{n,1};
%                 bothlist{bothlen,2}=flist{o,2};
%                 bothlist{bothlen,3}=mlist{n,2};
%                 mlist{n,1}={"both"};
%                 flist{o,1}={"both"};
%                 fprintf("Both %s\n",mlist{n,1}{1});
%                 matched=1;
%                 n=1000;
%                 o=1000;
%                 break
%             end
%         end
%     end
%     d=venn2(mlen,bothlen,flen);
%     if( d==-1 )
%         fprintf("No venn2 fit\n");
%         th=-10;
%         return;
%     end
%     ff=gcf;
%     ff.Units='normalized';
%     ff.Position=[0 0 1 1];
%     drawnow
%     ra=sqrt(mlen/3.141592);
%     circle(0,0,ra);
%     rb=sqrt(flen/3.141592);
%     circle(d,0,rb);
%     ff.CurrentAxes.YLim=[-6 6];
%     axis equal
%     drawnow
%     mtxt=sprintf("Males (%i)\n",mlen2);
%     ftxt=sprintf("Females (%i)\n",flen2);
%     bothtxt=sprintf("Both (%i)\n",bothlen);
%     includez=false;
%     single=true;
% 
% for rep=1:3
%     switch rep
%         case 1
%             txt=mtxt;
%             len1=mlen;
%             len2=mlen2;
%             xlist=mlist;
%         case 2
%             txt=ftxt;
%             len2=flen2;
%             len1=flen;
%             xlist=flist;
%         case 3
%             txt=bothtxt;
%             len2=bothlen;
%             len1=bothlen;
%             xlist=bothlist;
%         end
%     mm=0;
%     secreted=0;
%     receptors=0;
%     transporters=0;
%     enzymes=0;
%     peptides=0;
%     reqamon=0;
%     reqphm=0;
%     reqsvr=0;
%     split=1;
%     if(len2>10)
%         split=3;
%     end
%     if(len2>20)
%         split=4;
%     end
%     if(len2>30)
%         split=5;
%     end
%     if(len2>40)
%         split=6;
%     end
%     if(len2>60)
%         split=7;
%     end
%     for m=1:len1
%         if(~strcmp("both",xlist{m,1}{1}))
%             mm=mm+1;
%             txt=sprintf("%s%s",txt,xlist{m,1}{1});
%             if(includez)
%                 if(rep~=3)
%                     txt=sprintf("%s (Z=%f)",txt,xlist{m,2});
%                 else
%                     txt=sprintf("%s (Zf=%f, Zm=%f)",txt,xlist{m,2},xlist{m,3});
%                 end
%             else
%                 if(rep==3 && xlist{m,2}*xlist{m,3}<0)
%                     txt=sprintf("%s*",txt);
%                 end
%             end
%             if(mod(mm,split)==0 || single)
%                 txt=sprintf("%s\n",txt);
%             else
%                 txt=sprintf("%s    ",txt);
%             end
%             for c=1:2:332
%                 if(strcmp(xlist{m,1}{1},datasetabsolutefullboth.Gene(c)))
%                     if strcmp(datasetabsolutefullboth.isSecreted(c),"TRUE")
%                         secreted=secreted+1;
%                     end
%                     if strcmp(datasetabsolutefullboth.requiresPHM(c),"TRUE")
%                         reqphm=reqphm+1;
%                     end
%                     if strcmp(datasetabsolutefullboth.requiresSilver(c),"TRUE")
%                         reqsvr=reqsvr+1;
%                     end
%                     if strcmp(datasetabsolutefullboth.requiresAmon(c),"TRUE")
%                         reqamon=reqamon+1;
%                     end
%                     if strcmp(datasetabsolutefullboth.isNPeptide(c),"TRUE")
%                         peptides=peptides+1;
%                     end
%                     if strcmp(datasetabsolutefullboth.isEnzyme(c),"TRUE")
%                         enzymes=enzymes+1;
%                     end
%                     if strcmp(datasetabsolutefullboth.isReceptor(c),"TRUE")
%                         receptors=receptors+1;
%                     end
%                     if strcmp(datasetabsolutefullboth.isTransporter(c),"TRUE")
%                         transporters=transporters+1;
%                     end
%                     break
%                 end
%             end
%         end
%     end
%     txt=sprintf("%s\n\nSecretedThings: %i    Enzymes: %i\nReceptors: %i    Transporters: %i\nPeptides: %i (Amon: %i; Svr: %i; PHM: %i)",txt,secreted,enzymes,receptors,transporters,peptides,reqamon,reqsvr,reqphm);
%     switch rep
%         case 1
%             mtxt=txt;
%         case 2
%             ftxt=txt;
%         case 3
%             bothtxt=txt;
%         end
% end
% 
% 
% 
%     midpt=(ra+d-rb)/2;
%     text(    0, 0,mtxt,'HorizontalAlignment','Center','Color','blue');
%     text(    d, 0, ftxt,'HorizontalAlignment','Center','Color','red');
%     text(midpt, 0, bothtxt,'HorizontalAlignment','Center','Color','#ff00ff');
%     text(midpt,1,datasetzsmallboth.Properties.VariableNames{t},'HorizontalAlignment','Center','Color','black');
%     %text(midpt,max(ra,rb)+1,datasetzsmallboth.Properties.VariableNames{t},'HorizontalAlignment','Center');
%     %ff.Children.YLim=[ff.Children.YLim(1)-1 ff.Children.YLim(2)];
%     drawnow
%         saveas(gcf,strcat("PNGs\Venn15",datasetzsmallboth.Properties.VariableNames{t},".png"));
%                 print(gcf,'-vector','-dpdf',strcat("PDFs\Venn15",datasetzsmallboth.Properties.VariableNames{t},".pdf"));
% close all
% end
% return
% % 
% % 
% % %dataset=zscore(datasetdeltasmallboth{1:332,5:22});
% % %for n=1:8
% % %k=kmeans(dataset,n);
% % %for t=1:332
% % %    names{t}=strcat(datasetdeltasmallboth.Gene{t},"-",datasetdeltasmallboth.Sex{t});
% % %end
% % %make2dplotnozeros2(dataset(:,1),dataset(:,2), names, k, strcat("Delta ",num2str(n),"-Means Clustering not PCA"), "1111", "2222");
% % %make2dplotnozeros3(dataset(:,1),dataset(:,2), names, k, strcat("Delta ",num2str(n),"-Means Clustering not PCA, Cleaner"), "1111", "2222");
% % %end%
% % %
% % %return
% % %%%%%%%%%%%%%%%%%%%%%%%%%%%%%%%%%%%%%%%%%%%%


global datasets;
global columnnamess;
global rownamess;
global specialgenes;
global fname;
global suffix;
global savepdfs;
global savepngs;
savepdfs=true;
savepngs=true;
warning off

makeallcombos=true;

makeplots=savepdfs || savepngs;

specialgenes=cell(4,3);
specialgenes{1,1}="W1118";     specialgenes{1,2}="#00ff00";  specialgenes{1,3}="#00aa00";
specialgenes{2,1}="Phm";       specialgenes{2,2}="#dddd00";  specialgenes{2,3}="#aaaa00";
specialgenes{3,1}="amon";      specialgenes{3,2}="#dddd00";  specialgenes{3,3}="#aaaa00";
specialgenes{4,1}="Pal2";      specialgenes{4,2}="#dddd00";  specialgenes{4,3}="#aaaa00";
specialgenes{5,1}="svr";       specialgenes{5,2}="#dddd00";  specialgenes{5,3}="#aaaa00";
specialgenes{6,1}="AstC";      specialgenes{6,2}="#88ff00";  specialgenes{6,3}="#66aa00";
specialgenes{7,1}="Dh31";      specialgenes{7,2}="#00ffff";  specialgenes{7,3}="#00aaaa";
specialgenes{8,1}="Orcokinin"; specialgenes{8,2}="#ff00ff";  specialgenes{8,3}="#aa00aa";
specialgenes{9,1}="AstA";      specialgenes{9,2}="#00ff88";  specialgenes{9,3}="#00aa66";
specialgenes{10,1}="NPF";      specialgenes{10,2}="#ff8800"; specialgenes{10,3}="#aa6600";
specialgenes{11,1}="Tk";       specialgenes{11,2}="#0088ff"; specialgenes{11,3}="#0066aa";
specialgenes{12,1}="7B2";      specialgenes{12,2}="#dddd00"; specialgenes{12,3}="#aaaa00";
absolute=1;
delta=2;
full=1;
small=2;
both=1;
male=2;
female=3;
datasets=cell(2,2,3);
rownamess=cell(2,2,3);
columnnamess=cell(2,2,3);
explaineds=cell(2,2,3);
coeffs=cell(2,2,3);
scoress=cell(2,2,3);
fmdists=cell(2,2,3);
fmdists2=cell(2,2,3);
hdists=cell(2,2,3);
hdists2=cell(2,2,3);
md2fws=cell(2,2,3);
fd2mws=cell(2,2,3);
md2mws=cell(2,2,3);
fd2fws=cell(2,2,3);
angless=cell(2,2,3);

dataabs=readtable(strcat("CombinedData.csv"));
datadelta=dataabs;
for t=5:81
    for v=1:2:331
        val=(dataabs{v,t}-dataabs{331,t})/dataabs{331,t};
        if(isnan(val))
            val=0;
        end
        datadelta{v,t}=val;

        val=(dataabs{v+1,t}-dataabs{332,t})/dataabs{332,t};
        if(isnan(val))
            val=0;
        end
        datadelta{v+1,t}=val;
    end
end


for flx=1:2
    switch flx
        case 1
            fname="Abs";
            dataorig=dataabs;
        case 2
            fname="Delta";
            dataorig=datadelta;
    end
    for szx=1:2
        switch szx
            case 1
                datasize="Full";
            case 2
                datasize="Small";
        end
        for sxx=1:3
            switch sxx
                case 1
                    sexes="Both";
                case 2
                    sexes="Male";
                case 3
                    sexes="Female";
            end
            fprintf("%s %s %s\n",fname,datasize,sexes);
            suffix=strcat("-",datasize,"-",sexes);
            %dataorig=readtable(strcat(fname,".csv"));
            datasmall=table(dataorig.ScreenID, dataorig.Name, dataorig.Gene, dataorig.Sex, dataorig.Starvation, dataorig.Ionic, dataorig.Oxidative, dataorig.TAGf, dataorig.TAGs, dataorig.TAGc, dataorig.GlucoseF, dataorig.GlucoseS, dataorig.GlucoseC, dataorig.GlycogenF, dataorig.GlycogenS, dataorig.GlycogenC, dataorig.FeedingDurationAll, dataorig.NumEventsAll, dataorig.DaySleepF, dataorig.DaySleepS, dataorig.NightSleepF, dataorig.NightSleepS, ...
dataorig.FlyAtlas2Male,	dataorig.FlyAtlas2Female,	dataorig.FlyAtlas2Midgut,	dataorig.DimorphismIndex,	dataorig.DimorphismIndex2, dataorig.AstAEE, dataorig.AstCEE, dataorig.NPFEE, ...
'VariableNames',               ["ScreenID",        "Name",        "Gene",         "Sex",      "Starvation",        "Ionic",       "Oxidative",           "TAGf",     "TAGs",          "TAGc",         "GlucF",           "GlucS",          "GlucC",           "GlycogenF",         "GlycogenS",        "GlycogenC",       "FeedingDurationAll",       "NumEventsAll",             "DaySleepF",        "DaySleepS",        "NightSleepF",         "NightSleepS", ...       
         "FlyAtlas2Male",	"FlyAtlas2Female",     	"FlyAtlas2Midgut",	           "DimorphismIndex",	              "DimorphismIndex2", "AstAEE", "AstCEE", "NPFEE"]);
            origrows=size(dataorig,1);

            switch(suffix)
                case "-Full-Both"
                    dataset=dataorig;
                    if(flx==absolute)
                        datasetabsolutefullboth=dataset;
                    else
                        datasetdeltafullboth=dataset;
                    end
                    columns=size(dataset,2);
                    firstcolumn=6;
                    lastcolumn=81;
                    step=2;
                case "-Full-Male"
                    dataset=dataorig(2:2:origrows,:);
                    if(flx==absolute)
                        datasetabsolutefullmale=dataset;
                    else
                        datasetdeltafullmale=dataset;
                    end
                    columns=size(dataset,2);
                    firstcolumn=6;
                    lastcolumn=81;
                    step=1;
                case "-Full-Female"
                    dataset=dataorig(1:2:origrows,:);
                    if(flx==absolute)
                        datasetabsolutefullfemale=dataset;
                    else
                        datasetdeltafullfemale=dataset;
                    end
                    columns=size(dataset,2);
                    firstcolumn=5;
                    lastcolumn=81;
                    step=1;
                case "-Small-Both"
                    dataset=datasmall;
                    if(flx==absolute)
                        datasetabsolutesmallboth=dataset;
                    else
                        datasetdeltasmallboth=dataset;
                    end
                    firstcolumn=5;
                    columns=size(dataset,2);
                    lastcolumn=22;
                    step=2;
                case "-Small-Male"
                    dataset=datasmall(2:2:origrows,:);
                    if(flx==absolute)
                        datasetabsolutesmallmale=dataset;
                    else
                        datasetdeltasmallmale=dataset;
                    end
                    firstcolumn=5;
                    columns=size(dataset,2);
                    lastcolumn=22;
                    step=1;
                case "-Small-Female"
                    dataset=datasmall(1:2:origrows,:);
                    if(flx==absolute)
                        datasetabsolutesmallfemale=dataset;
                    else
                        datasetdeltasmallfemale=dataset;
                    end

                    firstcolumn=5;
                    columns=size(dataset,2);
                    lastcolumn=22;
                    step=1;
            end

            rows=size(dataset,1);
            genotypes=rows/step;

            standardizeddata=zscore(dataset{1:rows,firstcolumn:lastcolumn});
            %standardizeddata=dataset{1:rows,firstcolumn:lastcolumn};
            datasets{flx,szx,sxx}=standardizeddata;
            columnnames=dataset.Properties.VariableNames(firstcolumn:lastcolumn);
            columnnamess{flx,szx,sxx}=columnnames;

            %rownamess{flx,szx,sxx}=rownames;

            [coeffs{flx,szx,sxx},scores{flx,szx,sxx},latent,tsquared,explaineds{flx,szx,sxx},mu] = pca(standardizeddata);
            score=scores{flx,szx,sxx};
            pairnames=dataset.Gene(1:2:rows);
            scale=120;
            %PCA chart
            for fff=1:2
if(makeplots)
            figure('visible','off')
            hold on
end
            for t=1:step:(genotypes*step)
                c1="red";
                c2="blue";
                if(sexes=="Male")
                    c1="blue";
                end
                c5=c1;
                s=10;
                m='.';
                lw=0.1;
                lw2=0.1;
                tc='#888888';
                fw='normal';
                for nn=1:size(specialgenes,1)
                    if dataset.Gene(t)==specialgenes{nn,1}
                        c1=specialgenes{nn,2};
                        c2=specialgenes{nn,3};
                        s=15;
                        lw=2;
                        lw2=0.5;
                        m='o';
                        c5='#000000';
                        tc='black';
                        fw='bold';
                        break
                    end
                end
                if dataset.Gene{t}=="W1118"
                    if(sexes=="Both")
                        wxf=score(t,1);
                        wyf=score(t,2);
                        wxm=score(t+1,1);
                        wym=score(t+1,2);
                    end
                    if(sexes=="Female")
                        wxf=score(t,1);
                        wyf=score(t,2);
                    end
                    if(sexes=="Male")
                        wxm=score(t,1);
                        wym=score(t,2);
                    end
                end
                x1=score(t,1);
                y1=score(t,2);
                if(sexes=="Both")
                    x2=score(t+1,1);
                    y2=score(t+1,2);
if(makeplots)
                    plot([x1 x2], [y1 y2],'Color','#aaaaaa','LineWidth',lw);
                    plot(x2, y2, 'Marker','o','MarkerSize',s,'MarkerFaceColor',c2,'MarkerEdgeColor','black','LineWidth',lw2);
                    %text(x2+s/scale, y2+s/scale, dataset.Gene{t},"FontSize", 8,'Rotation',45,'Color',tc,'FontWeight',fw);
                    if(fff==1)
                        text(x2, y2, dataset.Gene{t},"FontSize", 8,'Rotation',45,'Color',tc,'FontWeight',fw,'HorizontalAlignment','center');
                    end
end
                end
                if(makeplots)
                plot(x1, y1, 'Marker','o','MarkerSize',s,'MarkerFaceColor',c1,'MarkerEdgeColor','black','LineWidth',lw2);
%                text(x1+s/scale, y1+s/scale, dataset.Gene{t},"FontSize", 8,'Rotation',45,'Color',tc,'FontWeight',fw);
                if(fff==1)
                    text(x1, y1, dataset.Gene{t},"FontSize", 8,'Rotation',45,'Color',tc,'FontWeight',fw,'HorizontalAlignment','center');
                end
                end
            end %first round

            %put thick lines on top
            for t=1:step:(genotypes*step)
                s=15;
                m='.';
                lw=0.1;
                tc='black';
                for nn=1:size(specialgenes,1)
                    if dataset.Gene(t)==specialgenes{nn,1}
                        c1=specialgenes{nn,2};
                        c2=specialgenes{nn,3};
                        s=15;
                        lw=2;
                        lw2=0.5;
                        x1=score(t,1);
                        y1=score(t,2);
                        if(sexes=="Both")
                            x2=score(t+1,1);
                            y2=score(t+1,2);
%                            plot([x1 x2], [y1 y2],'Color',specialgenes{nn,2},'LineWidth',lw);
if(makeplots)
    plot([x1 x2], [y1 y2],'Color','black','LineWidth',lw);
end
%                            plot(x2, y2, 'Marker','o','MarkerSize',s,'MarkerFaceColor',c2, 'MarkerEdgeColor','black','LineWidth',lw2);
%                            text(x2+s/scale, y2+s/scale, dataset.Gene{t},"FontSize", 8,'Rotation',45,'Color',tc,'FontWeight','bold');
                        end
%                        plot(x1, y1, 'Marker','o','MarkerSize',s,'MarkerFaceColor',c1,'MarkerEdgeColor','black','LineWidth',lw2);
%                        text(x1+s/scale, y1+s/scale, dataset.Gene{t},"FontSize", 8,'Rotation',45,'Color',tc,'FontWeight','bold');
                        break
                    end
                end
            end

            %put special dots and names on top
            for t=1:step:(genotypes*step)
                s=15;
                m='.';
                lw=0.1;
                tc='black';
                for nn=1:size(specialgenes,1)
                    if dataset.Gene(t)==specialgenes{nn,1}
                        c1=specialgenes{nn,2};
                        c2=specialgenes{nn,3};
                        s=15;
                        lw=2;
                        lw2=0.5;
                        x1=score(t,1);
                        y1=score(t,2);
                        if(sexes=="Both")
                            x2=score(t+1,1);
                            y2=score(t+1,2);
%                            plot([x1 x2], [y1 y2],'Color',specialgenes{nn,2},'LineWidth',lw);
%                            plot([x1 x2], [y1 y2],'Color','black','LineWidth',lw);
                            if(makeplots)
                                plot(x2, y2, 'Marker','o','MarkerSize',s,'MarkerFaceColor',c2, 'MarkerEdgeColor','black','LineWidth',lw2);
%                            text(x2+s/scale, y2+s/scale, dataset.Gene{t},"FontSize", 8,'Rotation',45,'Color',tc,'FontWeight','bold');
                            text(x2, y2, dataset.Gene{t},"FontSize", 8,'Rotation',45,'Color',tc,'FontWeight','bold','HorizontalAlignment','center');
                            end

                        end
                        if(makeplots)
                        plot(x1, y1, 'Marker','o','MarkerSize',s,'MarkerFaceColor',c1,'MarkerEdgeColor','black','LineWidth',lw2);
%                        text(x1+s/scale, y1+s/scale, dataset.Gene{t},"FontSize", 8,'Rotation',45,'Color',tc,'FontWeight','bold');
                        text(x1, y1, dataset.Gene{t},"FontSize", 8,'Rotation',45,'Color',tc,'FontWeight','bold','HorizontalAlignment','center');
                        end
                        break
                    end
                end
            end
            if(makeplots)
            ff=gcf;
            ff.Units='normalized';
            ff.Position=[0 0 1 1];
            tx=strcat(fname," ",suffix," MainPCA Cleaner");
            if(fff==1)
                tx=strcat(fname," ",suffix," MainPCA");
            end
            title(tx);
            saveas(gcf,strcat("PNGs\",tx,".png"));
            if(savepdfs)
                print(gcf,'-vector','-dpdf',strcat("PDFs\",tx,".pdf"));
            end
            end
            close all
            end


            if(sexes=="Both")
                angles=zeros(genotypes,1);
                fdisttomw=zeros(genotypes,1);
                fdisttofw=zeros(genotypes,1);
                mdisttofw=zeros(genotypes,1);
                mdisttomw=zeros(genotypes,1);
                for t=1:2:(rows-1)
                    fx=score(t,1);
                    mx=score(t+1,1);
                    fy=score(t,2);
                    my=score(t+1,2);
                    mdisttofw((t+1)/2)=sqrt((mx-wxf)^2+(my-wyf)^2);
                    fdisttomw((t+1)/2)=sqrt((fx-wxm)^2+(fy-wym)^2);
                    mdisttomw((t+1)/2)=sqrt((mx-wxm)^2+(my-wym)^2);
                    fdisttofw((t+1)/2)=sqrt((fx-wxf)^2+(fy-wyf)^2);
                    an=atan((my-fy)/(mx-fx))/3.14159*180;
                    if(isnan(an))
                        an=0;
                    end
                    angles((t+1)/2)=abs(an);
                end
                fd2mws{flx,szx,sxx}=fdisttomw;
                md2fws{flx,szx,sxx}=mdisttofw;
                fd2fws{flx,szx,sxx}=fdisttofw;
                md2mws{flx,szx,sxx}=mdisttomw;
                angless{flx,szx,sxx}=angles;
if(makeplots)
    makesortedplot(fdisttomw,pairnames,strcat(fname," ", suffix," Female Distance to Male Control Sorted"));
                makesortedplot(mdisttofw,pairnames,strcat(fname," ", suffix," Male Distance to Female Control Sorted"));

                makesortedplot(fdisttofw,pairnames,strcat(fname," ", suffix," Female Distance to Female Control Sorted"));
                makesortedplot(mdisttomw,pairnames,strcat(fname," ", suffix," Male Distance to Male Control Sorted"));
                makesortedplot(angles,pairnames,strcat(fname," ", suffix," Female-to-Male Angles Sorted"));
end
                fmdist=zeros(genotypes,1);
                for t=1:2:(rows-1)
                    fx=score(t,1);
                    mx=score(t+1,1);
                    fy=score(t,2);
                    my=score(t+1,2);
                    fmdist((t+1)/2)=sqrt((mx-fx)^2+(my-fy)^2);
                    hyperfmdist=0;
                    for j=1:size(score,2)
                        hyperfmdist=hyperfmdist+(score(t,j)-score(t+1,j))^2;
                    end
                    hdist((t+1)/2)=sqrt(hyperfmdist);
                end
                fmdists{flx,szx,sxx}=fmdist;
                hdists{flx,szx,sxx}=hdist;
   if(makeplots)
       makesortedplot(fmdist,pairnames,strcat(fname," ", suffix," Female to Male Distance Sorted"));
                makesortedplot(hdist,pairnames,strcat(fname," ", suffix," Female to Male HyperDistance Sorted"));
   end
            end
        end
    end
end

for t=1:genotypes
    datasetabsolutesmallboth.FMDist(2*t-1)=fmdists{absolute,small,both}(t);
    datasetabsolutesmallboth.FMDist(2*t)=fmdists{absolute,small,both}(t);
    
    datasetabsolutesmallboth.FDist2Mw(2*t-1)=fd2mws{absolute,small,both}(t);
    datasetabsolutesmallboth.FDist2Mw(2*t)=fd2mws{absolute,small,both}(t);
    datasetabsolutesmallboth.FDist2Fw(2*t-1)=fd2fws{absolute,small,both}(t);
    datasetabsolutesmallboth.FDist2Fw(2*t)=fd2fws{absolute,small,both}(t);

    datasetabsolutesmallboth.MDist2Fw(2*t-1)=md2fws{absolute,small,both}(t);
    datasetabsolutesmallboth.MDist2Fw(2*t)=md2fws{absolute,small,both}(t);
    datasetabsolutesmallboth.MDist2Mw(2*t-1)=md2mws{absolute,small,both}(t);
    datasetabsolutesmallboth.MDist2Mw(2*t)=md2mws{absolute,small,both}(t);

    datasetabsolutesmallboth.Angles(2*t-1)=angless{absolute,small,both}(t);
    datasetabsolutesmallboth.Angles(2*t)=angless{absolute,small,both}(t);

    datasetabsolutesmallboth.HDist(2*t-1)=hdists{absolute,small,both}(t);
    datasetabsolutesmallboth.HDist(2*t)=hdists{absolute,small,both}(t);
end
    
for t=1:genotypes
    datasetdeltasmallboth.FMDist(2*t-1)=fmdists{delta,small,both}(t);
    datasetdeltasmallboth.FMDist(2*t)=fmdists{delta,small,both}(t);
    
    datasetdeltasmallboth.FDist2Mw(2*t-1)=fd2mws{delta,small,both}(t);
    datasetdeltasmallboth.FDist2Mw(2*t)=fd2mws{delta,small,both}(t);
    datasetdeltasmallboth.FDist2Fw(2*t-1)=fd2fws{delta,small,both}(t);
    datasetdeltasmallboth.FDist2Fw(2*t)=fd2fws{delta,small,both}(t);

    datasetdeltasmallboth.MDist2Mw(2*t-1)=md2mws{delta,small,both}(t);
    datasetdeltasmallboth.MDist2Mw(2*t)=md2mws{delta,small,both}(t);
    datasetdeltasmallboth.MDist2Fw(2*t-1)=md2fws{delta,small,both}(t);
    datasetdeltasmallboth.MDist2Fw(2*t)=md2fws{delta,small,both}(t);
    
    datasetdeltasmallboth.Angles(2*t-1)=angless{delta,small,both}(t);
    datasetdeltasmallboth.Angles(2*t)=angless{delta,small,both}(t);

    datasetdeltasmallboth.HDist(2*t-1)=hdists{delta,small,both}(t);
    datasetdeltasmallboth.HDist(2*t)=hdists{delta,small,both}(t);
end


phenotypedimorphismindex=zeros(166,1);
pairnames=dataorig.Gene(1:2:332);
for t=1:genotypes
    idx=(fd2mws{delta,small,both}(t)-md2fws{delta,small,both}(t))/(fd2mws{delta,small,both}(t)+md2fws{delta,small,both}(t));
    if(isnan(idx))
        idx=0;
    end
    phenotypedimorphismindex(t)=idx;
    datasetdeltasmallboth.PhenotypeDimorphismIndex(2*t-1)=idx;
    datasetdeltasmallboth.PhenotypeDimorphismIndex(2*t)=idx;
end
if(makeplots)
makesortedplot2(phenotypedimorphismindex,pairnames,"Delta PhenotypeDimorphismIndex -- PCA, Sorted");
makesortedplot(datasetdeltasmallboth.FMDist(1:2:332),pairnames,"Delta PhenotypeDisparity -- PCA, Sorted");
makesortedplot2(datasetdeltasmallboth.DimorphismIndex2(1:2:332),datasetdeltasmallboth.Gene(1:2:332),"Delta ExpressionDimorphismIndex Sorted");
end

for t=5:34
    for v=1:166
        m=datasetdeltasmallboth{2*v,t};
        f=datasetdeltasmallboth{2*v-1,t};
        ix=(abs(f)-abs(m))/(abs(f)+abs(m));
        dispar=abs(f-m);
        if(isnan(ix))
            ix=0;
        end
        xdmidx(v)=ix;
        dispidx(v)=dispar;
    end
%    titletxt=strcat("Delta ", datasetdeltasmallboth.Properties.VariableNames{t}," Dimorphism Index x Expression Dimorphism Index");
%    fprintf("%s\n",titletxt);
%    make2dplotnozeros(xdmidx, datasetdeltasmallboth.DimorphismIndex2(1:2:332), datasetdeltasmallboth.Gene(1:2:332), datasetdeltasmallboth.Sex(1:2:332), titletxt,strcat(datasetdeltasmallboth.Properties.VariableNames{t}," Dimorphism Index"), "Expression Index");

%    titletxt=strcat("Delta ", datasetdeltasmallboth.Properties.VariableNames{t}," Disparity x Expression Dimorphism Index");
%    fprintf("%s\n",titletxt);
%    make2dplotnozeros(dispidx, datasetdeltasmallboth.DimorphismIndex2(1:2:332), datasetabsolutesmallboth.Gene(1:2:332), datasetdeltasmallboth.Sex(1:2:332), titletxt, strcat(datasetdeltasmallboth.Properties.VariableNames{t}," Disparity Index"), "Expression Index");

%    titletxt=strcat("Delta ", datasetdeltasmallboth.Properties.VariableNames{t}," Disparity x Dimorphism Index");
%    fprintf("%s\n",titletxt);
%    make2dplotnozeros(dispidx, xdmidx, datasetdeltasmallboth.Gene(1:2:332), datasetdeltasmallboth.Sex(1:2:332), titletxt, strcat(datasetdeltasmallboth.Properties.VariableNames{t}," Disparity Index"), strcat(datasetdeltasmallboth.Properties.VariableNames{t}," Dimorphism Index"));
    
    pairnames=dataorig.Gene(1:2:332);
    fprintf("%s dimorphism plot\n",datasetdeltasmallboth.Properties.VariableNames{t});
    if(makeplots)
    %%%makesortedplot2(xdmidx,pairnames,strcat("Delta ",datasetdeltasmallboth.Properties.VariableNames{t}," DimorphismIndex Sorted"));
    %%%makesortedplot(dispidx,pairnames,strcat("Delta ",datasetdeltasmallboth.Properties.VariableNames{t}," Disparity Sorted"));
    end
    close all
end


phenotypedimorphismindex2=zeros(166,1);
pairnames=dataorig.Gene(1:2:332);
for t=1:genotypes
    idx=(fd2fws{absolute,small,both}(t)-md2mws{absolute,small,both}(t))/(fd2fws{absolute,small,both}(t)+md2mws{absolute,small,both}(t));
    if(isnan(idx))
        idx=0;
    end
    phenotypedimorphismindex(t)=idx;
    datasetabsolutesmallboth.PhenotypeDimorphismIndex(2*t-1)=idx;
    datasetabsolutesmallboth.PhenotypeDimorphismIndex(2*t)=idx;
end
if(makeplots)
    %%%makesortedplot2(phenotypedimorphismindex,pairnames,"Abs PhenotypeDimorphismIndex -- PCA, Sorted");
%%%makesortedplot(datasetabsolutesmallboth.FMDist(1:2:332),pairnames,"Abs PhenotypeDisparity -- PCA, Sorted");
%%%makesortedplot2(datasetabsolutesmallboth.DimorphismIndex2(1:2:332),datasetabsolutesmallboth.Gene(1:2:332),"Abs ExpressionDimorphismIndex Sorted");
end

%for t=5:34
%    for v=1:166
%        m=datasetabsolutesmallboth{2*v,t};
%        f=datasetabsolutesmallboth{2*v-1,t};
%        ix=(abs(f)-abs(m))/(abs(f)+abs(m));
%        dispar=abs(f-m);
%        if(isnan(ix))
%            ix=0;
%        end
%        xdmidx(v)=ix;
%        dispidx(v)=dispar;
%    end
%    pairnames=dataorig.Gene(1:2:332);
%    fprintf("%s dimorphism plot\n",datasetabsolutesmallboth.Properties.VariableNames{t});
if(makeplots)
    %    makesortedplot(xdmidx,pairnames,strcat("Abs ",datasetabsolutesmallboth.Properties.VariableNames{t}," DimorphismIndex Sorted"));
%    makesortedplot(dispidx,pairnames,strcat("Abs ",datasetabsolutesmallboth.Properties.VariableNames{t}," Disparity Sorted"));
end
%    close all
%end


if(makeplots)
    %%%makesortedplot(datasetabsolutesmallboth.AstCEE(1:2:332),pairnames,"Expression in AstC EEs Sorted");
    %%%makesortedplot(datasetabsolutesmallboth.AstAEE(1:2:332),pairnames,"Expression in AstA EEs Sorted");
    %%%makesortedplot(datasetabsolutesmallboth.NPFEE(1:2:332),pairnames,"Expression in NPF EEs Sorted");
end

if(makeplots)

for nx=1:2
dataset=datasetabsolutesmallboth;
dname="Absolute";
    if nx==1
dataset=datasetdeltasmallboth;
dname="Delta";
    end
    figure
hold on
drawnow
xes=zeros(166,1);
for t=5:22
    name1=dataset.Properties.VariableNames{t};
    fdata=dataset{1:2:332,t};
    mdata=dataset{2:2:332,t};
    mf=max(fdata);
    mm=max(mdata);
    m=max(mf,mm);
    fdata=fdata/m;
    mdata=mdata/m;
    xes(1:166)=t;
    plot(xes+0.1,fdata,'Marker','o','Color','red','MarkerSize',10, 'MarkerEdgeColor','red');
    plot(xes-0.1,mdata,'Marker','o','Color','blue','MarkerSize',10, 'MarkerEdgeColor','blue');
    plot([t-0.4 t+0.4],[sum(mdata)/166 sum(mdata)/166],'LineWidth',3,'Color','blue');
    plot([t-0.4 t+0.4],[sum(fdata)/166 sum(fdata)/166],'LineWidth',3,'Color','red'); %,'Marker','_','Color','red','MarkerSize',30, 'MarkerEdgeColor','red');
    for v=1:166
        for nn=1:size(specialgenes,1)
            if dataset.Gene{v*2}==specialgenes{nn,1}
                text(t-0.2, mdata(v), specialgenes{nn,1}, "FontSize",5, 'Color', specialgenes{nn,3} ,'HorizontalAlignment','right');
                text(t+0.2, fdata(v), specialgenes{nn,1}, "FontSize",5, 'Color', specialgenes{nn,2});
            end
        end
    end
    [xttest1 xttest]=ttest2(mdata,fdata,'Vartype','unequal');
    pt="";
    text(t, 1.2, datasetdeltasmallboth.Properties.VariableNames{t},"FontSize",10, 'Rotation',45); %HorizontalAlignment', 'center');
    if(xttest<0.05)
        pt="*";
        if(xttest<0.01)
            pt="**";
            if(xttest<0.001)
                pt="***";
            end
        end
        text(t, 1.1, pt,"FontSize",20, 'HorizontalAlignment', 'center');
    end
    drawnow
    yline(0);
end
ff=gcf;
ff.Units='normalized';
ff.Position=[0 0 1 1];
ff.Children.YLim=[ff.Children.YLim(1) 1.5];
tx=strcat(dname, " phenotypes M vs F with t-tests");
title(tx);
saveas(gcf,strcat("PNGs\",tx,".png"));
if(savepdfs)
    print(gcf,'-vector','-dpdf',strcat("PDFs\",tx,".pdf"));
end
close all
end

end


dataset=datasetdeltasmallboth;
for n=1:8
k=kmeans(scores{delta,small,both},n);
for t=1:332
    names{t}=strcat(dataset.Gene{t},"-",dataset.Sex{t});
end
%%%make2dplotnozeros2([scores{delta,small,both}(:,1)],[scores{delta,small,both}(:,2)], names, k, strcat("Delta ",num2str(n),"-Means Clustering PCA1 x PCA2"), "PCA1", "PCA2");
%%%make2dplotnozeros3([scores{delta,small,both}(:,1)],[scores{delta,small,both}(:,2)], names, k, strcat("Delta ",num2str(n),"-Means Clustering PCA1 x PCA2 Cleaner"), "PCA1", "PCA2");
end

dataset=datasetabsolutesmallboth;
for n=1:8
k=kmeans(scores{absolute,small,both},n);
for t=1:332
    names{t}=strcat(dataset.Gene{t},"-",dataset.Sex{t});
end
%%%make2dplotnozeros2([scores{absolute,small,both}(:,1)],[scores{absolute,small,both}(:,2)], names, k, strcat("Abs ",num2str(n),"-Means Clustering PCA1 x PCA2"), "PCA1", "PCA2");
%%%make2dplotnozeros3([scores{absolute,small,both}(:,1)],[scores{absolute,small,both}(:,2)], names, k, strcat("Abs ",num2str(n),"-Means Clustering PCA1 x PCA2 Cleaner"), "PCA1", "PCA2");
end

tgs={"Phm", "Pal2", "amon", "svr", "NPF", "AstC", "Orcokinin", "Tk"};
for l=0:1
    ss=" Male ";
    if l==1
        ss=" Female ";
    end
for k=1:8
tg=tgs{k};            
dataset=datasetabsolutesmallboth;
target=166;
for t=1:166
    if(dataset.Gene{t*2}==tg)
        target=t;
        break;
    end
end
dists=zeros(166,1);
hdists=zeros(166,1);
x1=scores{absolute,small,both}(2*target-l,1);
y1=scores{absolute,small,both}(2*target-l,2);
for t=1:166
x2=scores{absolute,small,both}(2*t-l,1);
y2=scores{absolute,small,both}(2*t-l,2);
dists(t)=sqrt((x2-x1)^2+(y2-y1)^2);
for d=1:18
    hdists(t)=hdists(t)+(scores{absolute,small,both}(2*target-l,d)-scores{absolute,small,both}(2*t-l,d))^2;
end
hdists(t)=sqrt(hdists(t));
end
if(makeplots)
    %%%makesortedplot(dists,dataset.Gene((2-l):2:332),strcat("Abs Dist of", ss, "RNAi to",ss,tg,"-i"));
%%%makesortedplot(hdists,dataset.Gene((2-l):2:332),strcat("Abs HDist of", ss, "RNAi to",ss,tg, "-i"));
end

dataset=datasetdeltasmallboth;
target=166;
for t=1:166
    if(dataset.Gene{t*2}==tg)
        target=t;
        break;
    end
end
dists=zeros(166,1);
hdists=zeros(166,1);
x1=scores{delta,small,both}(2*target-l,1);
y1=scores{delta,small,both}(2*target-l,2);
for t=1:166
x2=scores{delta,small,both}(2*t-l,1);
y2=scores{delta,small,both}(2*t-l,2);
dists(t)=sqrt((x2-x1)^2+(y2-y1)^2);
for d=1:18
    hdists(t)=hdists(t)+(scores{delta,small,both}(2*target-l,d)-scores{delta,small,both}(2*t-l,d))^2;
end
hdists(t)=sqrt(hdists(t));
end
if(makeplots)
    %%%makesortedplot(dists,dataset.Gene((2-l):2:332),strcat("Delta Dist of",ss,"RNAi to",ss,tg,"-i"));
%%%makesortedplot(hdists,dataset.Gene((2-l):2:332),strcat("Delta HDist of",ss,"RNAi to",ss,tg,"-i"));
end
end
end


if(makeplots)
dataset=datasetabsolutesmallboth;
dname="Abs";
for t=5:22
    %%%makesortedplot([dataset{1:2:332,t}],dataset.Gene(1:2:332),strcat(dname," ", dataset.Properties.VariableNames{t}, " Females Sorted"));
    %%%makesortedplot([dataset{2:2:332,t}],dataset.Gene(2:2:332),strcat(dname," ", dataset.Properties.VariableNames{t}, " Males Sorted"));
    %%%make2dplotnozeros( [dataset{1:2:332,t}], [dataset{2:2:332,t}], {dataset.Gene{2:2:332}}, {dataset.Sex{2:2:332}}, strcat("Abs ",dataset.Properties.VariableNames{t}, " F x M"), strcat("Female ",dataset.Properties.VariableNames{t}), strcat("Male ",dataset.Properties.VariableNames{t}));
    make2dplotnozeros4( [dataset{1:2:332,t}], [dataset{2:2:332,t}], {dataset.Gene{2:2:332}}, {dataset.Sex{2:2:332}}, strcat("Abs ",dataset.Properties.VariableNames{t}, " F x M, Cleaner"), strcat("Female ",dataset.Properties.VariableNames{t}), strcat("Male ",dataset.Properties.VariableNames{t}));
    %        make2dplotnozeros([datasetdeltasmallboth{:,t}],   [datasetdeltasmallboth{:,v}],     datasetdeltasmallboth.Gene,   datasetdeltasmallboth.Sex, titletxt );

end

dataset=datasetdeltasmallboth;
dname="Delta";
for t=5:22
    %%%makesortedplot([dataset{1:2:332,t}],dataset.Gene(1:2:332),strcat(dname," ", dataset.Properties.VariableNames{t}, " Females Sorted"));
    %%%makesortedplot([dataset{2:2:332,t}],dataset.Gene(2:2:332),strcat(dname," ", dataset.Properties.VariableNames{t}, " Males Sorted"));
    %%%make2dplotnozeros(  [dataset{1:2:332,t}], ...
    %%%                    [dataset{2:2:332,t}], ...
    %%%                    {dataset.Gene{2:2:332}}, ...
    %%%                    {dataset.Sex{2:2:332}}, ...
    %%%                    strcat("Delta ",dataset.Properties.VariableNames{t}," F x M"), ...
    %%%                    strcat("Female ",dataset.Properties.VariableNames{t}), ...
    %%%                    strcat("Male ",dataset.Properties.VariableNames{t}));
    make2dplotnozeros4(  [dataset{1:2:332,t}], ...
                        [dataset{2:2:332,t}], ...
                        {dataset.Gene{2:2:332}}, ...
                        {dataset.Sex{2:2:332}}, ...
                        strcat("Delta ",dataset.Properties.VariableNames{t}," F x M, Cleaner"), ...
                        strcat("Female ",dataset.Properties.VariableNames{t}), ...
                        strcat("Male ",dataset.Properties.VariableNames{t}));
end

%SDFGSDFGSDF
make2dplotnozeros4(  [dataset.FDist2Mw(1:2:332)], ...
                        [dataset.MDist2Fw(2:2:332)], ...
                        {dataset.Gene{2:2:332}}, ...
                        {dataset.Sex{2:2:332}}, ...
                        strcat("Delta Dist to Ctrl F x M, Cleaner"), ...
                        strcat("Female Dist"), ...
                        strcat("Male Dist"));
end

if(makeplots)
    dataset=datasetabsolutesmallboth;
k=kmeans(scores{absolute,small,both},4);
for t=1:332
    names{t}=strcat(dataset.Gene{t},"-",dataset.Sex{t});
end
%%%make2dplotnozeros2([scores{absolute,small,both}(:,1)],[scores{absolute,small,both}(:,3)], names, k, strcat("Abs 4-Means Clustering PCA1 x PCA3"), "PCA1", "PCA3");
%%%make2dplotnozeros2([scores{absolute,small,both}(:,1)],[scores{absolute,small,both}(:,2)], names, k, strcat("Abs 4-Means Clustering PCA1 x PCA2"), "PCA1", "PCA2");
%%%make2dplotnozeros2([scores{absolute,small,both}(:,2)],[scores{absolute,small,both}(:,3)], names, k, strcat("Abs 4-Means Clustering PCA2 x PCA3"), "PCA2", "PCA3");
% make2dplotnozeros2([scores{absolute,small,both}(:,1)],[scores{absolute,small,both}(:,4)], names, k, strcat("Abs 4-Means Clustering PCA1 x PCA4"), "PCA1", "PCA4");
% make2dplotnozeros2([scores{absolute,small,both}(:,2)],[scores{absolute,small,both}(:,4)], names, k, strcat("Abs 4-Means Clustering PCA2 x PCA4"), "PCA2", "PCA4");
% make2dplotnozeros2([scores{absolute,small,both}(:,3)],[scores{absolute,small,both}(:,4)], names, k, strcat("Abs 4-Means Clustering PCA3 x PCA4"), "PCA3", "PCA4");
% make2dplotnozeros3([scores{absolute,small,both}(:,1)],[scores{absolute,small,both}(:,3)], names, k, strcat("Abs 4-Means Clustering PCA1 x PCA3 Cleaner"), "PCA1", "PCA3");
% make2dplotnozeros3([scores{absolute,small,both}(:,1)],[scores{absolute,small,both}(:,2)], names, k, strcat("Abs 4-Means Clustering PCA1 x PCA2 Cleaner"), "PCA1", "PCA2");
% make2dplotnozeros3([scores{absolute,small,both}(:,2)],[scores{absolute,small,both}(:,3)], names, k, strcat("Abs 4-Means Clustering PCA2 x PCA3 Cleaner"), "PCA2", "PCA3");
% make2dplotnozeros3([scores{absolute,small,both}(:,1)],[scores{absolute,small,both}(:,4)], names, k, strcat("Abs 4-Means Clustering PCA1 x PCA4 Cleaner"), "PCA1", "PCA4");
% make2dplotnozeros3([scores{absolute,small,both}(:,2)],[scores{absolute,small,both}(:,4)], names, k, strcat("Abs 4-Means Clustering PCA2 x PCA4 Cleaner"), "PCA2", "PCA4");
% make2dplotnozeros3([scores{absolute,small,both}(:,3)],[scores{absolute,small,both}(:,4)], names, k, strcat("Abs 4-Means Clustering PCA3 x PCA4 Cleaner"), "PCA3", "PCA4");

%make3dplot([scores{absolute,small,both}(:,1)],[scores{absolute,small,both}(:,2)],[scores{absolute,small,both}(:,3)], names, k, "Abs 4-Means Clustering", "PCA1", "PCA2", "PCA3");
end

if(~makeallcombos)
    return
end


global malelm;
global femalelm;

for t=5:24
%    for v=(t+1):24
    for v=5:24
        titletxt=strcat("Delta ",datasetdeltasmallboth.Properties.VariableNames{t}," x ",datasetdeltasmallboth.Properties.VariableNames{v});
        fprintf("%f %f %s\n",t,v,titletxt);
        %make2dplotnozeros([datasetdeltasmallboth{:,t}],   [datasetdeltasmallboth{:,v}],     datasetdeltasmallboth.Gene,   datasetdeltasmallboth.Sex, titletxt, datasetdeltasmallboth.Properties.VariableNames{t},datasetdeltasmallboth.Properties.VariableNames{v}  );
        titletxt=strcat("Delta ",datasetdeltasmallboth.Properties.VariableNames{t}," x ",datasetdeltasmallboth.Properties.VariableNames{v},", Cleaner");
        make2dplotnozeros5([datasetdeltasmallboth{:,t}],   [datasetdeltasmallboth{:,v}],     datasetdeltasmallboth.Gene,   datasetdeltasmallboth.Sex, titletxt, datasetdeltasmallboth.Properties.VariableNames{t},datasetdeltasmallboth.Properties.VariableNames{v}  );
        close all
    end
end
for t=5:24
%    for v=(t+1):24
    for v=5:24
        titletxt=strcat("Abs ",datasetabsolutesmallboth.Properties.VariableNames{t}," x ",datasetabsolutesmallboth.Properties.VariableNames{v});
        fprintf("%f %f %s\n",t,v,titletxt);
        %make2dplotnozeros([datasetabsolutesmallboth{:,t}],   [datasetabsolutesmallboth{:,v}],     datasetabsolutesmallboth.Gene,   datasetabsolutesmallboth.Sex, titletxt, datasetabsolutesmallboth.Properties.VariableNames{t}, datasetabsolutesmallboth.Properties.VariableNames{v});
        titletxt=strcat("Abs ",datasetabsolutesmallboth.Properties.VariableNames{t}," x ",datasetabsolutesmallboth.Properties.VariableNames{v},", Cleaner");
        make2dplotnozeros5([datasetabsolutesmallboth{:,t}],   [datasetabsolutesmallboth{:,v}],     datasetabsolutesmallboth.Gene,   datasetabsolutesmallboth.Sex, titletxt, datasetabsolutesmallboth.Properties.VariableNames{t}, datasetabsolutesmallboth.Properties.VariableNames{v});
        close all
    end
end

discordancedataset=datasetdeltasmallboth;
for t=5:22
    for j=1:166
        f=discordancedataset{j*2-1,t};
        m=discordancedataset{j*2,  t};

%        a=atan(f/m);
        a=(f-m)/(f+m);
        if isnan(a)
            a=0;
        end
        
        discordancedataset{j*2-1,t}=a;
        discordancedataset{j*2,t}=a;
    end
end

standardizeddata=zscore(discordancedataset{1:2:332,5:22});
%standardizeddata=discordancedataset{1:2:332,5:22};

[coeffsd,scoresd,latent,tsquared,explainedsd,mu] = pca(standardizeddata);
pairnames=dataset.Gene(1:2:332);
scale=120;
%if(makeplots)

%end
for fff=1:2
    figure('visible','on')
    hold on
    for t=1:166
        c1="#888888";
        c5=c1;
        s=10;
        m='.';
        lw=0.1;
        lw2=0.1;
        tc='#888888';
        fw='normal';
        for nn=1:size(specialgenes,1)
            if pairnames(t)==specialgenes{nn,1}
                c1=specialgenes{nn,2};
                c2=specialgenes{nn,3};
                s=15;
                lw=2;
                lw2=0.5;
                m='o';
                c5='#000000';
                tc='black';
                fw='bold';
                break
            end
        end
        x1=scoresd(t,1);
        y1=scoresd(t,2);
        plot(x1, y1, 'Marker','o','MarkerSize',s,'MarkerFaceColor',c1,'MarkerEdgeColor','black','LineWidth',lw2);
        if(fff==1)
            text(x1, y1, dataset.Gene{t},"FontSize", 8,'Rotation',45,'Color',tc,'FontWeight',fw,'HorizontalAlignment','center');
        end
        %    end
    end %first round


    %put special dots and names on top
    for t=1:166
        s=15;
        m='.';
        lw=0.1;
        tc='black';
        for nn=1:size(specialgenes,1)
            if pairnames(t)==specialgenes{nn,1}
                c1=specialgenes{nn,2};
                c2=specialgenes{nn,3};
                s=15;
                lw=2;
                lw2=0.5;
                x1=scoresd(t,1);
                y1=scoresd(t,2);
                %           if(makeplots)
                plot(x1, y1, 'Marker','o','MarkerSize',s,'MarkerFaceColor',c1,'MarkerEdgeColor','black','LineWidth',lw2);
                text(x1, y1, pairnames{t},"FontSize", 8,'Rotation',45,'Color',tc,'FontWeight','bold','HorizontalAlignment','center');
                %            end
                break
            end
        end
    end
    %if(makeplots)
    ff=gcf;
    ff.Units='normalized';
    ff.Position=[0 0 1 1];
    tx=strcat(fname," ",suffix," Discordance PCA Cleaner");
    if(fff==1)
        tx=strcat(fname," ",suffix," Discordance PCA");
    end
    title(tx);
    saveas(gcf,strcat("PNGs\",tx,".png"));
    if(savepdfs)
        print(gcf,'-vector','-dpdf',strcat("PDFs\",tx,".pdf"));
    end
    %end
    close all
end
```
