## Supplemental code 3 for "Gut hormone signaling drives sex differences in metabolism and behavior": make2dplot.m

```
function make2dplot(xdata,ydata,names,sexes,titletext)
global specialgenes;
global fname;
global suffix;
global savepdfs;
global savepngs;
nummale=0;
numfemale=0;
malex=zeros(1,1);
femalex=zeros(1,1);
maley=zeros(1,1);
femaley=zeros(1,1);
figure('visible','off')
hold on

for t=1:length(xdata)
    found=0;
    for n=1:length(specialgenes)
        if(names{t}==specialgenes{n})
            plot(xdata(t), ydata(t), 'Marker','.','MarkerSize',30,'Color',specialgenes{n,2});
            text(xdata(t), ydata(t), strcat(names{t},"-",sexes{t}),"FontSize", 12,'Rotation',45, 'Color',specialgenes{n,2});
            found=1;
            break
        end
    end
    if(sexes{t}=="M")
        cc='blue';
        nummale=nummale+1;
        malex(nummale)=xdata(t);
        maley(nummale)=ydata(t);
    else
        cc='red';
        numfemale=numfemale+1;
        femalex(numfemale)=xdata(t);
        femaley(numfemale)=ydata(t);
    end
    if found==0
        plot(xdata(t), ydata(t), 'Marker','.','MarkerSize',20,'Color',cc);
        text(xdata(t),ydata(t),strcat(names{t},"-",sexes{t}),"FontSize", 6,'Rotation',45);
    end
end
[malefit,malefitS]=polyfit(malex, maley, 1);
[femalefit,femalefitS]=polyfit(femalex, femaley, 1);
maleslope=malefit(1);
maleb=malefit(2);
femaleslope=femalefit(1);
femaleb=femalefit(2);
malelinex=[min(malex) max(malex)];
femalelinex=[min(femalex) max(femalex)];
maleliney=maleslope*malelinex+maleb;
femaleliney=femaleslope*femalelinex+femaleb;
plot(malelinex,maleliney,'LineWidth',4,'Color','blue');
plot(femalelinex,femaleliney,'LineWidth',4,'Color','red');
r2male=(malefitS.normr/(norm(maley-mean(maley))))^2;
if(isnan(r2male))
    r2male=1;
end
r2male=1-r2male;
r2female=(femalefitS.normr/(norm(femaley-mean(femaley))))^2;
if(isnan(r2female))
    r2female=1;
end
r2female=1-r2female;

title(strcat(titletext, " -- male slope=",num2str(malefit(1)), " (R2=",num2str(r2male),"); female slope=", num2str(femalefit(1)), " (R2=",num2str(r2female),")"));
ff=gcf;
ff.Units='normalized';
ff.Position=[0 0 1 1];
drawnow
if(savepngs)
    saveas(gcf,strcat("PNGs\",titletext,".png"));
end
if(savepdfs)
    print(gcf,'-vector','-dpdf',strcat("PDFs\",titletext,".pdf"));
end
end
```
