## Supplemental code 3 for "Gut hormone signaling drives sex differences in metabolism and behavior": make2dplotnozeros2.m

```
function make2dplotnozeros2(xdata,ydata,names,groups,titletext,xaxistext,yaxistext)
global specialgenes;
global fname;
global suffix;
global savepdfs;
global savepngs;
colors={[0 0.5 0.5] [1 0 0] [0 1 0] [0 0 1] [0.9 0.9 0] [1 0 1] [0 .9 .9] [1 0.5 0] [0.5 0 1] };
colornames={'Teal','Red','Green','Blue','Yellow','Magenta','Cyan','Orange','Indigo'};
nummale=0;
numfemale=0;
malex=zeros(1,1);
femalex=zeros(1,1);
maley=zeros(1,1);
femaley=zeros(1,1);
figure('visible','off')
hold on
tots=zeros(max(groups),1);
females=zeros(max(groups),1);

for t=1:length(xdata)
    found=0;
    for n=1:length(specialgenes)
        if(names{t}==specialgenes{n}) || (names{t}==strcat(specialgenes{n},"-M")) || (names{t}==strcat(specialgenes{n},"-F"))
            plot(xdata(t), ydata(t), 'Marker','.','MarkerSize',30,'Color',colors{mod(groups(t),8)+1});
            text(xdata(t), ydata(t), names{t}, "FontSize", 12,'Rotation',45, 'Color',specialgenes{n,2});
            found=1;
            break
        end
    end
    if((xdata(t)~=0 && ydata(t)~=0) || (found==1))
        if found==0
            plot(xdata(t), ydata(t), 'Marker','.','MarkerSize',20,'Color',colors{mod(groups(t),8)+1});
            text(xdata(t),ydata(t), names{t} ,"FontSize", 6,'Rotation',45);
        end
    end
    tots(groups(t))=tots(groups(t))+1;
    if(names{t}{1}(length(names{t}{1}))=='F')
        females(groups(t))=females(groups(t))+1;
    end
end
xlim2=xlim;
ylim2=ylim;
if( xlim2(1)<0 && xlim2(2)>0)
    xline(0);
end
if( ylim2(1)<0 && ylim2(2)>0)
    yline(0);
end
titletext2=titletext;
for(nn=1:max(groups))
titletext2=strcat(titletext2, " G",num2str(nn),"(",colornames{mod(nn,8)+1},"): ",num2str(females(nn)/tots(nn)*100),"% ");
end
titletext2=strcat(titletext2, "Female");
title(titletext2);
xlabel(xaxistext);
ylabel(yaxistext);
ff=gcf;
ff.Units='normalized';
ff.Position=[0 0 1 1];
drawnow
if(savepngs)
    saveas(gcf,strcat("PNGs\",titletext,".png"));
end
if(savepdfs)
    print(gcf,'-vector','-dpdf',strcat("PDFs\",titletext,".pdf"));
end
end
```
