## Supplemental code 3 for "Gut hormone signaling drives sex differences in metabolism and behavior": make2dplotnozeros4.m

```
function make2dplotnozeros4(xdata,ydata,names,sexes,titletext,xaxistext,yaxistext)
global specialgenes;
global fname;
global suffix;
global savepdfs;
global savepngs;

confint=1.96; %1.96=95%, 2.576=99%

nummale=0;
numfemale=0;
malex=zeros(1,1);
femalex=zeros(1,1);
maley=zeros(1,1);
femaley=zeros(1,1);
figure('visible','off')
hold on

for t=1:length(xdata)
    found=0;
    for n=1:length(specialgenes)
        if(names{t}==specialgenes{n})
            plot(xdata(t), ydata(t), 'Marker','o','MarkerSize',15,'MarkerFaceColor',specialgenes{n, 2},'MarkerEdgeColor','black','LineWidth',0.5);
            text(xdata(t), ydata(t), names{t}, "FontSize", 12,'Rotation',45, 'Color','black','FontWeight','bold','HorizontalAlignment','center');
            found=1;
            break
        end
    end
    if((xdata(t)~=0 && ydata(t)~=0) || (found==1))
        if(sexes{t}=="M")
            cc='#8888ff';
            nummale=nummale+1;
            malex(nummale)=xdata(t);
            maley(nummale)=ydata(t);
        else
            cc='#ff8888';
            numfemale=numfemale+1;
            femalex(numfemale)=xdata(t);
            femaley(numfemale)=ydata(t);
        end
        if found==0
            plot(xdata(t), ydata(t), 'Marker','o','MarkerSize',10,'MarkerFaceColor','#aaaaaa','MarkerEdgeColor','black','LineWidth',0.1);
        end
    end
end

for t=1:length(xdata)
    for n=1:length(specialgenes)
        if(names{t}==specialgenes{n})
            plot(xdata(t), ydata(t), 'Marker','o','MarkerSize',15,'MarkerFaceColor',specialgenes{n, 2},'MarkerEdgeColor','black','LineWidth',0.5);
            text(xdata(t), ydata(t), names{t}, "FontSize", 12,'Rotation',45, 'Color','black','FontWeight','bold','HorizontalAlignment','center');
            break
        end
    end
end

meandot=plot(mean(xdata), mean(ydata), 'Marker','o','MarkerSize',30,'MarkerFaceColor','green','MarkerEdgeColor','black','LineWidth',1);


[malefit,malefitS]=polyfit(malex, maley, 1);
[femalefit,femalefitS]=polyfit(femalex, femaley, 1);
maleslope=malefit(1);
maleb=malefit(2);
femaleslope=femalefit(1);
femaleb=femalefit(2);
malelined=(max(malex)-min(malex))/100;
malelinex=[min(malex) max(malex)];
femalelinex=[min(femalex) max(femalex)];
maleliney=maleslope*malelinex+maleb;
femaleliney=femaleslope*femalelinex+femaleb;
[xttest1 xttest]=ttest2(malex,femalex,'Vartype','unequal');
[yttest1 yttest]=ttest2(maley,femaley,'Vartype','unequal');

mline=plot(malelinex,maleliney,'LineWidth',4,'Color','black');
malexpoints=[min(malex):malelined:max(malex)];
[malefitfit,malefitdelta]=polyval(malefit, malexpoints, malefitS);
lx=length(malexpoints);
bx=zeros(2*lx);
by=zeros(2*lx);
for t=1:lx
    bx(t)=malexpoints(t);
    bx(2*lx-t+1)=malexpoints(t);
    by(t)=malefitfit(t)+confint*malefitdelta(t);
    by(2*lx-t+1)=malefitfit(t)-confint*malefitdelta(t);
end
c=[0.8 0.8 0.8];
mci=fill(bx,by,c,'EdgeColor','#999999','FaceAlpha',0.5);

for t=1:length(xdata)
    found=0;
    for n=1:length(specialgenes)
        if(names{t}==specialgenes{n})
            found=1;
            break
        end
    end
    if(found==0)
        disty=abs((maleslope*xdata(t)+maleb)-ydata(t));
        for jj=1:100
        xx=min(malex)+malelined*jj;
        if xx>xdata(t)
            break
        end
        end
        if disty>confint*malefitdelta(jj)
            text(xdata(t), ydata(t), names{t}, "FontSize", 12,'Rotation',45, 'Color','black','FontWeight','bold','HorizontalAlignment','center');
        end
    end
end


    xlim2=xlim;
    ylim2=ylim;
    xmin=xlim2(1);
    xmax=xlim2(2);
    ymin=ylim2(1);
    ymax=ylim2(2);
zline=plot([max(xmin,ymin) min(xmax,ymax)], [max(xmin,ymin) min(xmax,ymax)], ':','LineWidth',1,'Color','#aaaaaa');
uistack(mline,'bottom');
uistack(zline,'bottom');
uistack(meandot,'bottom');
uistack(mci,'bottom');
ttxt=titletext;
r2male=0;
r2female=0;
if(nummale>1)
    r2male=(malefitS.normr/(norm(maley-mean(maley))))^2;
    if(isnan(r2male))
        r2male=1;
    end
    r2male=1-r2male;
    ttxt=strcat(ttxt, " -- slope=",num2str(malefit(1)), " (R2=",num2str(r2male),")");
else
    ttxt=strcat(ttxt," (no fit)");
end

xlim2=xlim;
ylim2=ylim;
maxslope=(ylim2(2)-ylim2(1))/(xlim2(2)-xlim2(1));
if( xlim2(1)<0 && xlim2(2)>0)
    xline(0);
end
if( ylim2(1)<0 && ylim2(2)>0)
    yline(0);
end
ttxt2="";
if(r2male>0.1 || r2female>0.1)
    ttxt2="(Some r2)";
end
if(r2male>0.3 || r2female>0.3)
    ttxt2="(Decent r2)";
end
if(r2male>0.7 || r2female>0.7)
    ttxt2="(Good r2)";
end
sloperatio=abs(maleslope/femaleslope);
if(isnan(sloperatio))
    sloperatio=1;
end
if( ...
        (((sloperatio>1.5) || (sloperatio<0.667)) && r2male>0.1 && r2female>0.1) ...
        || (~isnan(maleslope) && (abs(maleslope) > maxslope/25) && r2male>0.1) ...
        || (~isnan(femaleslope) && (abs(femaleslope) > maxslope/25) && r2female>0.1 ) ...
        )
    ttxt2=strcat(ttxt2,"(Slopes)");
end
if(yttest<0.05 || xttest<0.05)  
    ttxt2=strcat(ttxt2,"(Centers)");
end
ttxt=strcat(ttxt,ttxt2);
title(ttxt);
xlabel(xaxistext);
ylabel(yaxistext);
ff=gcf;
ff.Units='normalized';
ff.Position=[0 0 1 1];
drawnow
if(savepngs)
    saveas(gcf,strcat("PNGs\",titletext,ttxt2,".png"));
end
if(savepdfs)
    print(gcf,'-vector','-dpdf',strcat("PDFs\",titletext,ttxt2,".pdf"));
end
end
```
