## Supplemental code 3 for "Gut hormone signaling drives sex differences in metabolism and behavior": make2dplotnozeros5.m

```
function make2dplotnozeros5(xdata,ydata,names,sexes,titletext,xaxistext,yaxistext)
global specialgenes;
global fname;
global suffix;
global savepdfs;
global savepngs;
global malelm;
global femalelm;
global malex
global femalex
global malefit
global femalefit
global femalefitS
global malefitS
global maley
global femaley

nummale=0;
numfemale=0;
malex=zeros(1,1);
femalex=zeros(1,1);
maley=zeros(1,1);

confint=1.96; %1.96=95%, 2.576=99%

femaley=zeros(1,1);
figure('visible','off')
hold on

for t=1:length(xdata)
    if(strcmp(sexes{t},'M'))
        cc='#8888ff';
        nummale=nummale+1;
        malex(nummale)=xdata(t);
        maley(nummale)=ydata(t);
    else
        cc='#ff8888';
        numfemale=numfemale+1;
        femalex(numfemale)=xdata(t);
        femaley(numfemale)=ydata(t);
    end
    plot(xdata(t), ydata(t), 'Marker','o','MarkerSize',10,'MarkerFaceColor',cc,'MarkerEdgeColor','black','LineWidth',0.1);
end


[malefit,malefitS]=polyfit(malex, maley, 1);
[femalefit,femalefitS]=polyfit(femalex, femaley, 1);
malelm=fitlm(malex,maley);
femalelm=fitlm(femalex,femaley);
maleslopep=table2array(malelm.Coefficients);
maleslopep=maleslopep(2,4);
femaleslopep=table2array(femalelm.Coefficients);
femaleslopep=femaleslopep(2,4);

maleslope=malefit(1);
maleb=malefit(2);
femaleslope=femalefit(1);
femaleb=femalefit(2);
malelined=(max(malex)-min(malex))/100;
malelinex=[min(malex):malelined:max(malex)];
femalelined=(max(femalex)-min(femalex))/100;
femalelinex=[min(femalex):femalelined:max(femalex)];
maleliney=maleslope*malelinex+maleb;
femaleliney=femaleslope*femalelinex+femaleb;
[xttest1 xttest]=ttest2(malex,femalex,'Vartype','unequal');
[yttest1 yttest]=ttest2(maley,femaley,'Vartype','unequal');

malexpoints=[min(malex):malelined:max(malex)];
femalexpoints=[min(femalex):femalelined:max(femalex)];
[malefitfit,malefitdelta]=polyval(malefit, malexpoints, malefitS);
[femalefitfit,femalefitdelta]=polyval(femalefit, femalexpoints, femalefitS);

mline=plot(malexpoints,malefitfit,'LineWidth',4,'Color','blue');

fline=plot(femalexpoints,femalefitfit,'LineWidth',4,'Color','red');
lx=length(malexpoints);
bx=zeros(2*lx);
by=zeros(2*lx);
for t=1:lx
    bx(t)=malexpoints(t);
    bx(2*lx-t+1)=malexpoints(t);
    by(t)=malefitfit(t)+confint*malefitdelta(t);
    by(2*lx-t+1)=malefitfit(t)-confint*malefitdelta(t);
end
c=[0.9 0.9 1];
mci=fill(bx,by,c,'EdgeColor','#9999ff','FaceAlpha',0.5);

lx=length(femalexpoints);
bx=zeros(2*lx);
by=zeros(2*lx);
for t=1:lx
    bx(t)=femalexpoints(t);
    bx(2*lx-t+1)=femalexpoints(t);
    by(t)=femalefitfit(t)+confint*femalefitdelta(t);
    by(2*lx-t+1)=femalefitfit(t)-confint*femalefitdelta(t);
end
c=[1 0.9 0.9];
fci=fill(bx,by,c,'EdgeColor','#ff9999','FaceAlpha',0.5);


for t=1:nummale
%    found=0;
%    for n=1:length(specialgenes)
%        if(names{t}==specialgenes{n})
%            found=1;
%            break
%        end
%    end
%    if(found==0)
        disty=abs((maleslope*malex(t)+maleb)-maley(t));
        for jj=1:100
        xx=min(malex)+malelined*jj;
        if xx>malex(t)
            break
        end
        end
        if disty>2*malefitdelta(jj)
            text(malex(t), maley(t), names{t}, "FontSize", 12,'Rotation',45, 'Color','black','FontWeight','bold','HorizontalAlignment','center');
        end
%    end
end


for t=1:numfemale
%    found=0;
%    for n=1:length(specialgenes)
%        if(names{t}==specialgenes{n})
%            found=1;
%            break
%        end
%    end
%    if(found==0)
        disty=abs((femaleslope*femalex(t)+femaleb)-femaley(t));
        for jj=1:100
        xx=min(femalex)+femalelined*jj;
        if xx>femalex(t)
            break
        end
        end
        if disty>2*femalefitdelta(jj)
            text(femalex(t), femaley(t), names{t}, "FontSize", 12,'Rotation',45, 'Color','black','FontWeight','bold','HorizontalAlignment','center');
        end
%    end
end


% 
% 
% 
% 
% global specialgenes;
% global fname;
% global suffix;
% global savepdfs;
% global savepngs;
% nummale=0;
% numfemale=0;
% malex=zeros(1,1);
% femalex=zeros(1,1);
% maley=zeros(1,1);
% femaley=zeros(1,1);
% figure('visible','off')
% hold on
% 
% for t=1:length(xdata)
%     sexes{t}
%     found=0;
%     for n=1:length(specialgenes)
%         if(names{t}==specialgenes{n})
%             %plot(xdata(t), ydata(t), 'Marker','o','MarkerSize',15,'MarkerFaceColor',specialgenes{n, 2},'MarkerEdgeColor','black','LineWidth',0.5);
%             %text(xdata(t), ydata(t), names{t}, "FontSize", 12,'Rotation',45, 'Color','black','FontWeight','bold','HorizontalAlignment','center');
%             found=1;
%             break
%         end
%     end
%     if((xdata(t)~=0 && ydata(t)~=0) || (found==1))
%         if(strcmp(sexes{t},'M'))
%             cc='#8888ff';
%             nummale=nummale+1;
%             malex(nummale)=xdata(t);
%             maley(nummale)=ydata(t);
%         else
%             cc='#ff8888';
%             numfemale=numfemale+1;
%             femalex(numfemale)=xdata(t);
%             femaley(numfemale)=ydata(t);
%         end
%         if found==0
%             plot(xdata(t), ydata(t), 'Marker','o','MarkerSize',10,'MarkerFaceColor',cc,'MarkerEdgeColor','black','LineWidth',0.1);
%         end
%     end
% end
% 
% for t=1:length(xdata)
%     for n=1:length(specialgenes)
%         if(names{t}==specialgenes{n})
%             if(strcmp(sexes{t},'M'))
%                 nummale=nummale+1;
%             end
%             if(strcmp(sexes{t},'F'))
%                 numfemale=numfemale+1;
%             end
%             plot(xdata(t), ydata(t), 'Marker','o','MarkerSize',15,'MarkerFaceColor',specialgenes{n, 2},'MarkerEdgeColor','black','LineWidth',0.5);
%             text(xdata(t), ydata(t), strcat(names{t},"-",sexes{t}), "FontSize", 12,'Rotation',45, 'Color','black','FontWeight','bold','HorizontalAlignment','center');
%             break
%         end
%     end
% end
% 
% [malefit,malefitS]=polyfit(malex, maley, 1);
% [femalefit,femalefitS]=polyfit(femalex, femaley, 1);
% 
% 
% maleslope=malefit(1);
% maleb=malefit(2);
% femaleslope=femalefit(1);
% femaleb=femalefit(2);
% malelinex=[min(malex) max(malex)];
% femalelinex=[min(femalex) max(femalex)];
% maleliney=maleslope*malelinex+maleb;
% femaleliney=femaleslope*femalelinex+femaleb;
% [xttest1 xttest]=ttest2(malex,femalex,'Vartype','unequal');
% [yttest1 yttest]=ttest2(maley,femaley,'Vartype','unequal');
% 
% malexpoints=[min(malex):max(malex)];
% femalexpoints=[min(femalex):max(femalex)];
% [malefitfit,malefitdelta]=polyval(malefit, malexpoints, malefitS);
% [femalefitfit,femalefitdelta]=polyval(femalefit, femalexpoints, femalefitS);
% 
% mline=plot(malexpoints,malefitfit,'LineWidth',4,'Color','blue');
% 
% fline=plot(femalexpoints,femalefitfit,'LineWidth',4,'Color','red');
% lx=length(malexpoints);
% bx=zeros(2*lx);
% by=zeros(2*lx);
% for t=1:lx
%     bx(t)=malexpoints(t);
%     bx(2*lx-t+1)=malexpoints(t);
%     by(t)=malefitfit(t)+2*malefitdelta(t);
%     by(2*lx-t+1)=malefitfit(t)-2*malefitdelta(t);
% end
% c=[0.9 0.9 1];
% mci=fill(bx,by,c,'EdgeColor','#9999ff');
% 
% lx=length(femalexpoints);
% bx=zeros(2*lx);
% by=zeros(2*lx);
% for t=1:lx
%     bx(t)=femalexpoints(t);
%     bx(2*lx-t+1)=femalexpoints(t);
%     by(t)=femalefitfit(t)+2*femalefitdelta(t);
%     by(2*lx-t+1)=femalefitfit(t)-2*femalefitdelta(t);
% end
% c=[1 0.9 0.9];
% fci=fill(bx,by,c,'EdgeColor','#ff9999');


%mline2=plot(malexpoints,malefitfit+2*malefitdelta,'LineWidth',2,'Color','blue');
%mline3=plot(malexpoints,malefitfit-2*malefitdelta,'LineWidth',2,'Color','blue');
%fline2=plot(femalexpoints,femalefitfit+2*femalefitdelta,'LineWidth',2,'Color','red');
%fline3=plot(femalexpoints,femalefitfit-2*femalefitdelta,'LineWidth',2,'Color','red');


uistack(mline,'bottom');
uistack(fline,'bottom');
uistack(mci,'bottom');
uistack(fci,'bottom');
%uistack(mline2,'bottom');
%uistack(fline2,'bottom');
%uistack(mline3,'bottom');
%uistack(fline3,'bottom');

ttxt=titletext;
r2male=0;
r2female=0;
if(nummale>1)
    r2male=(malefitS.normr/(norm(maley-mean(maley))))^2;
    if(isnan(r2male))
        r2male=1;
    end
    r2male=1-r2male;
    ttxt=strcat(ttxt, " -- male slope=",num2str(malefit(1)), " (p=",num2str(maleslopep),", R2=",num2str(r2male),")");
else
    ttxt=strcat(ttxt," (no male fit)");
end

if(numfemale>1)
    r2female=(femalefitS.normr/(norm(femaley-mean(femaley))))^2;
    if(isnan(r2female))
        r2female=1;
    end
    r2female=1-r2female;
    ttxt=strcat(ttxt," -- female slope=", num2str(femalefit(1)), " (p=",num2str(femaleslopep),", R2=",num2str(r2female),")");
else
    ttxt=strcat(ttxt," (no female fit)");
end
ttxt=strcat(ttxt," (x p=",num2str(xttest),"; y p=",num2str(yttest),")");
xlim2=xlim;
ylim2=ylim;
maxslope=(ylim2(2)-ylim2(1))/(xlim2(2)-xlim2(1));
if( xlim2(1)<0 && xlim2(2)>0)
    xline(0);
end
if( ylim2(1)<0 && ylim2(2)>0)
    yline(0);
end
ttxt2="";
if(r2male>0.1 || r2female>0.1)
    ttxt2="(Some r2)";
end
if(r2male>0.3 || r2female>0.3)
    ttxt2="(Decent r2)";
end
if(r2male>0.7 || r2female>0.7)
    ttxt2="(Good r2)";
end
sloperatio=abs(maleslope/femaleslope);
if(isnan(sloperatio))
    sloperatio=1;
end
if( ...
        (((sloperatio>1.5) || (sloperatio<0.667)) && r2male>0.1 && r2female>0.1) ...
        || (~isnan(maleslope) && (abs(maleslope) > maxslope/25) && r2male>0.1) ...
        || (~isnan(femaleslope) && (abs(femaleslope) > maxslope/25) && r2female>0.1 ) ...
        )
    ttxt2=strcat(ttxt2,"(Slopes)");
end
if(yttest<0.05 || xttest<0.05)  
    ttxt2=strcat(ttxt2,"(Centers)");
end
ttxt=strcat(ttxt,ttxt2);
title(ttxt);
xlabel(xaxistext);
ylabel(yaxistext);

ff=gcf;
ff.Units='normalized';
ff.Position=[0 0 1 1];
drawnow
strcat("PNGs\",titletext,ttxt2,".png")
if(savepngs)
    saveas(gcf,strcat("PNGs\",titletext,ttxt2,".png"));
end
if(savepdfs)
    print(gcf,'-vector','-dpdf',strcat("PDFs\",titletext,ttxt2,".pdf"));
end
gcf
end
```
