## Supplemental code 3 for "Gut hormone signaling drives sex differences in metabolism and behavior": makesortedplot2.m

```
function makesortedplot2(unsorteddata,unsortednames,titletext)
    global specialgenes;
    global fname;
    global suffix;
global savepdfs;
global savepngs;
[sorteddata,xx]=sort(unsorteddata);
    dy=(max(sorteddata)-min(sorteddata))/150;
    sortednames=unsortednames(xx);
    figure('visible','off')
    hold on
    plot(sorteddata, 'Marker','.','MarkerSize',10,'Color','black','LineWidth',1);
    for t=1:length(sorteddata)
        found=0;
        for n=1:length(specialgenes)
            if(sortednames{t}==specialgenes{n})
                plot(t, sorteddata(t), 'Marker','.','MarkerSize',30,'Color',specialgenes{n,2},'LineWidth',1);
                text(t, sorteddata(t)+dy, sortednames{t},"FontSize", 12,'Rotation',90, 'Color',specialgenes{n,3});
%                text(t, sorteddata(t)+dy, sortednames{t},"FontSize", 12,'Rotation',90, 'Color','black');
                found=1;
                break
            end
        end
        if found==0
            text(t,sorteddata(t)+dy,sortednames{t},"FontSize", 6,'Rotation',90);
        end
    end
    ff=gcf;
    ff.Units='normalized';
    ff.Position=[0 0 1 1];
    ff.Children.YLim=[-1 1];
    %tx=strcat(fname,suffix,titletext);
    title(titletext);
    xlabel("Genotype");
    ylabel("Value");
    xlim2=xlim;
    ylim2=ylim;
    xline(0);
    yline(0);
    drawnow
    if(savepngs)
        saveas(gcf,strcat("PNGs\",titletext,".png"));
    end
    if(savepdfs)
        print(gcf,'-vector','-dpdf',strcat("PDFs\",titletext,".pdf"));
    end
    close
end
```
