## Supplemental code 3 for "Gut hormone signaling drives sex differences in metabolism and behavior": pcachart.m

```
function pcachart(dataset,rows,cols)
global specialgenes;

d=zscore(dataset{rows,cols});
[coeffs,scores,latent,tsquared,explained,mu] = pca(d);

for t=1:size(d,1)
    c1="red";
    if(dataset.Sex{rows(t)}=="M")
        c1="blue";
    end
    s=20;
    m='.';
    lw=0.1;
    offset=0.03;
    tcolor='black';
    sz=8;
    for nn=1:size(specialgenes,1)
        if dataset.Gene(rows(t))==specialgenes{nn,1}
            c1=specialgenes{nn,2};
            s=50;
            lw=1;
            offset=0.06;
            sz=12;
            tcolor=c1;
            break
        end
    end
    x1=scores(t,1);
    y1=scores(t,2);
    plot(x1, y1, 'Marker',m,'MarkerSize',s,'Color',c1);
    hold on
    text(x1+offset, y1+offset, strcat(dataset.Gene{rows(t)},'-',dataset.Sex{rows(t)}),"FontSize", sz,'Rotation',45,'Color',tcolor);
end %first round

for t=1:size(d,1)
    found=0;
    for nn=1:size(specialgenes,1)
        if dataset.Gene(rows(t))==specialgenes{nn,1}
            x1=scores(t,1);
            y1=scores(t,2);
            c1=specialgenes{nn,2};
            plot(x1, y1, 'Marker',m,'MarkerSize',50,'Color',c1);
            hold on
            text(x1+0.06, y1+0.06, strcat(dataset.Gene{rows(t)},'-',dataset.Sex{rows(t)}),"FontSize", 12,'Rotation',45,'Color',c1);
            break
        end
    end
end
end
```
