## Supplemental code 3 for "Gut hormone signaling drives sex differences in metabolism and behavior": test1.m

```
function test(dataset,listn,sex,gene1,gene2)
    close all force
    range=5:22;
    rown=size(dataset,1);
    if(strcmp(listn,'allgenes'))
        if(strcmp(sex(1:4),'male'))
            list2=2:2:rown;
        end
        if(strcmp(sex(1:4),'fema'))
            list2=1:2:rown;
        end
        if(strcmp(sex(1:4),'both'))
            list2=1:rown;
        end
    end
    listx={};
    if(strcmp(listn,'peptides'))
%        listx={'cg13229','cg15614','cg31431','gr28b','gr36c','lkr','msr2','or49b','ppk17','phm','vglut2','cg7272','cg4991','cg16700','cg13248','cg13743','eaat2','amon','svr','pal2','asta','burs','dh44','dh31','gpb5','gpa2','npf','proc','crz','w1118','snpf','astc','trpa1','7b2','astcc','ccha1','ccha2','rya','cnma','dsk','ilp3','itp','lk','mip','ms','natalisin','nplp1','nplp3','nplp4','orcokinin','sifa','sut2','sut3','tk'};
        listx={'phm','vglut2','cg7272','cg4991','cg16700','cg13248','cg13743','eaat2','amon','svr','pal2','asta','burs','dh44','dh31','gpb5','gpa2','npf','proc','crz','w1118','snpf','astc','trpa1','7b2','astcc','ccha1','ccha2','rya','cnma','dsk','ilp3','itp','lk','mip','ms','natalisin','nplp1','nplp3','nplp4','orcokinin','sifa','sut2','sut3','tk'};
        list2=[];
        n=0;
        for t=1:length(listx)
            f=findgene2(dataset,strcat(listx{t},'m'));
            if f==-1
                fprintf("No such thing as %s\n",listx{t});
            else
                n=n+1;
                if(strcmp(sex,'both'))
                    list2(2*n-1)=findgene2(dataset,strcat(listx{t},'m'));
                    list2(2*n)=findgene2(dataset,strcat(listx{t},'f'));
                end
                if(strcmp(sex(1:4),'male'))
                    list2(n)=findgene2(dataset,strcat(listx{t},'m'));
                end
                if(strcmp(sex(1:4),'fema'))
                    list2(n)=findgene2(dataset,strcat(listx{t},'f'));
                end
            end
        end
    end

    %tree=cg2(dataset, list2, range, 'Spearman');
    %findintree2(tree,gene1,gene2);
    findcorrelation3(dataset,list2,range,gene1,gene2,listn,sex);
    %plot(tree);
    %name=strcat("Trees\",gene1," ",gene2," tree ",listn," ",sex,".pdf")
    %print(gcf,'-vector','-dpdf',name);
    %close(gcf)
    close all force
end
```
