## Supplemental code 3 for "Gut hormone signaling drives sex differences in metabolism and behavior": treetester.m

```
%d=datasetabsolutesmallboth;
d=datasetdeltasmallboth;
list={'phm','vglut2','cg7272','cg4991','cg16700','cg13248','cg13743','eaat2','amon','svr','pal2','asta','burs','dh44','dh31','gpb5','gpa2','npf','proc','crz','w1118','snpf','astc','trpa1','7b2','astcc','ccha1','ccha2','rya','cnma','dsk','ilp3','itp','lk','mip','ms','natalisin','nplp1','nplp3','nplp4','orcokinin','sifa','sut2','sut3','tk'};
list2=[];
n=0;
for t=1:length(list)
    f=findgene2(d,strcat(list{t},'m'));
    if f==-1
        fprintf("No such thing as %s\n",list{t});
    else
        n=n+1;
        list2(2*n-1)=findgene2(d,strcat(list{t},'m'));
        list2(2*n)=findgene2(d,strcat(list{t},'f'));
    end
end
tree=cg2(d, list2, 5:22, 'spearman');
findintree2(tree,'svr','astc');
```
