## Supplemental code 3 for "Gut hormone signaling drives sex differences in metabolism and behavior": untitled.m

```
try
    z=datasetdeltasmallboth.Name{1};
catch asdfasd
    load("datasets.mat");
end

ds2=datasetdeltasmallboth;

for x={'peptides' 'allgenes'}
    for s={'female' 'male'}
        for t={'phm','vglut2','cg7272','cg4991','cg16700','cg13248','cg13743','eaat2','amon','svr','pal2','asta','burs','dh44','dh31','gpb5','gpa2','npf','proc','crz','w1118','snpf','astc','trpa1','7b2','astcc','ccha1','ccha2','rya','cnma','dsk','ilp3','itp','lk','mip','ms','natalisin','nplp1','nplp3','nplp4','orcokinin','sifa','sut2','sut3','tk'}
            test1(ds2,x{1},s{1},t{1},'astc');
        end
    end
end
   
return


ds2{333,1}={'Avg'};
ds2{333,2}={'Avg'};
ds2{333,3}={'Avg'};
ds2{333,4}={'F'};
ds2{333,5:22}=0;
ds2{334,1}={'Avg'};
ds2{334,2}={'Avg'};
ds2{334,3}={'Avg'};
ds2{334,4}={'M'};
ds2{334,5:22}=0;
for t=5:22
    for r=1:2:332
        ds2{333,t}=ds2{333,t}+ds2{r,t}/166;
        ds2{334,t}=ds2{334,t}+ds2{r+1,t}/166;
    end
end

for x={'peptides' 'allgenes'}
    for s={'female' 'male'}
        test(ds2,x{1},s{1},'Avg','AstC');
    end
end
   
```
