## Supplemental code 3 for "Gut hormone signaling drives sex differences in metabolism and behavior": venn2.m

```
function bx=venn2(a,ab,b)
    ax=0;
    ay=0;
    ar=sqrt(a/3.141592);
    br=sqrt(b/3.141592);
    bx=ar+br;
    by=0;
    d=0;
    if(ab>b || ab>a || ab<0 || a<0 || b<0)
        fprintf("Venn2 bad values: A=%f    AB=%f    B=%f\n",a,ab,b);
        bx=-1;
        return
    end
    direction=(ar+br)/100;
    overlap=circleintersection(ax,ay,ar,bx,by,br);
    d=overlap-ab;
    dsign=abs(d)/d;
    if isnan(dsign)
        d=0;
        dsign=0;
    end
    count=0;
    while abs(d)>ab/100000000
        count=count+1;
        olddsign=dsign;
        bx=bx-direction;
        overlap=circleintersection(ax,ay,ar,bx,by,br);
        %circle(bx,by,br);
        %drawnow
        d=overlap-ab;
        dsign=abs(d)/d;
        if isnan(dsign)
            dsign=0;
        end
        if(dsign ~= olddsign)
            direction=direction*-0.5;
            %dsign=dsign*-1;
        end
        if(count>10000)
            fprintf("No venn2 fit\n");
            bx=-1;
            break
        end
    end
end
```
