## Supplemental code 3 for "Gut hormone signaling drives sex differences in metabolism and behavior": venn3.m

```
function th=venn3(txta,a,txtb,b,txtc,c,txtab,ab,txtac, ac,txtbc, bc,txtabc,abc)
    th=0;
    if(ab>a || ab>b ||ac>a || ac>c ||bc>b || bc>c ||abc>a||abc>b||abc>b)
        fprintf("Venn3 bad values: A=%f    B=%f    C=%f\n",a,b,c);
        fprintf("Venn3 bad values: AB=%f    AC=%f    BC=%f    ABC=%f\n",ab,ac,bc, abc);
        th=-10;
        return
    end
    d=venn2(a,ab,b);
    d2=venn2(b,bc,c);
    if( d==-1 || d2==-1)
        fprintf("No venn2 fit\n");
        th=-10;
        return;
    end
    ra=sqrt(a/3.141592);
    circle(0,0,ra);
    rb=sqrt(b/3.141592);
    circle(d,0,rb);
    direction=0.01;
    th=0;
    cx=d2*cos(th);
    cy=d2*sin(th);
    ac2=circleintersection(0,0,sqrt(a/3.141592),d+cx,cy,sqrt(c/3.141592));
    dd=ac2-ac;
    dsign=abs(dd)/dd;
    if isnan(dsign)
        dd=0;
        dsign=0;
    end
    th=0-direction;
    count=0;
    while(abs(dd)>ac/100000000)
        olddsign=dsign;
        count=count+1;
        th=th+direction;
        cx=d2*cos(th);
        cy=d2*sin(th);
        ac2=circleintersection(0,0,sqrt(a/3.141592),d+cx,cy,sqrt(c/3.141592));
        %circle(d+cx,cy,sqrt(c/3.141592));
        %drawnow
        dd=ac2-ac;
        dsign=abs(dd)/dd;
        if isnan(dsign)
            dsign=0;
        end
        if(dsign ~= olddsign)
            direction=direction*-0.5;
            %dsign=dsign*-1;
        end
        if(count>10000)
            fprintf("No fit\n");
            th=-10;
            break
        end
    end
    rc=sqrt(c/3.141592);
    circle(d+cx,cy,rc);
    ax=0;
    ay=0;
    bx=d;
    by=0;
    cx=d+cx;
    atobslope=(by-ay)/(bx-ax);
    atocslope=(cy-ay)/(cx-ax);
    btocslope=(cy-by)/(cx-bx);
    abangle=atan(by/bx);
    acangle=atan(cy/cx);
    acx1=cos(acangle)*ra;
    acy1=sin(acangle)*ra;
    acx2=cx+cos(acangle+3.141592)*rc;
    acy2=cy+sin(acangle+3.141592)*rc;
    text((acx1+acx2)/2,(acy1+acy2)/2,strcat(txtac," ",num2str(ac-abc)),'HorizontalAlignment','Center');

    abx1=cos(abangle)*ra;
    aby1=sin(abangle)*ra;
    abx2=bx+cos(abangle+3.141592)*rb;
    aby2=by+sin(abangle+3.141592)*rb;
    text((abx1+abx2)/2,(aby1+aby2)/2,strcat(txtab," ",num2str(ab-abc)),'HorizontalAlignment','Center');

    bcx1=bx+cos(th)*rb;
    bcy1=by+sin(th)*rb;
    bcx2=cx+cos(th+3.141592)*rc;
    bcy2=cy+sin(th+3.141592)*rc;
    text((bcx1+bcx2)/2,(bcy1+bcy2)/2,strcat(txtbc," ",num2str(bc-abc)),'HorizontalAlignment','Center');

    text( (abx1+abx2+acx1+acx2+bcx1+bcx2)/6,  (acy1+acy2+aby1+aby2+bcy1+bcy2)/6,   strcat(txtabc," ",num2str(abc)),  'HorizontalAlignment','Center');
    
    text(ax+ra*cos(3.1415+(abangle+acangle)/2),ay+ra*sin(3.1415+(abangle+acangle)/2),strcat(txta," ", num2str(a-ab-ac-abc)),'HorizontalAlignment','Center');
    text(bx+rb*cos(1.5*3.1415+(th+acangle)/2),by+rb*sin(1.5*3.1415+(th+acangle)/2),strcat(txtb," ",num2str(b-ab-bc-abc)),'HorizontalAlignment','Center');
    text(cx+rb*cos((th+acangle)/2),cy+rc*sin((th+acangle)/2),strcat(txtc," ",num2str(c-bc-ac-abc)),'HorizontalAlignment','Center');
            saveas(gcf,strcat("PNGs\",txtabc,".png"));
            %if(savepdfs)
                print(gcf,'-vector','-dpdf',strcat("PDFs\",txtabc,".pdf"));
%            end

end
```
